# Spatiotemporal Single-Cell Analysis Reveals T Cell Clonal Dynamics and Phenotypic Plasticity in Human Graft-versus-Host Disease

**DOI:** 10.1101/2025.05.24.655962

**Authors:** Lingting Shi, Ajna Uzuni, Ximi K. Wang, Michael Pressler, David W. Harle, Shami Chakrabarti, Rodney Macedo, Kirubel Belay, Christian A. Gordillo, Erik Raps, Jia Yi (Ady) Zhang, Achille Nazaret, Joy L. Fan, Yinuo Jin, Xumin Shen, Joshua S. Fuller, Tamjeed Azad, Jessie Huang, Pranik Chainani, Julian A. Abrams, Armando Del Portillo, Markus Y. Mapara, Mohamed Alhamar, Megan Sykes, José L. McFaline-Figueroa, Elham Azizi, Ran Reshef

**Affiliations:** Irving Institute for Cancer Dynamics, Columbia University, New York, NY, 10027, USA; Department of Biomedical Engineering, Columbia University, New York, NY, 10027, USA; Columbia Center for Translational Immunology, Columbia University Irving Medical Center, New York, NY, 10032, USA; Herbert Irving Comprehensive Cancer Center, Columbia University, New York, NY, 10032, USA; Columbia Institute of Human Nutrition, Columbia University, New York, NY, 10032, USA; Department of Computer Science, Columbia University, New York, NY, 10027, USA; Department Statistics and Data Science, Yale University, CT, 06520, USA; Division of Digestive and Liver Diseases, Columbia University Irving Medical Center, Columbia University, New York, NY, 10032, USA; Department of Pathology and Cell Biology, Columbia University Irving Medical Center, Columbia University, New York, NY, 10032, USA; Department of Pathology and Laboratory Medicine, Henry Ford Health, Detroit, MI, 48202, USA; Data Science Institute, Columbia University, New York, NY, 10027, USA

**Keywords:** Graft-versus-host disease, allogeneic hematopoietic cell transplantation, alloreactive T cells, single-cell genomics, spatial transcriptomics, phenotypic plasticity, temporal dynamics, probabilistic model

## Abstract

Allogeneic hematopoietic cell transplantation (alloHCT) is curative for various hematologic diseases but often leads to acute graft-versus-host disease (GVHD), a potentially life-threatening complication. We leverage GVHD as a uniquely tractable disease model to dissect complex T-cell–mediated pathology in 27 alloHCT recipients. We integrate pre-transplant identification of alloreactive T-cells with longitudinal tracking across blood and gut, using mixed lymphocyte reaction-based clonal “fingerprinting”, TCR clonotyping, single-cell RNA/TCR sequencing, and spatial transcriptomics. Using DecompTCR, a novel computational tool for longitudinal TCR analysis, we uncover clonal expansion programs linked to GVHD severity and TCR features. Multi-omics profiling of gut biopsies reveals enrichment and clonal expansion of CD8⁺ effector and ZNF683(Hobit)⁺ resident memory T-cells, cytolytic remodeling of regulatory and unconventional T-cells, and localization of CD8⁺ effector T-cells near intestinal stem cells in crypt loss regions. This framework defines dynamic immune circuit rewiring and phenotypic plasticity with implications for biomarkers and therapies.

**Graphical Abstract:** 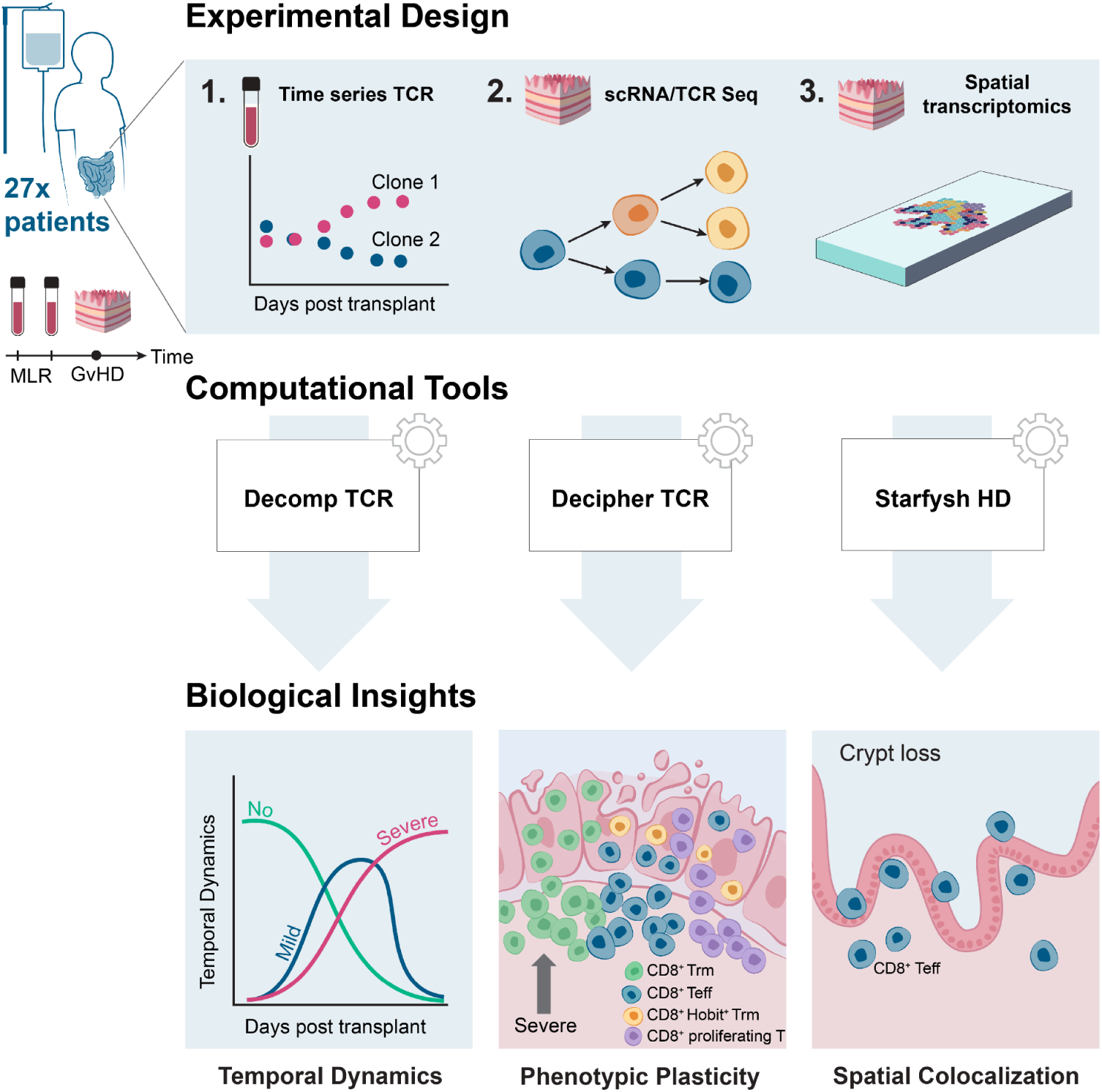

**Highlights:**

- Persistent expansion of diverse alloreactive T cell clones is a hallmark of severe GVHD
- DecompTCR reveals dynamic clonal expansion programs linked to GVHD severity and clinical outcome
- CD8+ T cell clones exhibit phenotypic plasticity *in vivo* across intestinal tissue compartments in GVHD
- High-resolution spatial profiling shows CD8+ effector T cells localize near intestinal stem cell niches and drive epithelial injury in GVHD

## INTRODUCTION

Allogeneic hematopoietic cell transplantation (alloHCT) is a curative therapy for various malignant and nonmalignant hematological diseases, including leukemia, myelodysplastic syndrome, and sickle cell disease. Despite significant progress in prophylactic and treatment strategies, acute graft-versus-host disease (GVHD) remains a major barrier to the success of HCT^1,2^, with incidence rates ranging from 30% to 70%^3^. More than 50% of patients who develop severe GVHD die within the first year after the transplant^4^.

Acute GVHD is mediated by donor-derived T lymphocytes that recognize recipient antigens, resulting in the activation of CD4^+^ helper and CD8^+^ cytotoxic T cells and leading to the subsequent destruction of host (recipient) tissue^5^. These alloreactive T cells specifically recognize and respond to mismatched major or minor histocompatibility antigens on host cells. While extensive studies using animal models have yielded critical insights into GVHD pathogenesis and informed therapeutic development^5–10^, a molecular and cellular characterization of GVHD in humans is still limited. In particular, there remains a critical need to prospectively identify clinically-relevant alloreactive clones, longitudinally track their dynamics *in vivo*, and precisely map their phenotypic states and interactions within affected tissues in patients, to guide biomarker discovery and targeted interventions.

GVHD also presents a uniquely tractable model for studying human T cell-mediated tissue pathology: the immune response is prospectively identifiable and shaped by therapeutic interventions, and the clinical course is closely monitored. As such, GVHD provides a rare opportunity to dissect *in vivo* T cell dynamics, including clonal expansion, phenotypic plasticity, tissue residency, and resistance to immunosuppression, in a well-defined human context. These insights thus have relevance not only to HCT but also to other diseases driven by dysregulated T cell responses, including intestinal transplantation, cancer, autoimmunity, and chronic infection.

To achieve a comprehensive view of human GVHD, we developed and applied an integrated multimodal framework to dissect the clonal, transcriptional and spatial dynamics of the T cell response (**Fig. 1A**). We first defined a “fingerprint” of the donor-derived alloreactive T cell response by combining mixed lymphocyte reactions (MLR) with high-throughput T cell receptor (TCR) sequencing targeting the third complementarity-determining region (CDR3) of the TCRβ chain (a key region of antigen specificity). This strategy has been previously employed in tolerance studies in solid organ or combined marrow/organ transplantation^11–14^, but applied here for the first time to standard human alloHCT. Using this approach, we then longitudinally tracked alloreactive T-cell clones in 20 alloHCT recipients for up to 2 years, uncovering distinct temporal patterns associated with GVHD^11^.

**Figure 1.**
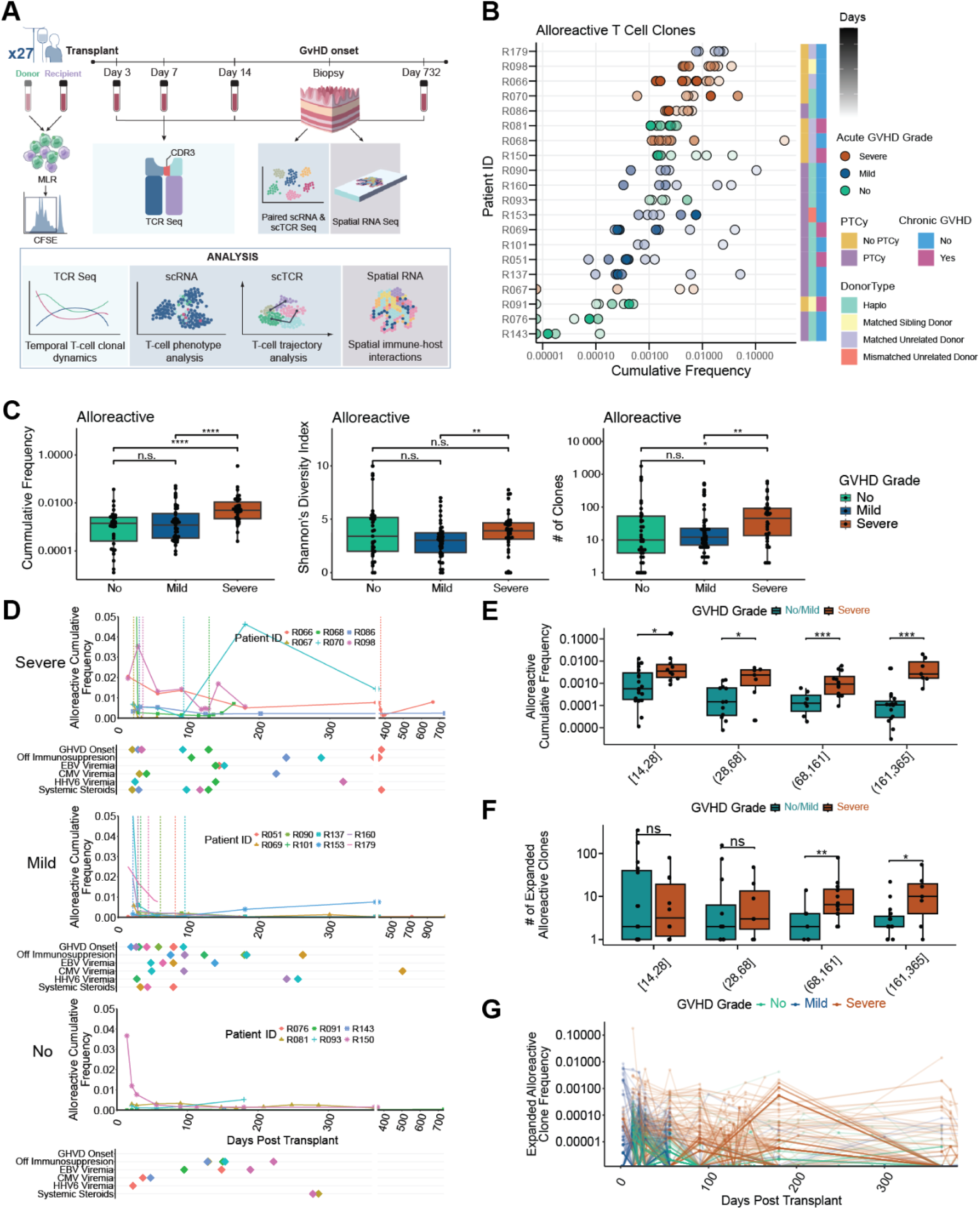
Severe GVHD is Associated with Expansion and Increased Diversity of Alloreactive Clones in the Blood. **A)** Schematic representation of the dataset and the analytical workflow. The diagram provides an overview of the data acquisition and downstream analyses conducted. **B)** Overview of the time-series TCR repertoire dataset. Patients were ordered vertically based on median cumulative alloreactive T cell frequency. Each dot represents a PBMC sample, color-coded by the patient’s acute GVHD grade, with darker shades denoting samples collected at later time points. **C)** Alloreactive T cell frequencies, diversity, and unique clone counts in patients grouped by GVHD grade (Benjamini-Hochberg adjusted Wilcoxon rank-sum test). **D)** Cumulative alloreactive frequency over time in patients with severe (top), mild (middle), and no (bottom) GVHD. The vertical color-coded dotted lines indicate GVHD onset. Other key clinical events such as cessation of immunosuppression, steroid use, and viral reactivations (EBV, CMV, HHV6) are shown below the respective plots. **E, F)** Cumulative frequency and number of expanded alloreactive clones over time (Benjamini-Hochberg adjusted Wilcoxon rank-sum test, * P < 0.05, ** P < 0.01, *** P < 0.001, **** P < 0.0001). **G)** Frequency over time of expanded alloreactive clones in patients with severe, mild, and no GVHD (color-coded).

Existing techniques to analyze TCR repertoire data^15–22^ face limitations in resolving clonal expansion dynamics over time and specifically separating alloreactive from non-alloreactive clones within the complex post-transplant environment. We thus introduce DecompTCR, a novel computational framework that uncovers latent clonal expansion programs^23^, leading to the identification of persistent expansion of subsets of alloreactive T cell clones in severe GVHD. Furthermore, we performed integrated single-cell RNA and TCR sequencing on patient gut biopsies, adapting our Decipher^23^ tool into DecipherTCR, to trace phenotypic transitions of individual T-cell clones across tissue compartments. Finally, we employed high-resolution spatial transcriptomics (ST) of patient gut biopsies and developed StarfyshHD (a VisiumHD adaptation of our spatial deconvolution and sample integration tool Starfysh^24^) to map the tissue microenvironment. Together, we revealed phenotypic T cell state transitions across tissue compartments and enrichment of cytotoxic CD8^+^ effector T cells in proximity to intestinal stem cells (ISCs) in severe GVHD.

Collectively, these integrated analyses constitute the first spatiotemporal and clonally-linked atlas of the human T cell response in GVHD. By combining prospective clonal tracking, time-resolved modeling, and tissue-level single-cell and spatial phenotyping, our study reveals fundamental principles of clonal persistence, phenotypic plasticity, and tissue remodeling in GVHD, offering new avenues for biomarker discovery and therapeutic targeting. Our findings from GVHD, a uniquely tractable disease model of human T cell-mediated pathology, illuminate principles of immune circuit dynamics and therapeutic resistance. Finally, our computational toolbox for multi-modal dissection of the temporal and spatial dynamics of T cell responses can be applied in diverse contexts including infection, autoimmune disease, and cancer.

## RESULTS

### Severe GVHD is marked by a persistently expanded and highly diverse alloreactive T cell repertoire

To identify and track human alloreactive T cells, we expanded donor-derived T cells that respond to recipient antigens through MLR of pre-transplant donor and recipient peripheral blood mononuclear cells (PBMC), collected from 20 donor-recipient pairs (**Supplementary Table 1,2,3**). Alloreactive T cell repertoires were identified using high-throughput sequencing of TCRβ VDJ recombination and CDR3 amino acid chains of the proliferating fraction of the MLR and an unstimulated donor sample as previously described^11^ (**Methods**). We conducted a time-series profiling of TCRβ CDR3 clonal repertoires in PBMCs from transplant recipients, spanning from day +3 to day +732 post-transplant (median of 8.5 time points per patient, range 5-12). Patients were categorized by varying GVHD grades (grades 1-2 denoted as mild and 3-4 as severe)^25^ and by the use of post-transplant cyclophosphamide (PTCy) prophylaxis, given its known impact on T cell population diversity after transplant^26^. Our cohort encompasses both malignant and nonmalignant hematological disorders and a range of donor types, from HLA-matched to mismatched and haploidentical (**Supplementary Table 1**).

We identified a total of 51,411 unique alloreactive clones across 20 MLRs, ranging from 256 to 6,243 clones per donor-recipient pair (**Methods**). We first computed the cumulative frequency of these alloreactive T cells at each timepoint and ranked patients according to the median frequency across all timepoints (**Fig. 1B**). This analysis revealed that severe GVHD is associated with higher cumulative frequencies of alloreactive clones, a trend not observed for non-alloreactive clones (**Supplementary Fig. 1A; Methods**). No observable differences in clonal diversity (Shannon’s index) were seen in both alloreactive and non-alloreactive clones (**Supplementary Fig. 1B, C**). A strong correlation was observed between the cumulative alloreactive frequency and GVHD: Recipients exhibiting a frequency consistently below 0.001 for all time points remained free of GVHD (Fisher’s exact test, p = 0.01754). Conversely, a cumulative frequency exceeding 0.01 was associated with either acute or chronic GVHD, with the majority of these cases presenting with severe disease. We further studied the repertoire of alloreactive clones by quantifying the frequencies, diversity, and number of circulating unique alloreactive clones (**Fig. 1C; Supplementary Fig. 2**). Severe GVHD was distinguished by significantly higher cumulative alloreactive frequency and number of unique alloreactive clones detectable post-transplant compared to mild and no GVHD (p < 0.05), as well as greater alloreactive clonal diversity compared to mild GVHD (p < 0.01), indicating a broader and more diverse alloreactive T cell repertoire.

We next examined the timing of alloreactive clonal expansion post-transplant by analyzing the dynamics of each patient’s cumulative frequency of alloreactive clones over time (**Fig. 1D**). Severe GVHD patients exhibited persistently higher alloreactive frequencies throughout the first year post-transplant, with early peaks in the number of detectable circulating alloreactive clones, suggestive of heightened immune reactivity. In contrast, patients with mild/no GVHD showed a consistent decline in the cumulative frequency of alloreactive clones over time, indicative of immune tolerance. Stratifying by time intervals within the first year post-transplant confirmed higher frequencies in severe GVHD across all time intervals (p < 0.05, **Fig. 1E**). We also quantified unique expanded alloreactive clones, using patient-specific thresholds to account for sampling differences (**Methods**). Severe GVHD patients showed a greater number of expanded alloreactive clones, particularly at later time points (>68 days; **Fig. 1F**), and these clones showed higher clonal persistence (**Fig. 1G**). These findings suggest sustained antigenic stimulation and a failure to establish peripheral tolerance. CD4^+^ and CD8^+^ T cell populations shared similar trends in the overall cumulative frequencies, although the higher number and diversity of unique alloreactive clones in severe GVHD was driven mostly by CD8^+^ T cells (**Supplementary Fig. 3**). Together, these data characterize severe GVHD by a robust and persistent expansion of a highly diverse alloreactive T cell repertoire, potentially explaining prolonged antigen stimulation and impaired immune tolerance.

### Distinct temporal dynamics of alloreactive clonal expansion define GVHD severity

The variable timing of clonal expansion and diversification of alloreactive clones across GVHD grades (**Fig. 1G**) prompted us to systematically characterize their temporal dynamics. To this end, we developed DecompTCR, a probabilistic model that decomposes longitudinal data (specifically, the frequencies of each T cell clone) into a small set of representative patterns (basis functions) and assigns weights reflecting each clone’s similarity to these patterns. Adapted from a framework for analysis of gene expression dynamics over time^23^, DecompTCR identified distinct temporal patterns of maximal clonal expansion (**Fig. 2A**), enabling each T cell clone’s dynamics to be modeled as a weighted combination of these patterns even with sparse or irregular sampling, thereby reconstructing and interpolating their behavior over time (**Methods, Supplementary Fig. 4A**). For the example clone highlighted in Fig. 2A, the model assigned the highest weight to the green pattern (basis 3), as it most closely resembles the clone’s dynamics. Performance of the model was confirmed using simulated data (**Methods; Supplementary Fig. 4B-C).** Importantly, the output of DecompTCR can be used for unbiased stratification of clones according to their temporal dynamics. Specifically, alloreactive T-cell clones were grouped using hierarchical clustering based on their decomposition weights for downstream analysis. Interestingly, we found that different grades of GVHD were associated with distinct representative patterns (**Fig. 2B**), while no clinical information was used by the algorithm, and clones were characterized only based on their time-resolved frequencies. We found two temporal patterns enriched in no GVHD (Basis 4 and Basis 8, proportions 62% and 63%, respectively), marked by an early and steep decrease in frequency over time, indicating clonal contraction and immune tolerance (**Methods, Supplementary Fig. 5A, B**). In contrast, mild GVHD was enriched for a transient clonal expansion at early time points, followed by a decrease over time (Basis 1), matching the early onset of GVHD in these patients in our dataset and their general responsiveness to treatment. Severe GVHD exhibited greater complexity and enrichment in several distinct patterns (Bases 2, 7, and 9) that either expanded over time and remained elevated or contracted but at later time points. These differential dynamics highlight the association between the temporal patterns of alloreactive clones and grades of GVHD and allow us to distinguish clones that are more likely pathogenic.

**Figure 2.**
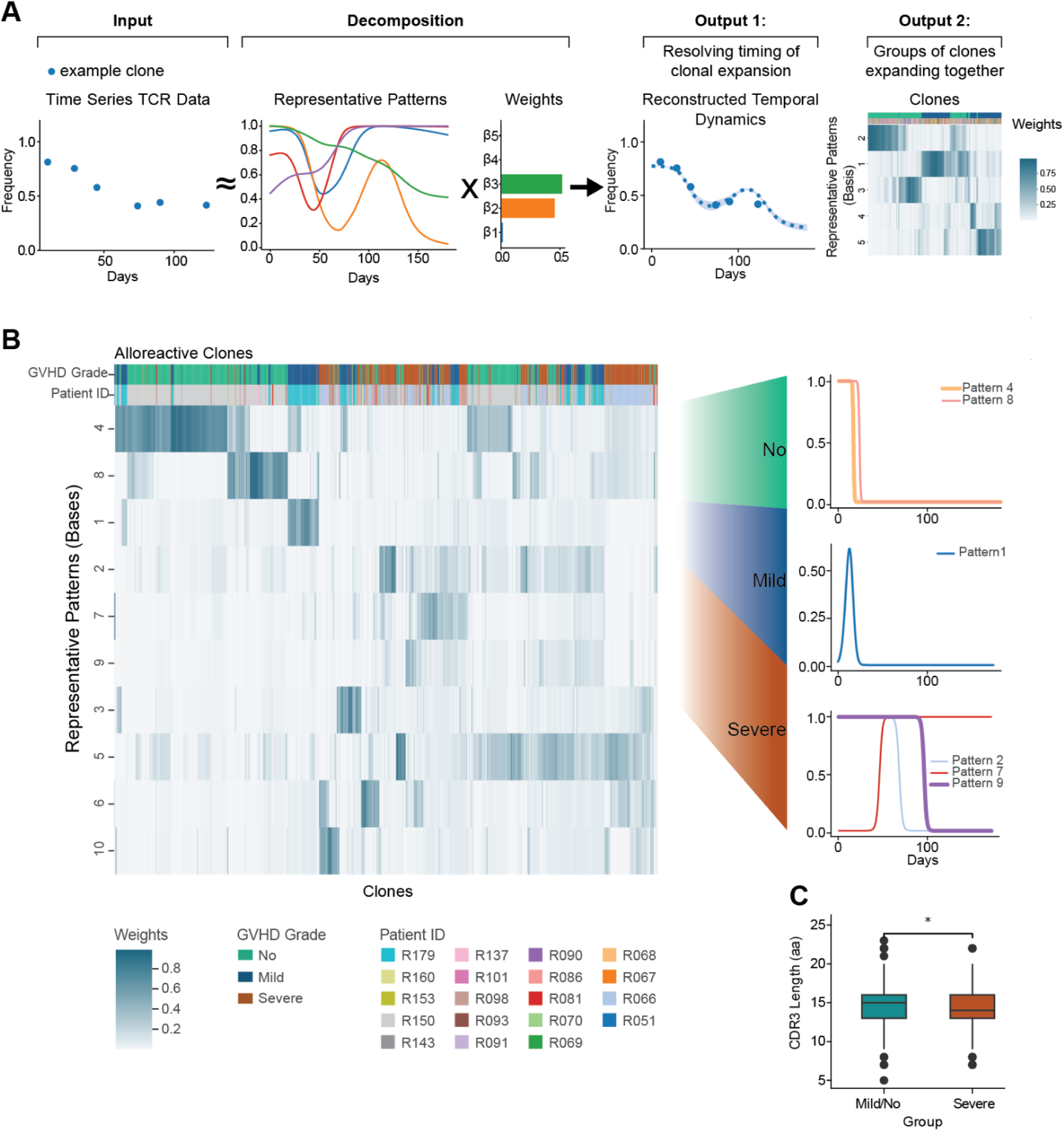
Time-Resolved Dynamic Modeling of T Cell Clones Demonstrates Associations between Alloreactive Clones and GVHD Severity. **(A)** Simulated data illustrating basis decomposition of individual T cell clones using DecompTCR. **(B)** Basis functions and corresponding weights derived from DecompTCR applied to alloreactive clones. The heatmap displays the weights of different bases (rows) with patient IDs and GVHD grades annotated across the columns. **(C)** Basis decomposition reveals shorter CDR3 amino acid length in T cell clones associated with severe GVHD-specific patterns (Wilcoxon rank-sum test with Holm adjustment, * P < 0.05, ** P < 0.01, *** P < 0.001, **** P < 0.0001).

The distinction between clinically relevant dynamic patterns within the alloreactive T cell repertoire motivated us to examine sequence-level features associated with pathogenicity. We found increased amino acid sequence similarity among alloreactive clones in mild and severe GVHD compared to no GVHD (**Supplementary Fig. 5C**), suggesting conserved CDR3 motifs among alloreactive clones that are truly pathogenic, irrespective of HLA specificity. Additionally, alloreactive clones associated with decompTCR from severe GVHD temporal patterns (bases 2, 7 & 9) have lower hydrophobicity in the CDR3 sequence compared to clones from no GVHD temporal patterns (bases 4 & 8), a difference that is not observed when comparing alloreactive clones from different clinical grades of GVHD alone without basis decomposition (**Supplementary Fig. 5D**). Lastly, alloreactive clones contained in severe GVHD patterns were characterized by shorter CDR3 sequences, suggesting that a skewed repertoire characterized by shorter and less hydrophobic CDR3 regions - features known to influence thymic selection and antigen affinity - may underpin the persistent expansion and pathogenicity of alloreactive clones in severe GVHD (**Fig. 2C**). These findings demonstrate the value of resolving disease grade-specific temporal patterns in T cell repertoire data to uncover features of pathogenic clones.

### PTCy selectively modulates alloreactive T cell dynamics

Given the distinct temporal patterns associated with GVHD severity, we next examined how post-transplant high-dose cyclophosphamide (PTCy) impacts alloreactive clones. PTCy is recognized as a highly effective strategy for preventing both severe acute and chronic GVHD^1^, thus its use in GVHD prophylaxis is increasing, especially among patients undergoing HCT with HLA-matched donors^27^. The model for the protective mechanism of PTCy has been described to eliminate proliferating T cells and overall decreases T cell diversity early post-transplant, and while the overall diversity of the TCR repertoire remains low after PTCy even 2 years after the transplant, the specific dynamics of the alloreactive T cell clones are not well defined^26,28^.

To characterize the impact of PTCy specifically on alloreactive and non-alloreactive T cell clones, we compared the clonal frequencies in unstimulated donor samples, at day 3 (D3, prior to administration of PTCy), and at the engraftment time point after PTCy treatment (generally around day 14). We discovered that PTCy results in unique dynamics of the alloreactive T cell repertoire (**Fig. 3A, B**). Alloreactive clones - which had very low cumulative frequencies in most unstimulated donors (0.03%-0.5%) - were expanded in all patients on D3, with a significantly higher ratio of alloreactive to non-alloreactive cumulative clonal frequency in comparison to that of the donor (p < 0.01), demonstrating T-cell allo-antigen recognition and preferential clonal expansion immediately post-transplant **(Fig. 3A).**

**Figure 3.**
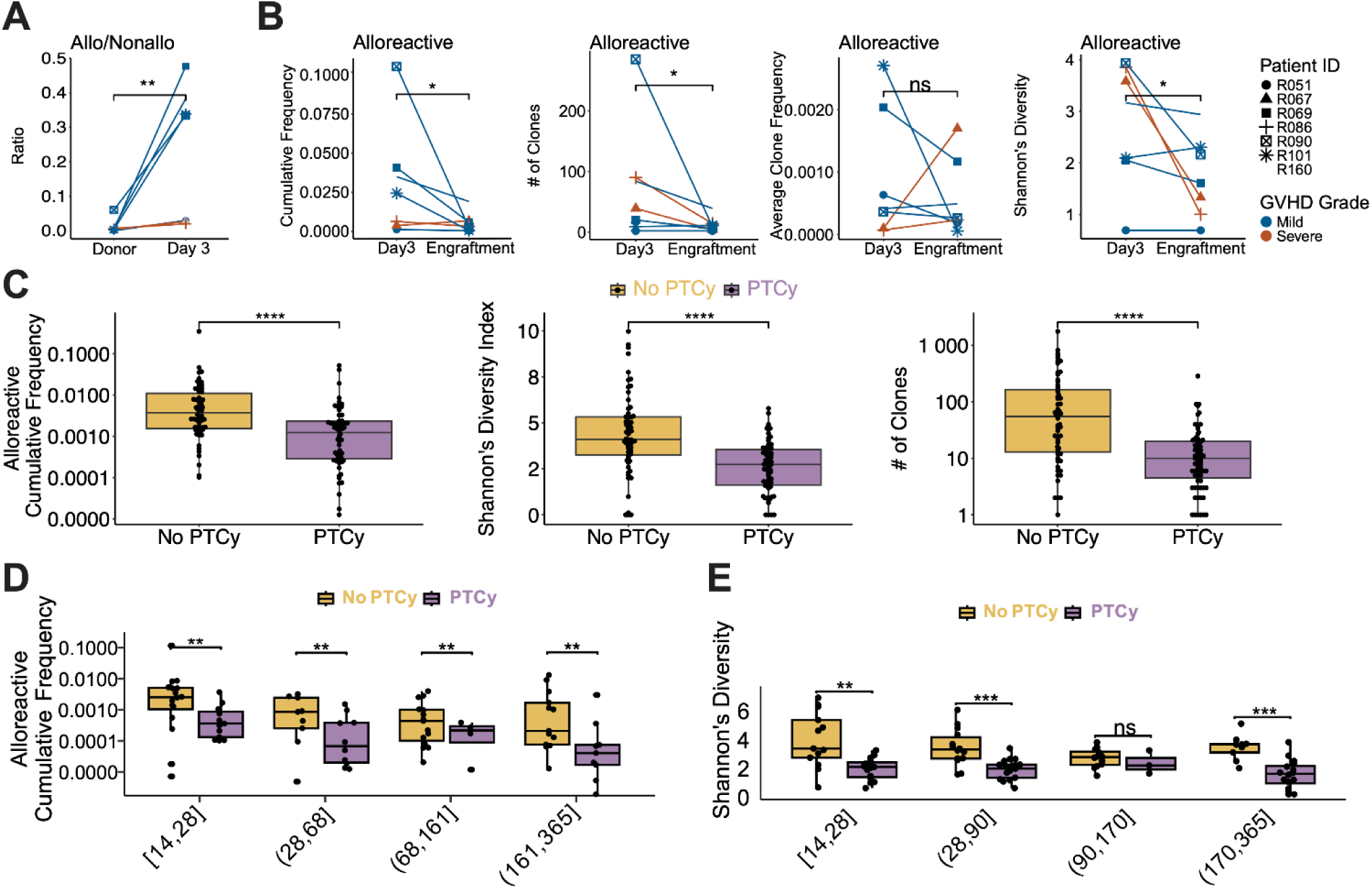
PTCy Selectively Depletes Expanding Alloreactive T Cell Clones. (**A**) Cumulative frequency of alloreactive clones normalized to non-alloreactive clones on donor sample and Day 3 (paired t-test, one-sided, R090 was excluded due to an abnormally high ratio, indicating it was an outlier.). (**B**) Analysis of cumulative frequency, number of clones, average clone frequency, and Shannon’s diversity for alloreactive clones comparing D3 and the engraftment time point in PTCy-treated patients. (**C**) Comparison of cumulative frequency, Shannon’s diversity index, and number of unique clones between PTCy-treated and untreated patients. (**D**) Temporal comparison of cumulative alloreactive frequency across multiple time points (Benjamini-Hochberg adjusted Wilcoxon rank-sum test). (E) Temporal dynamics of Shannon’s diversity were analyzed for all clones, comparing PTCy-treated patients with untreated patients (Benjamini-Hochberg adjusted Wilcoxon rank-sum test,* P < 0.05, ** P < 0.01, *** P < 0.001, **** P < 0.0001).

Following the administration of PTCy, we observed a significant reduction in the cumulative frequency, number and diversity of alloreactive clones (p < 0.0006; **Fig. 3B**). Of the seven PTCy patients, the two that developed severe GVHD demonstrated the least expansion of alloreactive clones on D3 and no significant reduction after PTCy, which could explain the development of severe GVHD despite treatment. In contrast, non-alloreactive clones showed no significant difference following PTCy in cumulative frequency, number of unique clones, average clone frequency or overall clonal diversity **(Supplementary Fig. 6A)**. Collectively, these findings show that PTCy preferentially depletes alloreactive clones and suggest that this depletion is contingent upon early clonal expansion - consistent with the mechanism of action of cyclophosphamide whose active metabolites cause DNA damage preferentially in dividing cells^28^.

We further compared PTCy-treated and untreated patients to delineate broader impacts on the alloreactive repertoire. Patients receiving PTCy demonstrated consistently lower cumulative frequencies, reduced numbers of expanded alloreactive clones, and decreased Shannon diversity compared to those of untreated individuals **(Fig. 3C)**, findings observed at the earliest time point of engraftment and remained consistent across multiple time intervals post-transplant (**Fig. 3D, E, Supplementary Fig. 6B**). Untreated patients displayed persistence of a greater number of non-alloreactive clones, confirming that PTCy’s selectivity predominantly targets the alloreactive component of the T cell repertoire (**Supplementary Fig. 6C**). Notably, the long-term reduction in Shannon’s diversity following PTCy was driven predominantly by a loss of diversity in alloreactive CD4⁺ T cells rather than CD8⁺ T cells, especially evident at early post-transplant time points (days 14–28; **Supplementary Fig. 6D**). PTCy also consistently lowered the number of unique clones compared to no PTCy patients over time, and the impact on CD8^+^ clones was still observed at the later time points (**Supplementary Fig. 7A-D**). These data thus reveal that while PTCy preferentially modulates alloreactive clonal dynamics, incomplete depletion or breakthrough expansion of certain pathogenic clones may underlie severe GVHD events. This highlights the importance of investigating individual clonal dynamics in evaluating prophylactic strategies.

### Enrichment and expansion of CD8^+^ effector and proliferating T cells in gut tissue affected by severe GVHD

Following our analysis of T cell clonal dynamics in the blood, we then sought to investigate how these circulating clonal features translate into altered tissue infiltration and T cell phenotype in GVHD-affected organs, focusing on the GI tract, which is most commonly involved in severe GVHD. We performed single-cell transcriptomic and TCR profiling of immune cells on 50 samples within 28 unique gut tissue specimens from 14 transplant recipients experiencing GI symptoms in addition to 2 normal donors (ND) (**Supplementary Table 4,5,6**; **Methods**, **Supplementary Fig. 8A**). We annotated immune populations using established lineage markers (**Fig. 4A; Methods, Supplementary Table 7**), and observed a marked depletion of B cells in post-transplant biopsies and an increasing trend of T cell proportion among the immune populations with an increase in GVHD severity (**Fig. 4B**).

**Figure 4.**
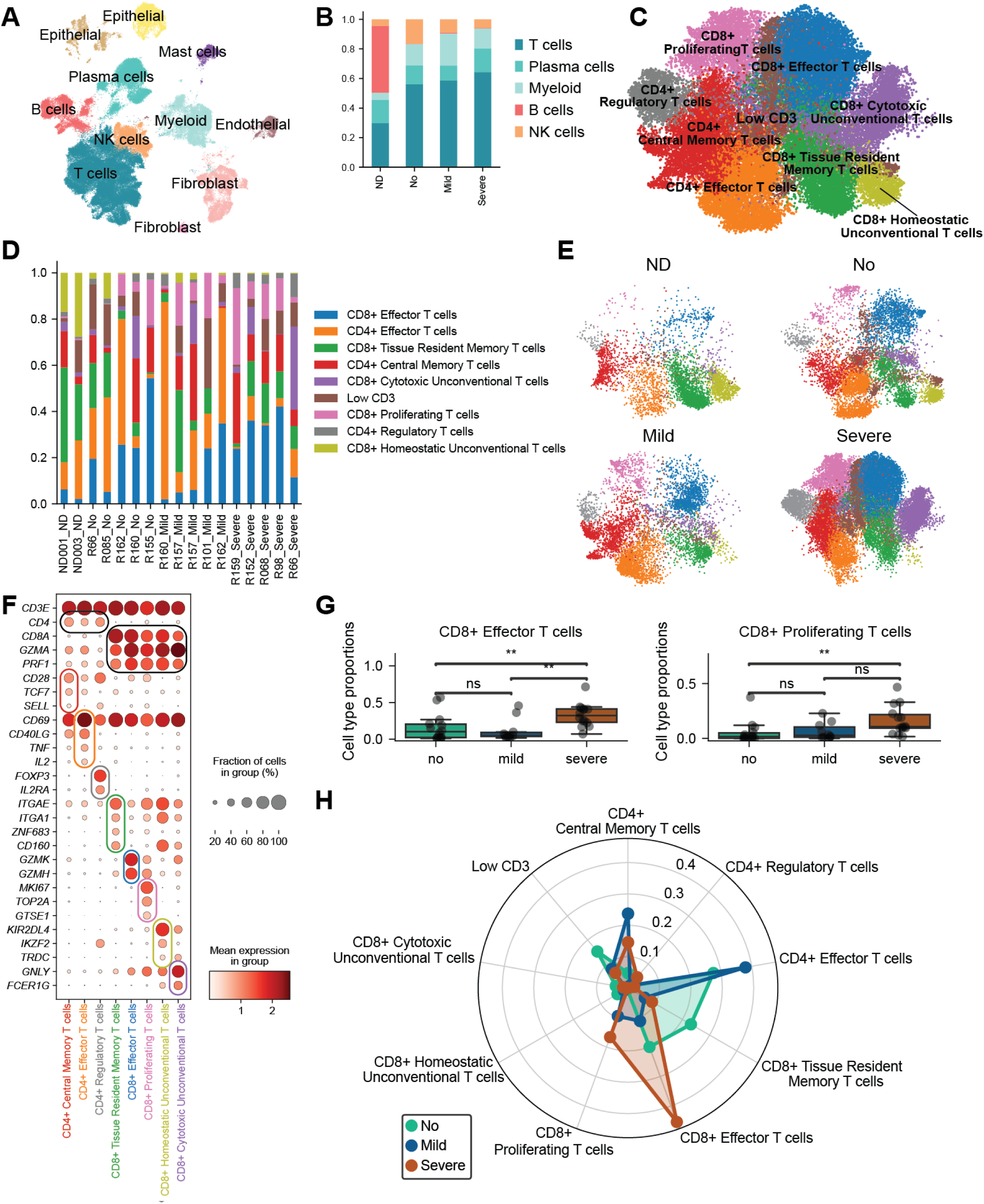
Enrichment of T Cell States in Severe GVHD. (**A**) UMAP visualization of scRNA-seq data from gut tissue biopsies with cell type annotation. (**B**) Proportions of immune cell populations across different GVHD grades. (**C**) UMAP visualization highlighting T cell states. (**D**) Proportions of cell states for each patient. (**E**) UMAP representation of cells stratified by GVHD grades versus normal donor (ND). (**F**) Representative gene list utilized for cell type annotation. (**G**) Cell type proportions for CD8^+^ effector T cells and CD8^+^ proliferating T cells within all T cells grouped by GVHD grade; each dot represents an individual sample (no GVHD and ND were combined as no). (**H**) Radar plot showing proportions of various T cell states by GVHD grade (no GVHD and ND were combined as no).

Given the enriched infiltration of T cells in severe GVHD tissues (**Fig. 4B**), we further examined their phenotypic states through subclustering of the T cell compartment. T cell states were annotated based on differentially expressed genes and classical marker genes (**Fig. 4C, D, F, Supplementary Table 7, Supplementary Fig. 9**). Within the CD8^+^ T cell compartment, five distinct cell states were observed with varying expression levels of effector molecules, integrins, proliferation, and activation markers. Similarly, we identified three distinct CD4^+^ T cell states (**Fig. 4F**).

In ND tissue, CD8^+^ T cells were predominantly resident memory T cells (Trm), expressing classical resident markers such as *ITGAE, ITGA1, CD69* and *CD160*, and CD8^+^ homeostatic unconventional T cells, characterized by expression of γδ TCRs, resident integrins, *IKZF2* and killer receptors like *KIR2DL4* (**Fig. 4E, F**). In contrast, post-transplant tissues showed a predominance of CD8^+^ effector T cells characterized by reduced expression of resident integrins and upregulation of *ITGB2*, consistent with an infiltrating phenotype. Of note, these effector cells had significantly elevated expression of classic cytotoxic markers (such as *GZMA, GZMB, PRF1*, and *GNLY)* and were highly enriched for *GZMK* and *GZMH*. The upregulation of *GZMK* in these cells is particularly relevant given the relatively recent documentation of *GZMK^+^*cytotoxic T-cells in several chronic inflammatory pathologies and ‘inflammaging’^29–33^. Additionally, a CD8^+^ proliferating T cell population (expressing *MKI67, TOP2A,* and *GTSE1*) was nearly absent in normal donors but was significantly expanded in post-transplant biopsies. Notably, a CD8^+^ cytotoxic unconventional T cell population was present in severe GVHD, expressing elevated effector molecules (*GZMA, PRF1, GNLY, FCER1G)* (**Supplementary Fig. 10**), and was largely absent from tissues of ND or no GVHD patients.

Comparative analyses revealed significant shifts in the composition of tissue-infiltrating T cell states in post-transplant biopsies across GVHD severity. Remarkably, CD8^+^ effector and proliferating T cells were significantly enriched in severe GVHD patients compared to mild or no GVHD patients (p < 0.01, **Fig. 4G**). Mild GVHD tissue, in contrast, is marked by a predominance of CD4^+^ central memory and effector T cells, and the no GVHD group exhibits a predominance of CD8^+^ tissue resident T cells, homeostatic unconventional T cells and CD4^+^ effector T cells (**Fig. 4H, Supplementary Fig. 11A**). These differences highlight shifts in T cell state composition with disease severity.

TCR clonal profiling showed that CD8^+^ effector T cells and proliferating CD8^+^ T cells have lower clonal diversity (higher Gini index) in severe GVHD patients, suggesting these specific T cell states are enriched due to clonal expansion as opposed to other subsets (**Supplementary Fig. 11B**). These results thus link systemic clonal expansion with tissue-specific enrichment of cytotoxic and proliferating CD8⁺ T cells. Although similar enrichment of these subsets was observed irrespective of prior PTCy treatment, PTCy-treated patients exhibited reduced CD8⁺ Trm populations, possibly reflecting delayed reconstitution of immune homeostasis in the tissues (**Supplementary Fig. 11C**).

### Proinflammatory reprogramming of unconventional and regulatory T cell states

Beyond shifts in conventional T cell proportions, we observed a striking phenotypic transformation within the unconventional T cell compartment in patients with GVHD. Intestinal unconventional T-cells, characterized by high expression of NK receptors and the Treg-associated transcription factor Helios (*IKZF2*) ^34,35^, are enriched in the intraepithelial compartment where they play critical roles modulating tissue homeostasis by promoting mucosal barrier function and tissue repair in response to injury^36^. Of note, we found that homeostatic unconventional T-cells were largely depleted in patients with GVHD (**Fig. 4E, Supplementary Fig. 11A**). Additionally, GVHD was associated with the transition of unconventional T cells towards a highly altered cytotoxic state in a select number of patients (**Supplementary Fig. 12 and Supplementary Table 8)**. This phenotypic shift was marked by a metabolic shift from oxidative phosphorylation to glycolysis (Upregulated: *LDHA, GAPDH, SLC2A3,* Downregulated: *NDUFA1, COX14, ATP5ME)*, alongside an upregulation of activating signaling adapters (*TYROBP, FCER1G*), cytolytic molecules (*GZMK, GZMA, GNLY*), pro-inflammatory cytokines (*TNF, IFNG*), cytoskeletal remodeling genes (*PFN1, CFL1, RAC2, TMSB4X*), and an altered integrin profile (elevated: *ITGB2,* reduced: *ITGAE*). Furthermore, we observed a significant downregulation of hallmark killer receptors (*KIR2DL4, KIR3DL2, KLRC2*), as well as genes associated with regulatory functions (*IKZF2, ENTPD1, NT5E*) and tissue regeneration (*AREG*). In addition, the glucocorticoid receptor *NR3C1* was downregulated in these altered unconventional T cells, potentially contributing to steroid resistance often seen in severe GVHD. To this end, cytotoxic unconventional T cells were particularly dominant (∼36% of all tissue-infiltrating T-cells) in patient R66, who experienced highly aggressive steroid-refractory disease. Notably, a subsequent biopsy from this patient following GVHD resolution showed the disappearance of this cytotoxic population and the re-emergence of its homeostatic counterpart (**Fig. 4D** – sample R66_no). Together, these findings suggest that gastrointestinal GVHD is associated with profound disturbance of the intestinal unconventional T-cell pool, with these cells either being lost or highly altered to an activated state with elevated motility and proinflammatory potential.

Surprisingly, we observed that regulatory T cells (Tregs) are also enriched in severe GVHD within CD4^+^ T cells (p < 0.01, **Supplementary Fig. 13D**) but display a profoundly altered phenotype. Although Tregs in severe GVHD maintain classical markers (e.g., *FOXP3*, *IKZF2*, *IL2R* and *CTLA4*), analysis of differentially expressed genes (DEG) revealed an atypical upregulation of a cytolytic signature (*PRF1*, *GZMA*, *GZMB*), type I interferon response genes (*GBP2*, *GBP5*, *IRF1*, *IRF9*), and receptors for pro-inflammatory cytokines such as IL-1, IL-18 and IL-6 (**Supplementary Fig. 13G and Supplementary Table 9, 10)**. In addition, we observed downregulation of oxidative phosphorylation genes (e.g., *COX* genes and ATP synthase subunits), indicating an atypical metabolic state. Interestingly, expression of the signaling mediators *JAK1*, *STAT1,* and *STAT3* was also elevated in Tregs from severe GVHD compared to other conditions, further suggesting functional reprogramming of Tregs towards a pro-inflammatory phenotype (p < 0.03).

These alterations point to a breakdown in tissue immune homeostasis and suggest convergent mechanisms of effector and regulatory T cell dysfunction driving severe disease. Specifically, both effector and regulatory compartments are skewed toward a pro-inflammatory, cytotoxic phenotype in severe GVHD, thereby contributing to immune dysregulation and tissue damage.

### CD8^+^ T cell clones exhibit phenotypic plasticity and mobility across tissue compartments

The observed clonal expansion of CD8^+^ effector and proliferating T cell states in GVHD-affected gut tissue prompted us to investigate whether these expansions reflect local proliferation or *in situ* differentiation. As our single-cell sequencing was performed on T cells isolated separately from the lamina propria (LP) and intraepithelial (IE) compartments (see **Methods**), we defined “mobile” clones as those shared across both compartments. Across all T cells, mobile clones displayed lower clonal diversity (higher Gini index) compared to clones restricted to a single compartment, indicating clonal expansion (**Supplementary Fig. 11D**). Among conventional CD8⁺ T cells, some of the most expanded mobile clones were detected in multiple phenotypic states suggesting *in situ* state transition as cells move between LP and IE tissue compartments **(Fig. 5A).** Notably, we observed enrichment of mobile clones in the Trm subset (p = 5.4e-180; **Supplementary Fig. 11D**). Further sub-clustering within the CD8 mobile Trm cells identified a distinct subset (orange cluster in **Fig. 5B**) that co-expressed canonical residency markers such as *ITGAE* and cytolytic molecules alongside multiple exhaustion markers (e.g., *PDCD1 [PD-1], CTLA4, HAVCR2 [TIM-3]*), while showing low *IL-7R* expression (**Fig. 5C**). This co-expression of exhaustion markers alongside retained cytotoxic markers (*GZMA, GZMB*) suggests a state of chronic activation, distinct from quiescent Trm cells. This phenotype likely reflects cells progressing towards exhaustion under persistent stimulation within the GVHD microenvironment while maintaining effector functions. Analysis of transcription factor expression within the CD8^+^ Trm sub-clusters revealed that this population expressed *PRDM1* (BLIMP-1) and *RUNX3* at levels comparable to other Trm cells, but was distinguished by high expression of *ZNF683* (Hobit) (**Supplementary Fig. 14**). Interestingly, the Hobit+ resident memory, along with CD8^+^ proliferating T cells, are uniquely identified in our dataset (**Supplementary Fig. 15**), as we explored annotated single-cell RNA Seq data from IBD and healthy gut tissue^37,38^.

**Figure 5.**
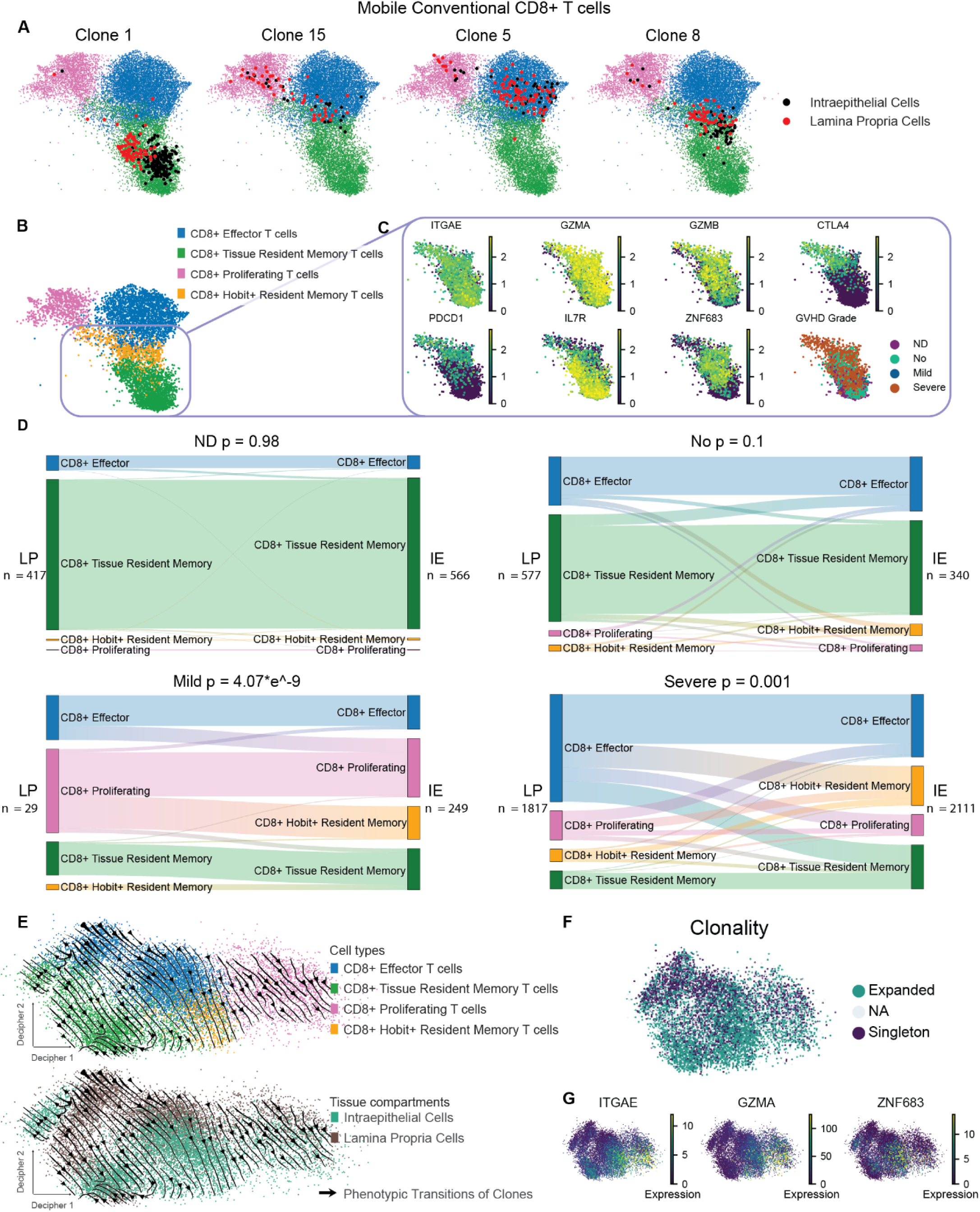
Phenotypic Plasticity of T Cells Across Tissue Compartments in GVHD Patients. **(A)** UMAP projections of CD8^+^ T cell populations, color-coded by cluster annotation. Key expanding clones are marked (Black: IE, Red: LP). **(B)** UMAP plot displaying CD8^+^ T cell subpopulations, including effector T cells, tissue resident memory T cells, proliferating T cells, and Hobit^+^ resident memory T cells. **(C)** Expression profiles of key genes (*ITGAE, GZMA, GZMB, CTLA4, PDCD1, IL7R)* across GVHD severity grades, shown with corresponding color intensity scales. **(D)** Sankey diagrams illustrating cell state transitions across GVHD severity groups (ND, no, mild, severe, Fisher’s exact test, n is the number of cells). **(E)** Decipher analysis projecting CD8^+^ T cell populations based on Decipher 1 and 2 axes. Arrows represent the transition of shared clones from LP to IE. **(F)** T cell clonality in Decipher space showing expanded clones and singletons. (G) Gene expression of *ITGAE, ZNF683*, and *GZMA* is plotted on the decipher space.

We next examined T cell clonal expansion across different GVHD grades and found that the homeostatic Trm population appeared to be the most expanded across all grades (**Supplementary Fig. 16**). However, each grade displayed distinct expansion patterns. In severe GVHD, CD8^+^ effector T cells were predominantly expanded, whereas mild or no GVHD samples were enriched in non-expanding CD4^+^ effector T cells. Additionally, no and ND samples showed increased expansion in CD8^+^ Trm and CD8^+^ homeostatic unconventional T cells.

Subsequently, we systematically investigated phenotypic shifts in mobile conventional CD8^+^ T cell clones across different grades of GVHD. In ND, T cells within the same clone showed no significant phenotypic change between the LP and IE compartments (p = 0.98). Similarly, transplant patients without GVHD displayed no significant phenotypic change (p = 0.1). In contrast, we observed significantly more dynamic phenotypic shifts between LP and IE compartments in the mild and severe GVHD patients (p < 0.001; **Fig. 5D**). This phenotypic plasticity is mainly observed among CD8^+^ effector and proliferating T cell clones. Meanwhile, the Trm population largely maintains its phenotype across compartments, with the exception of the CD8^+^ Hobit Resident Memory T cells that are rarely observed in ND, and display frequent transitions with effector and proliferating cell states across tissue compartments. Together, these results suggest that GVHD is associated with dynamic phenotypic shifts and cell state transitions across tissue compartments, resulting in a chronically activated CD8^+^ Trm subset, characterized by high *ZNF683* expression in the intraepithelial layer.

To contextualize the observed phenotypic plasticity within the T cell differentiation and activation trajectory, we employed Decipher^23^, an analytical tool that characterizes both shared and unique transcriptional programs across LP and IE compartments while preserving the global geometry of T cell trajectories in a 2D embedding space generated directly by the model (**Fig. 5E**). The Decipher 1 component captures shared T cell state transitions from resident towards effector and proliferating phenotypes coinciding with elevated expression of the cytotoxic marker *GZMA* (**Fig. 5E, G**), while the Decipher component 2 captures the difference between tissue-specific T cell trajectories (LP and IE). The dominant migration of CD8^+^ effector cell clones from the LP into the epithelial compartment is accompanied by the acquisition of residency signatures, including expression of *ITGAE* and *ZNF683* (Hobit)^39–41^(**Fig. 5G**).

To better understand this clonal differentiation and migration, we incorporated single-cell TCR data in the Decipher 2D space, using our DecipherTCR tool, to investigate whether T cells sharing the same TCR exhibit transcriptional shifts as they move from LP to IE regions. This integrative analysis captures trajectories of phenotypic state transitions of dominant T cell clones in the transcriptomic embedding space generated from scRNA-seq data (**Methods**), which revealed that CD8^+^ effector T clones change phenotypes and differentiate into Hobit^+^Trm by acquiring tissue-resident transcriptional signatures upon migrating from the LP to the IE region, where they appear more clonally expanded (**Fig. 5E, F, G**). This phenotypic shift, previously uncharacterized in human GVHD, likely promotes their retention within the epithelium^42^, consequently contributing to epithelial damage. Interestingly, we further observed more subtle transitions between Hobit^+^ Trm cells and CD8^+^ proliferating T cells in smaller clones (<20 cells; **Supplementary Fig. 17D**).

### CD8^+^ effector T cell hubs spatially localize with intestinal stem cell niches in severe gut GVHD

Since work in murine models identified intestinal stem cells as critical targets in gut GVHD pathology^43,44^, to further understand how CD8**^+^** effector T cells contribute to tissue damage, we leveraged high-resolution spatial transcriptomics (10x Genomics Visium HD, 2µM) to map their localization. We analyzed eight gut biopsies: four from mild GVHD cases, three from severe GVHD cases, and one ND sample (**Supplementary Table 11**). Initial clustering of spatial coordinates based on transcriptomic measurements recapitulated histological structures of gastrointestinal tissue (**Fig. 6A**). However, these clusters reflected mixed cell populations that could not resolve distinct immune cell types, possibly due to lateral RNA diffusion across spots and overlapping cells in the 3D tissue architecture^45,46^ (**Supplementary Fig. 18A**). This problem persisted even with binning spots according to segmented cells^47^ (**Supplementary Fig. 19**). To enable fine-grained spatial resolution of cell states, we adapted our tool Starfysh^24^, a deep generative model for deconvolving refined cell states from integration of spatial transcriptomic data and histology images. Our new computational framework, StarfyshHD, expands its application to smaller spot sizes as in Visium HD data (**Methods**).

**Figure 6.**
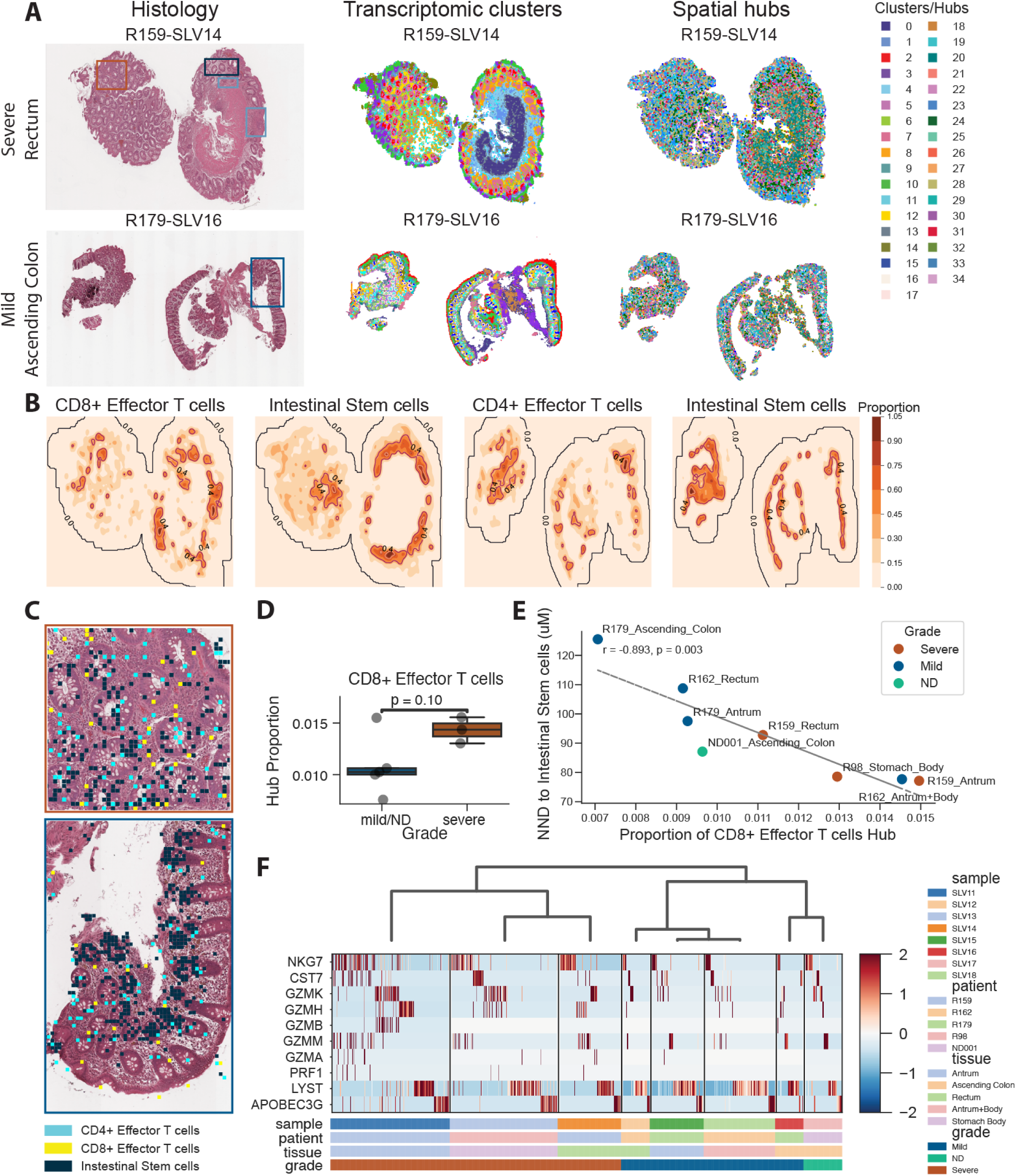
CD8^+^ Effector T Cells Cluster in Proximity to Intestinal Stem Cells in Severe GVHD. **(A)** Histology, transcriptomic clusters, and spatial hubs in representative colon samples from severe (SLV14) and mild (SLV16) GVHD patients. **(B)** Contour plots showing CD8^+^/CD4^+^ Effector T cell and ISC proportions from Starfysh deconvolution. **(C)** Zoomed-in views of SLV14 and SLV16 highlight hubs enriched for CD8^+^/CD4^+^ Effector T cells and ISCs. **(D)** Box plot comparing CD8^+^ Effector T cell proportions between severe and mild/no GVHD (t-test). **(E)** Scatter plot of mean nearest neighbor distance (NND) from CD8^+^ Effector T cells to ISCs versus CD8^+^ Effector T cell proportion. **(F)** Heatmap displaying CD8^+^ Effector T cell signature expression across all samples.

A key advantage of Starfysh is its ability to integrate spatial transcriptomic data across samples and identify shared spatial hubs with similar cell type compositions (**Supplementary Fig. 20**)^24^. By leveraging gene signatures generated from our single-cell RNA-Seq data (**Supplementary Table 7**), StarfyshHD identified spatial hubs enriched for specific cell states, including intestinal stem cells (ISCs) and CD8^+^ effector T cells (prominent in severe patient R159) and CD4^+^ effector T cells (prominent in mild patient R179; **Fig. 6A, B; Supplementary Fig. 18C**). Notably, the identified ISC-enriched hub aligns with crypt base locations within the tissue architecture, validating the deconvolution approach (**Fig. 6C, Supplementary Fig. 21A, B**).

Comparative analysis across all eight samples showed a trend toward higher proportions of CD8^+^ effector T cell hubs in severe GVHD compared to mild GVHD patients or normal donors (p = 0.1; **Fig. 6D**). While this finding did not reach statistical significance (possibly due to limited sample size), it is consistent with our discovery in the scRNA-seq data (**Fig. 4G**). We also observed a correlation between the proportion of CD8^+^ effector T cell hubs and their mean nearest neighbor distance to ISC hubs, suggesting that a higher abundance of CD8^+^ effector T cells is associated with closer proximity to ISCs (**Fig. 6E**). Interestingly, one outlier sample (from patient R162, classified as mild GVHD at the time of biopsy) exhibited both a high proportion of CD8^+^ effector T cell hub and close proximity to ISCs. Notably, this patient subsequently progressed to more significant GVHD involving upper and lower GI tract as well as the skin, requiring high-dose steroids. Furthermore, hierarchical clustering based on gene signatures of those CD8^+^ effector T cell-enriched hubs reveals that hubs from severe GVHD samples generally exhibited higher expression of cytotoxic genes and clustered separately from mild GVHD samples (**Fig. 6F**). Thus, regions enriched for CD8^+^ effector T cells in severe GVHD patients demonstrate a more pronounced cytotoxic transcriptional profile.

### Immune circuit rewiring at the site of crypt loss in severe GVHD gut tissue

To further explore the spatial interplay between immune effectors and tissue damage, we examined a rectal biopsy (sample SLV14) from a severe GVHD patient (R159), containing well-demarcated regions of crypt loss with some residual crypts showing increased crypt apoptosis. We performed region-specific spatial analyses comparing crypt-intact and crypt-loss areas within this sample (**Fig. 7A**). Mapping the distribution of epithelial and T cell types revealed a striking enrichment of CD8^+^ effector T cells specifically within the crypt-loss regions compared to crypt-intact areas, while ISCs were significantly depleted in crypt-loss regions, as expected (**Fig. 7A, B**, p < 0.0001).

**Figure 7.**
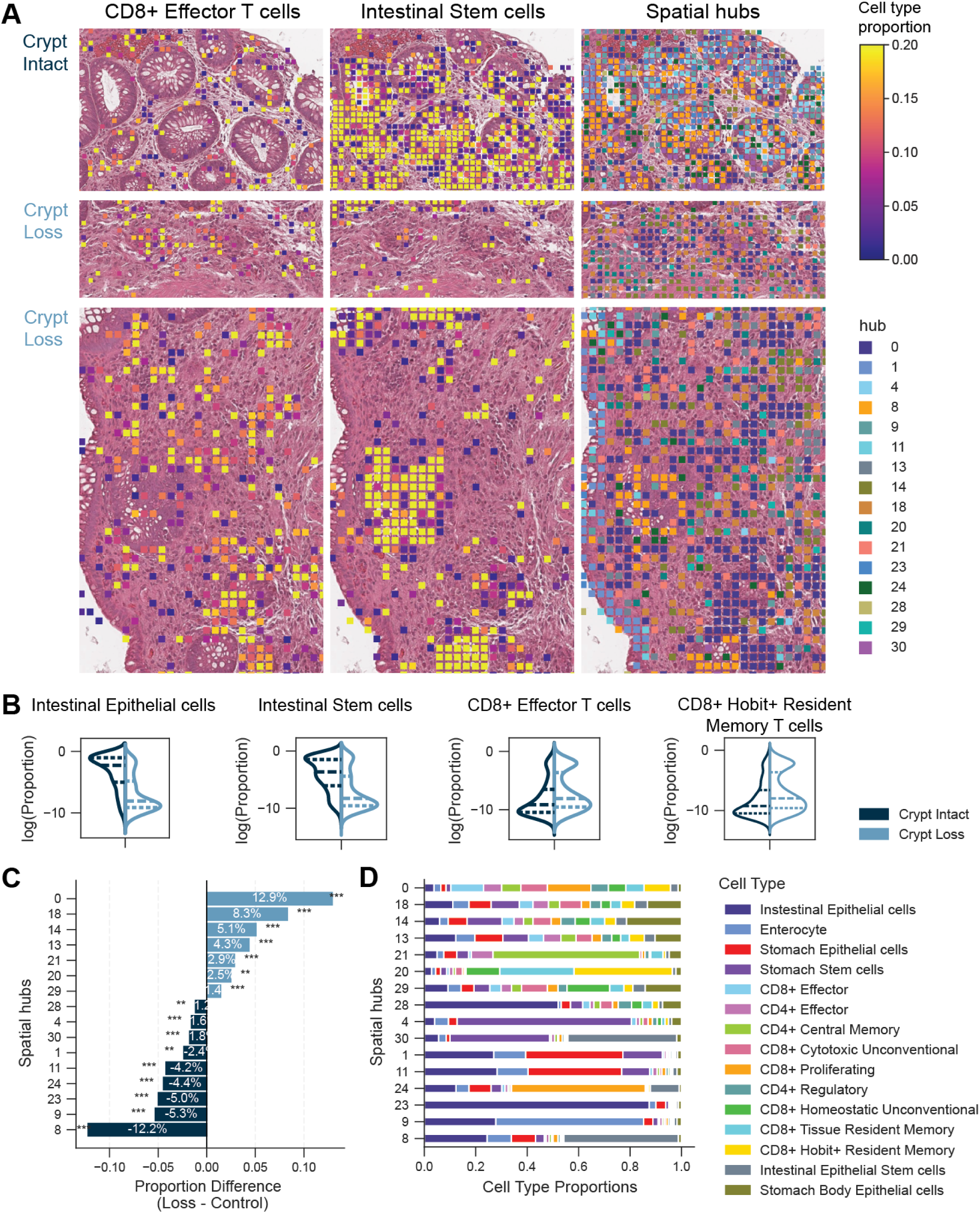
Collaborative T Cell Response in Crypt Loss Regions. **(A)** CD8^+^ Effector T cell (left) and ISC (middle) proportions inferred from StarfyshHD, along with spatial hubs that are enriched in either crypt intact and crypt loss region (right, see C) in crypt loss and intact regions of SLV14 (crypt loss contains area of crypt loss and increase crypt apoptosis). Histology images are shown in the background. **(B)** Violin plots comparing cell type proportions between crypt loss and intact regions (Mann-Whitney U-test, p<0.0001). **(C)** Hubs enriched or depleted in crypt loss regions (Fisher’s exact test). **(D)** A stacked bar plot shows cell type composition in the differentially abundant hubs.

We next assessed cellular hubs in these regions. Hub 0 was enriched considerably in crypt loss areas, while hub 8 - dominated by epithelial cells and ISCs - was significantly depleted (**Fig. 7A,C)**. To understand the cellular composition of this pathology-associated hub, we examined the deconvoluted subsets. We found that hub 0 consisted of diverse T cell subsets, including CD8^+^ effector, CD8^+^ proliferating, CD8^+^ Trm, CD4^+^ effector, and CD4^+^ central memory T cells, whereas hub 8 consisted mainly of intestinal epithelial cells and ISCs (**Fig. 7D**). This is consistent with a coordinated immune response, either leading to tissue destruction or participating in a tissue repair effort.

We then further explored interactions among cellular hubs in crypt loss and crypt intact regions using CellPhoneDB^48^, a repository of ligand-receptor interactions. We identified significant interactions between ISCs, CD4^+^ effector T cells hub and Tregs hub in crypt intact regions, possibly indicating a more restrained cytolytic immune response (**Supplementary Fig. 22**). In contrast, the crypt loss regions had stronger interactions between intestinal epithelial stem cells hub and CD8^+^ effector T cells hub, which supports our previous finding in association of CD8^+^ effector T cells hub. Interestingly, we observed interactions between hub 0 and ISCs in crypt loss regions but not in crypt intact regions, suggesting altered epithelial-immune communication during crypt damage.

These observations support a model in which expanded and spatially localized CD8^+^ effector T cell populations are the direct mediators of epithelial injury in human gut GVHD^49^, particularly targeting ISC-rich niches. Moreover, the presence of diverse T cell subtypes within injury-associated hubs raises the possibility of both destructive and compensatory immune functions in the context of GVHD. Altogether, spatial transcriptomic analysis uncovers tissue-level immune organization associated with GVHD severity and highlights the pathological relevance of effector T cell proximity to epithelial stem cell compartments.

## Discussion

In this comprehensive spatiotemporal analysis of human graft-versus-host disease (GVHD), we redefine the disease as a dynamic interplay of clonal expansion, phenotypic plasticity, and spatial immune rewiring. By integrating pre-transplant alloreactive T cell identification with longitudinal tracking and high-resolution spatial transcriptomics, we provide the first human framework to resolve how donor T cells evolve into pathogenic clones and interact with host tissues. This approach offers a rare window into the *in vivo* dynamics of antigen-driven human T cell responses in clinically annotated patients and provides a framework that extends beyond GVHD to other immune-mediated diseases, including cancer, infection, and autoimmunity.

This study introduces a two-tiered strategy for T cell repertoire analysis: biological enrichment through mixed lymphocyte reactions (MLR) combined with computational modeling using our novel tool, DecompTCR, enabling the identification of pathogenic clones with distinct biophysical and transcriptional characteristics. T cell clones exhibiting temporal patterns associated with severe GVHD had distinct TCR characteristics, including shorter CDR3β length and lower hydrophobicity. These features echo properties described for publicly shared, cross-reactive T cells^50,51^, suggesting that increased risk for severe GVHD may involve donor T cells with a greater capacity for promiscuous recognition of recipient allo-antigens. DecompTCR proved effective in differentiating GVHD severity based on the temporal dynamics of alloreactive clones without using clinical data, highlighting its potential for longitudinal T-cell repertoire analysis and, more broadly, for identifying clonal trajectories in immune-mediated pathology.

Our study also interrogated the impact of PTCy, which is commonly used for GVHD prophylaxis. The selective effect that we demonstrated on the frequency, number, and diversity of alloreactive clones aligns with and provides human *in vivo* support for the hypothesis that PTCy preferentially targets proliferating alloreactive clones^26,52^. Our findings also suggest that this mechanism does not work effectively if early expansion of alloreactive clones has not occurred, thereby exposing the patient to severe GVHD and supporting a potential use of this finding as a biomarker. These results highlight how therapeutic outcomes are shaped by early clonal behavior, reinforcing the importance of temporally resolved clonal data to understand treatment resistance, and inform early intervention and patient stratification strategies.

Moving from peripheral blood to the site of tissue damage, our single-cell analysis of gastrointestinal tissue revealed a marked increase in infiltrative, clonally expanded cytotoxic CD8^+^ T cells in GVHD, compared to healthy tissue. We identified proliferating T cells along with an enrichment of clonally expanded *GZMK*^+^ CD8^+^ effector T cells in severe GVHD^53^. Though its role is less described than other lytic molecules^54,55^, Granzyme K has been implicated in inducing targeted cell death and complement-mediated tissue inflammation in multiple disease processes^30,56–62^. The expansion of these cells in GVHD reinforces a broader role for non-canonical cytotoxic programs in immune-mediated tissue injury.

We observed a shift from CD8⁺ homeostatic to cytotoxic unconventional T cells during GVHD. The homeostatic population, enriched in normal donors, is associated with tissue protection and immune homeostasis. Its loss post-transplant, and, in some cases, an enrichment of a cytotoxic population, suggests that the inflammatory environment in GVHD drives unconventional T cells toward cytotoxic tissue-damaging function. This transition highlights the plasticity of unconventional T cells under inflammatory pressure and may reflect a generalizable mode of immune circuit rewiring under chronic stimulation.

Intriguingly, we observed complex dynamics within the Treg compartment. Our experimental finding of Treg enrichment in severe GVHD has been demonstrated previously, but was thought to be simply insufficient to counterbalance CD8^+^ T cell–driven inflammation^53^. Our analysis uncovered a significant phenotypic rewiring of Tregs present in severe GVHD. Despite maintaining canonical Treg markers, they atypically upregulated cytolytic genes (*GZMA*, *PRF1*) and pro-inflammatory signaling pathways^63–65^. This suggests that in severe GVHD, Tregs adopt dysfunctional, pro-inflammatory characteristics, contributing to the failure of immune homeostasis rather than effectively suppressing the alloresponse. Cytolytic Tregs were previously identified in the context of inflammatory bowel disease (IBD)^66^, as well as in mouse tumor microenvironments, demonstrating the capacity to induce CD8^+^ T cell death in a Granzyme B and perforin-dependent manner^67^.

A key discovery enabled by our integrated single-cell RNA and TCR sequencing and DecipherTCR tool was the significant T cell phenotypic plasticity within the gut tissue. We demonstrated a novel phenotypic shift between CD8^+^ effector T cells and Trms, particularly in severe GVHD. While most findings of T cell phenotype plasticity have come from animal models^68–77^, this particular shift has been reported in the context of human intestinal transplantation, where it was associated with rejection, and in IBD^78–80^. Our study further revealed an enrichment of *ZNF683* (Hobit)^+^ Trm cells in GVHD patients. Hobit is increasingly recognized as a regulator of cytotoxic T cell function and tissue residency^41,81,82^. In the setting of leukemia, we previously identified *ZNF683*⁺ CD8⁺ T cells as clonally expanded cytotoxic effectors in acute myeloid leukemia patients responding to donor lymphocyte infusions, where these cells coordinated multicellular immune networks and mediated antitumor responses in the bone marrow microenvironment^82^. Hobit expression is modulated by transcription factors such as *EOMES* and defines a transition state during the phenotypic conversion of CD8⁺ effector to Trm in peripheral tissue^42^. It has also been found to regulate key genes involved in T cell activation and cytotoxicity, such as *TCF7*, *LMO2*, and *CD69*, suggesting a direct role in shaping effector function, as well as suppressing tissue exit genes such as *CCR7*, *S1PR1* (**Supplementary Fig. 14**). The enrichment seen in our study, within cells co-expressing activation and exhaustion markers, may signify a chronically activated yet functionally potent tissue localizing T cell state contributing to GVHD pathology, potentially representing an intermediate exhaustion state amenable to therapeutic intervention.

Spatial transcriptomics represents a transformative approach for studying the complex and compartmentalized environments of the gastrointestinal tract. The application of spatial transcriptomics to gut tissues allows for the preservation of intricate intestinal architecture while capturing gene expression *in situ*, which is essential for understanding spatial heterogeneity across the crypt-villus axis and between different anatomical regions. Spatial resolution is particularly relevant as acute gut GVHD involves localized pathologies including epithelial injury, loss of Paneth cells, fibrosis, and altered immune cell organization^83^. While single-cell RNA-seq has revealed shifts in immune and epithelial cell populations in GVHD and IBD, spatial approaches like multiplexed imaging and spatial transcriptomics uniquely resolve localized immune attacks, crypt-specific epithelial damage, and spatial zonation loss, offering insights into disease pathogenesis that bulk or single-cell suspension methods cannot capture^37,84^. To that end, StarfyshHD allows us to deconvolve spatial cell type composition and colocalization within human gut biopsies. Specifically, we discovered unique immune compositions enriched within regions of the crypt loss region and highlighted the proximity of CD8^+^ effector T cells to areas of severe tissue damage in gut GVHD. These findings extend observations from murine models that implicate CD4^+^ and CD8 T^+^ cells co-localizing with crypt regions^43,44^, and reveal in human tissue a predominance of clonally expanded, phenotypically plastic CD8^+^ effector T cells driving epithelial injury and crypt loss.

In conclusion, this study provides a multi-omic, spatiotemporal view of the human T cell response in GVHD. These findings offer potential avenues for developing novel biomarkers based on T cell temporal dynamics or phenotypic plasticity to improve alloHCT outcomes. More broadly, this work offers an integrated experimental and computational blueprint for dissecting immune-mediated tissue pathology, with implications for autoimmunity, transplantation, and cancer immunotherapy.

### Limitations of the study

Our study is limited by clinical constraints, where there is an obvious difficulty producing a great number of longitudinal donor-patient data points. In particular, tissue biopsies from post-transplant patients without any GI symptoms were not available. However, we were able to include in our study biopsies from symptomatic patients who had no pathologic evidence of GVHD and whose symptoms resolved without treatment. TCR clonotypes detected in peripheral blood are identified and sequenced exclusively on the receptor beta chain but not the alpha chain. The beta chain is the most variable and epitope-interacting portion of the TCR, and studies have shown that CDR3α may play little role in antigen recognition^85^. On the other hand, the paired scRNA and TCR data contain both alpha and beta chains, allowing a more precise identification of individual clones.

Technical factors impacted the quality of gut biopsies analyzed by single-cell RNA sequencing, including tissue dissociation and freeze-thaw cycles that may bias cell-type representation. As our focus was on T cells, which are relatively resilient to these effects, this impact was likely minimal. Variability in RNA integrity may have arisen from processing timing. For FFPE samples analyzed with Visium HD, formalin fixation can lead to RNA fragmentation, reducing transcript capture sensitivity.

## Lead contact

Further information and requests for resources should be directed to and will be fulfilled by the Lead Contacts, Elham Azizi, and Ran Reshef.

## Materials availability

This study did not generate new and unique reagents.

## Data and code availability

All data generated in this study are included in this published article and its supplementary information. The data discussed in this manuscript will be deposited in the National Center for Biotechnology Information’s Gene Expression Omnibus (GEO) upon publication. Code relating to data processing and figure generation is available in our GitHub repository. Patient information is available in the supplementary table. Any additional information required to reanalyze the data reported in this paper is available from the lead contact upon request.

## Acknowledgements

We thank the following cores and resources for their support: the Columbia Gastrointestinal Tissue Bank for providing normal donor biopsies, the Human Immune Monitoring Core, the CCTI Biobank Core, the Flow Cytometry Core, and the NMDP for access to unrelated donor samples. We gratefully acknowledge the Columbia and NMDP Institutional Review Boards and all patients who generously consented to participate in this study. We are also grateful to Jianing Fu, Aleksandar Obradovic, Christopher Parks, Susan Dewolf, and Yufeng Shen for helpful discussions.

## Author contributions

L.S., A.U., X.W., E.A., and R.R., conceived the study. X.W., K.B., and A.U. developed MLR protocols and X.W., A.U., S.C., D.W.H performed sample processing and data collection. E.A., L.S., M.P. designed and developed DecompTCR. E.A. and L.S. designed and developed Decipher TCR, and E.A., L.S, and J.A.Z. designed and developed StarfyshHD. R.R., M.M., and C.G. provided clinical samples and data and maintained regulatory oversight. CCTI Human Immune Monitoring Core performed all 10X data acquisition experiments. L.S., M.P., D.W.H., J.A.Z., X.S., J.S.F., T.A., J.H., and P.C. analyzed the data. L.S., M.P., J.A.Z. X.S., J.S.F., T.A., J.H., P.C., R.R., A.U., X.W., D.W.H., S.C., R.M., C.G., E.R., M.M., M.S., and J.L.M. interpreted the data. E.A., R.R., L.S., A.U., D.W.H., and X.W. wrote the manuscript.

## Supplemental information

**Supplementary Table 1.** Patient and Donor Characteristics (n=20)

| Supplementary Table 1. Patient and Donor Characteristics (n=20) |  |
| --- | --- |
| **Transplant Donor Sex** |  |
| Male, n | 16 |
| Female, n | 4 |
| **Transplant Recipient Sex** |  |
| Male, n | 13 |
| Female, n | 7 |
| **Donor Age**, Median (Range) | 29.5 (18 - 64) |
| **Recipient Age**, Median (Range) | 57.5 (28 - 66) |
| **Acute GVHD Grade** |  |
| None, n | 6 |
| Mild (Grade 1-2), n | 8 |
| Severe (Grade 3-4), n | 6 |
| **Chronic GVHD Grade** |  |
| None | 15 |
| Mild | 3 |
| Moderate-Severe | 2 |
| **Underlying Disease** |  |
| Acute Myeloid Leukemia, n | 12 |
| Chronic Myelomonocytic Leukemia, n | 1 |
| Myelofibrosis, n | 2 |
| Myelodysplastic Syndrome, n | 2 |
| Cutaneous T-cell Lymphoma, n | 1 |
| T-Cell Prolymphocytic Leukemia, n | 1 |
| Sickle Cell Disease, n | 1 |
| **Donor Type** |  |
| Matched Unrelated (10/10 HLA), n | 5 |
| Matched Sibling (10/10 HLA), n | 2 |
| Haploidentical, n | 12 |
| Mismatched Unrelated (9/10 HLA), n | 1 |
| **Post Transplant Cyclophosphamide** |  |
| Yes, n | 12 |
| No, n | 8 |

### Supplementary figures

**Supplementary figure 1.**
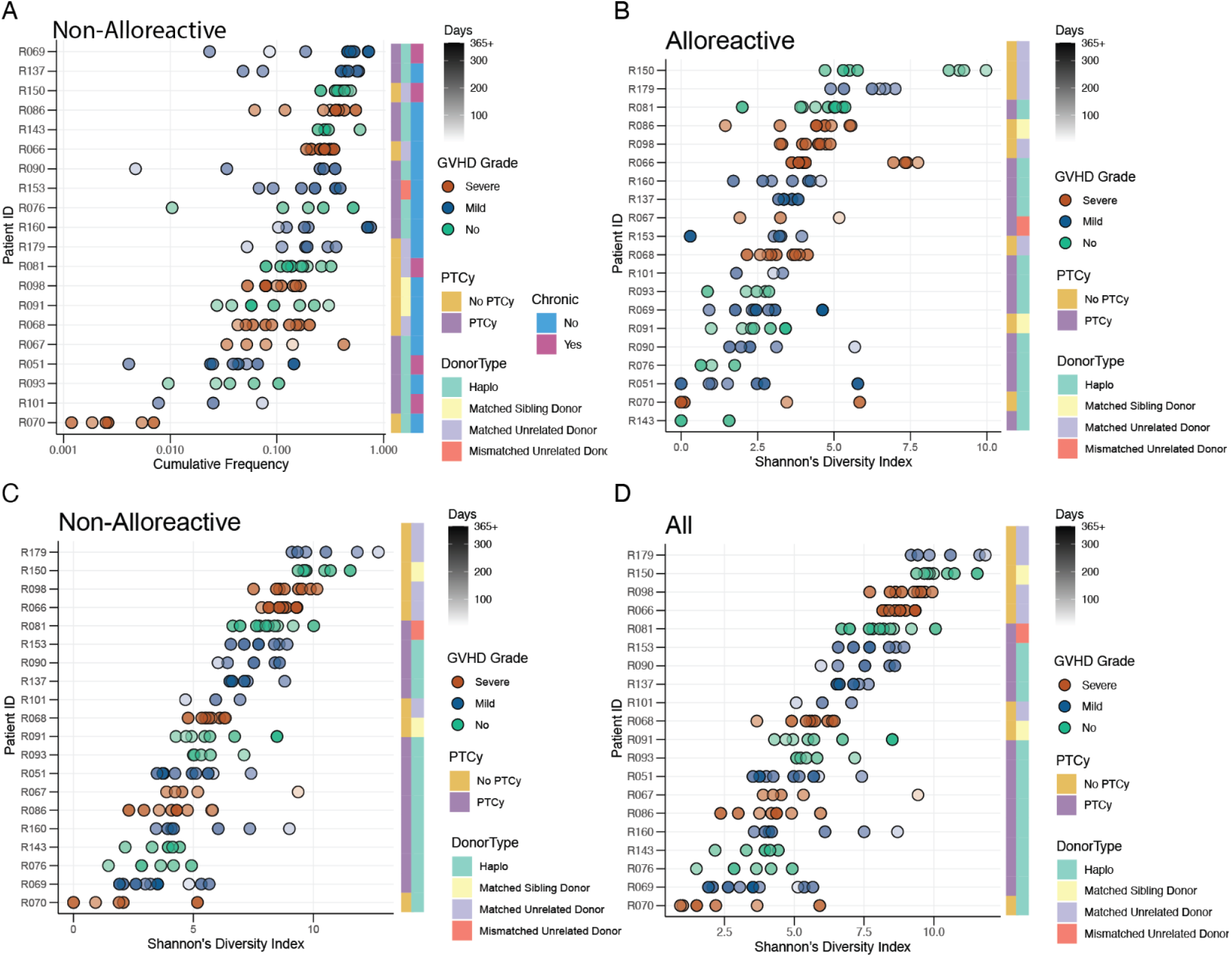
Overview of the time-series TCR repertoire data. **A**, **B**, **C**, **D**) show frequency or diversity for alloreactive, non-alloreactive, and all clones. Each patient is color-coded by their grade of GVHD, and the shade indicates the time point. Patients were ranked vertically based on the median value.

**Supplementary figure 2.**
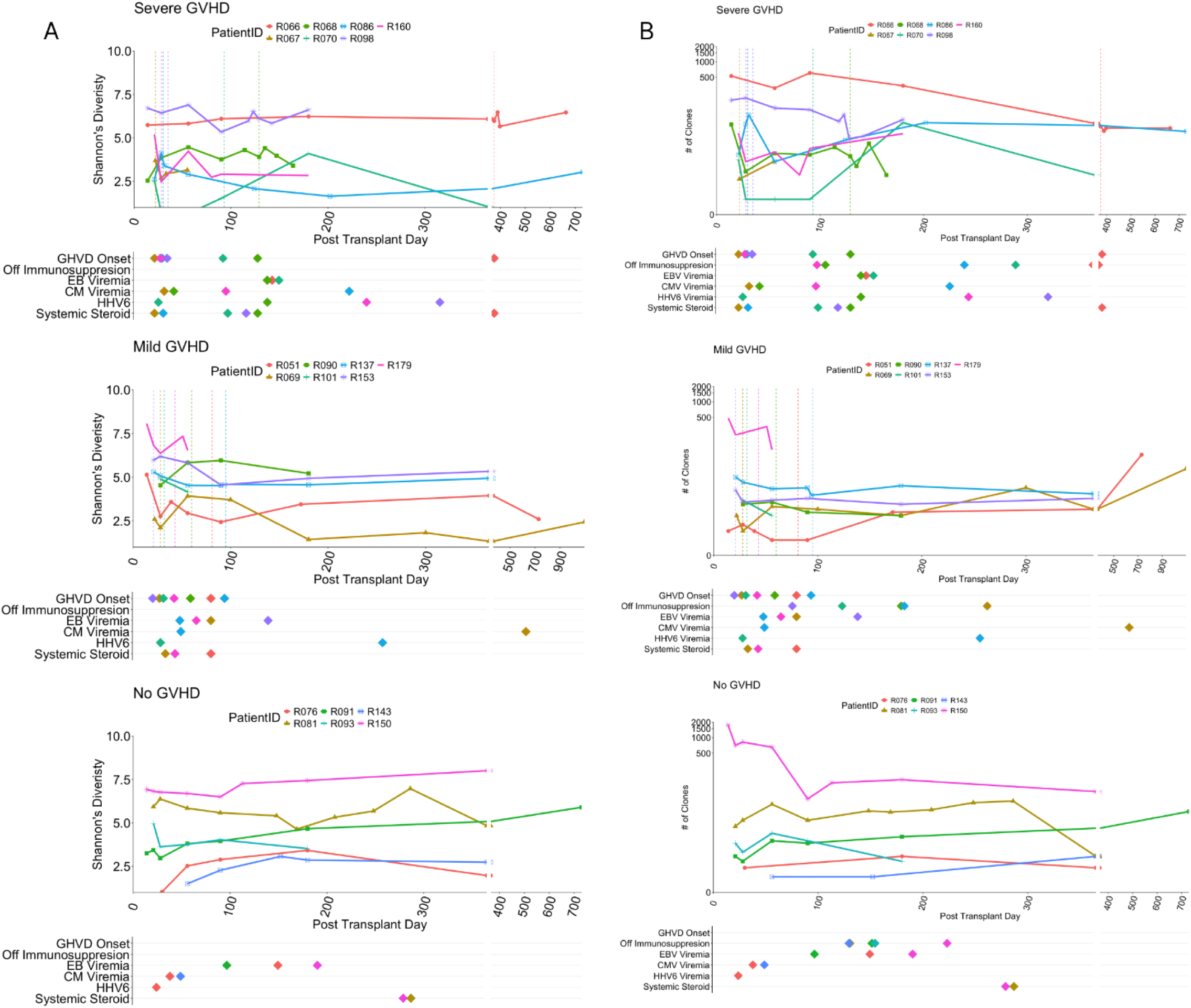
Overview of the time-series TCR repertoire data. Diversity (**A**) or number (**B**) of detectable alloreactive clones over time is shown for each patient. Matching clinical information is provided below each plot.

**Supplementary figure 3.**
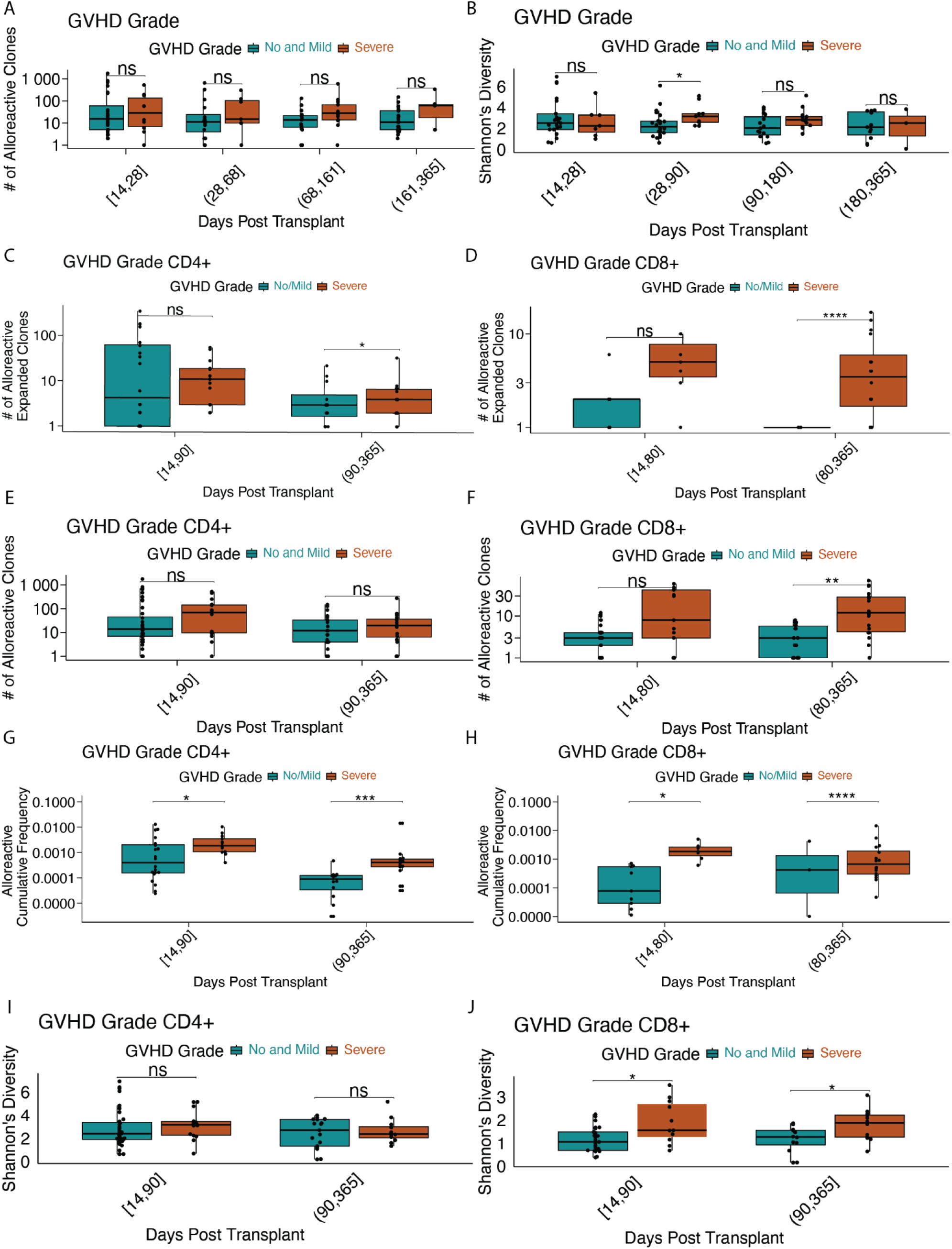
Comparison of alloreactive T-cell clones (including CD4^+^ and CD8^+^ subsets) between severe and no/mild GVHD across timepoints post-transplant. (**A**) Number of alloreactive clones across four post-transplant time windows in severe versus no/mild GVHD patients. (**B**) Shannon’s diversity index of alloreactive clones comparing severe and no/mild GVHD. (**C–D**) Number of expanded CD4⁺ (**C**) and CD8⁺ (**D**) alloreactive clones at early and late timepoints. (**E–F**) Total number of CD4⁺ (**E**) and CD8⁺ (**F**) alloreactive clones across timepoints. (**G–H**) Cumulative frequency of alloreactive CD4⁺ (**G**) and CD8⁺ (**H**) clones. (**I–J**) Shannon’s diversity index of CD4⁺ (**I**) and CD8⁺ (**J**) alloreactive clones at early and late timepoints. Statistical significance was assessed using the Wilcoxon test. *p<0.05; **p<0.01; ***p<0.001; ****p<0.0001; ns, not significant.

**Supplementary figure 4.**
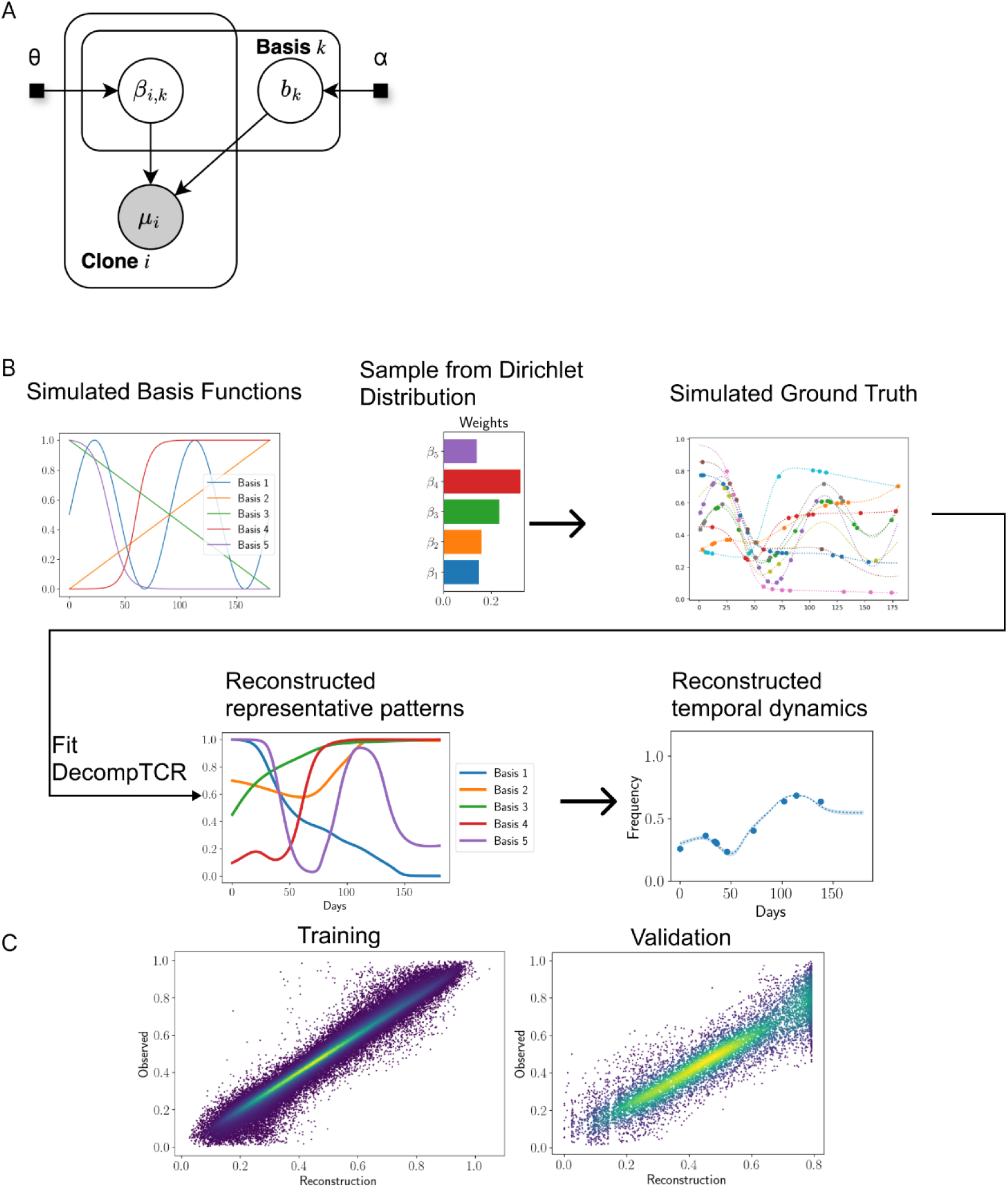
DecompTCR performance on simulated data. **A)** Plate model for DecompTCR. **B)** Simulation process and application of DecompTCR on simulated data. **C)** Scatter plots show the training and validation performance of DecompTCR.

**Supplementary figure 5.**
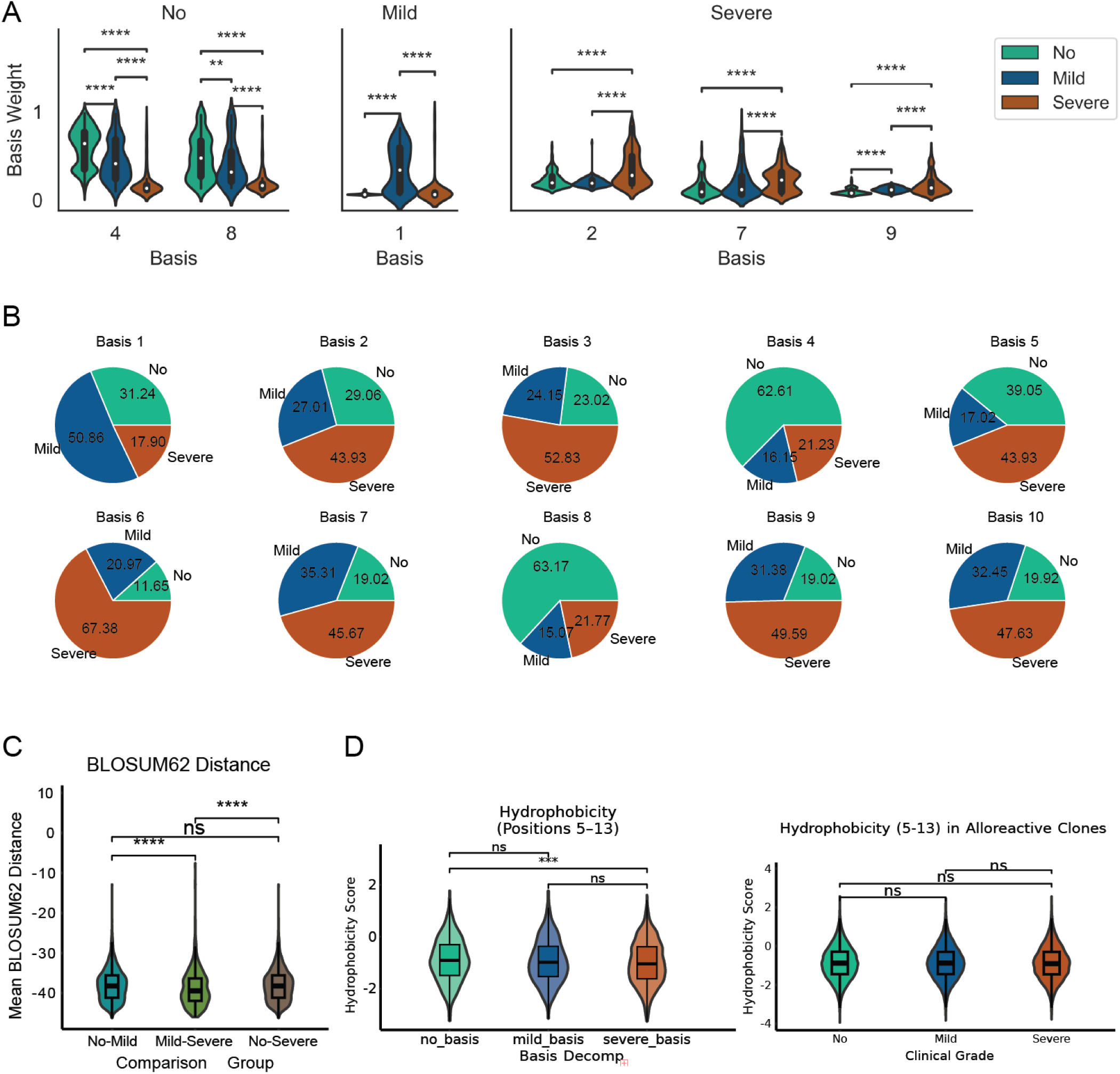
Assignment of basis to each grade of GVHD. **A)** Violin plots of basis weights for clones stratified by GVHD grades show statistically significant enrichment of specific patterns in different GVHD grades (Benjamini-Hochberg adjusted Wilcoxon rank-sum test). (**B**) Proportions of clones from each grade of GVHD in individual bases. (**C**) BLOSUM62 distance comparing different grades of GVHD. **(D**) Hydrophobicity of amino acid positions from 5-13, comparing clones representing different grades of GVHD with and without basis decomposition.

**Supplementary figure 6.**
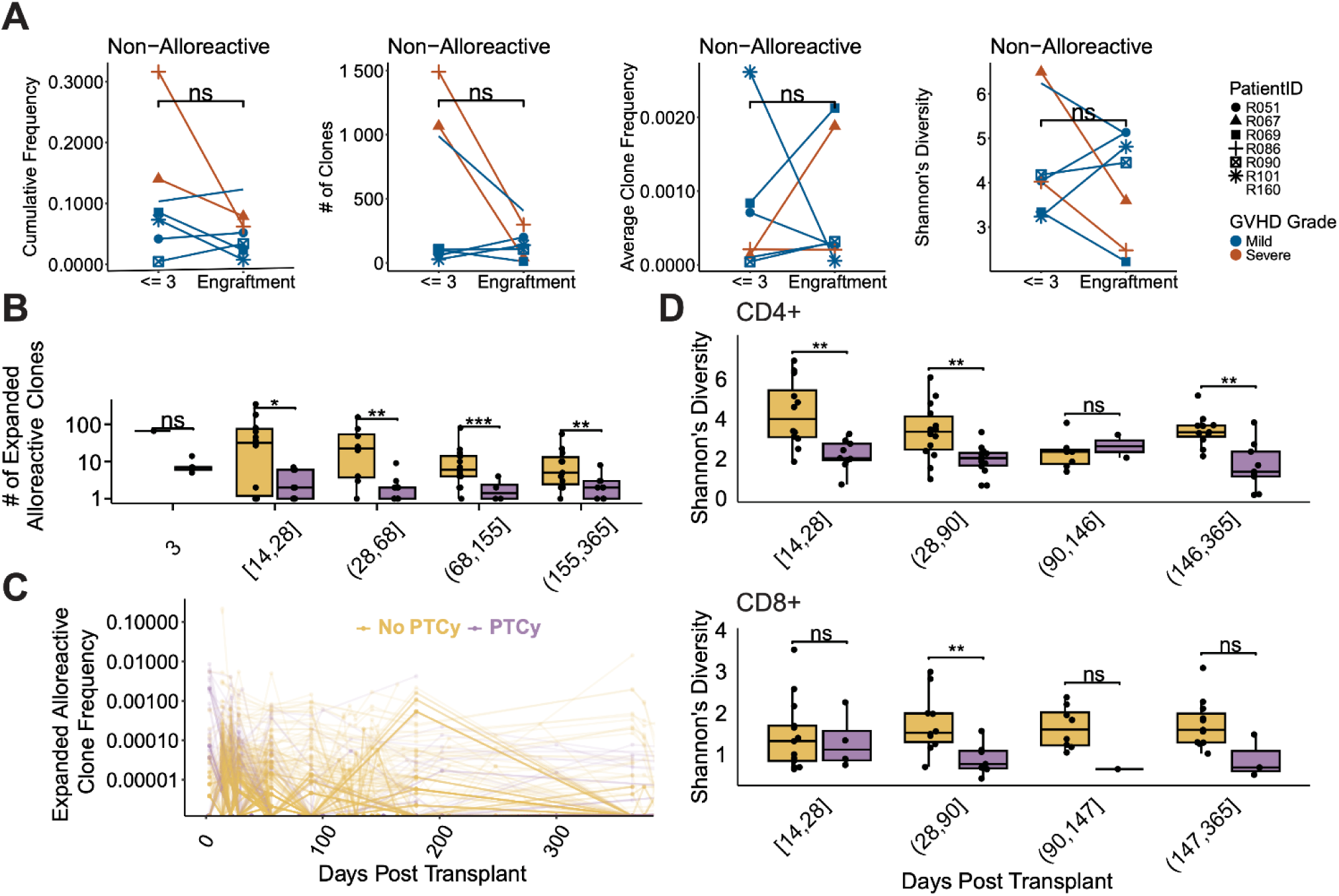
PTCy Selectively Depletes Expanding Alloreactive T Cell Clones. (**A**) Analysis of cumulative frequency, number of clones, average clone frequency, and Shannon’s diversity for non-alloreactive clones comparing D3 and the engraftment time point in PTCy-treated patients. (**B**, **C**) Temporal comparison of the number of expanded alloreactive clones and their corresponding individual clone frequency in PTCy-treated versus untreated patients across multiple time points (Benjamini-Hochberg adjusted Wilcoxon rank-sum test). (**D**) Temporal dynamics of Shannon’s diversity were analyzed for CD4 clones and CD8 clones, comparing PTCy-treated patients with untreated patients (Benjamini-Hochberg adjusted Wilcoxon rank-sum test, * P < 0.05, ** P < 0.01, *** P < 0.001, **** P < 0.0001).

**Supplementary figure 7.**
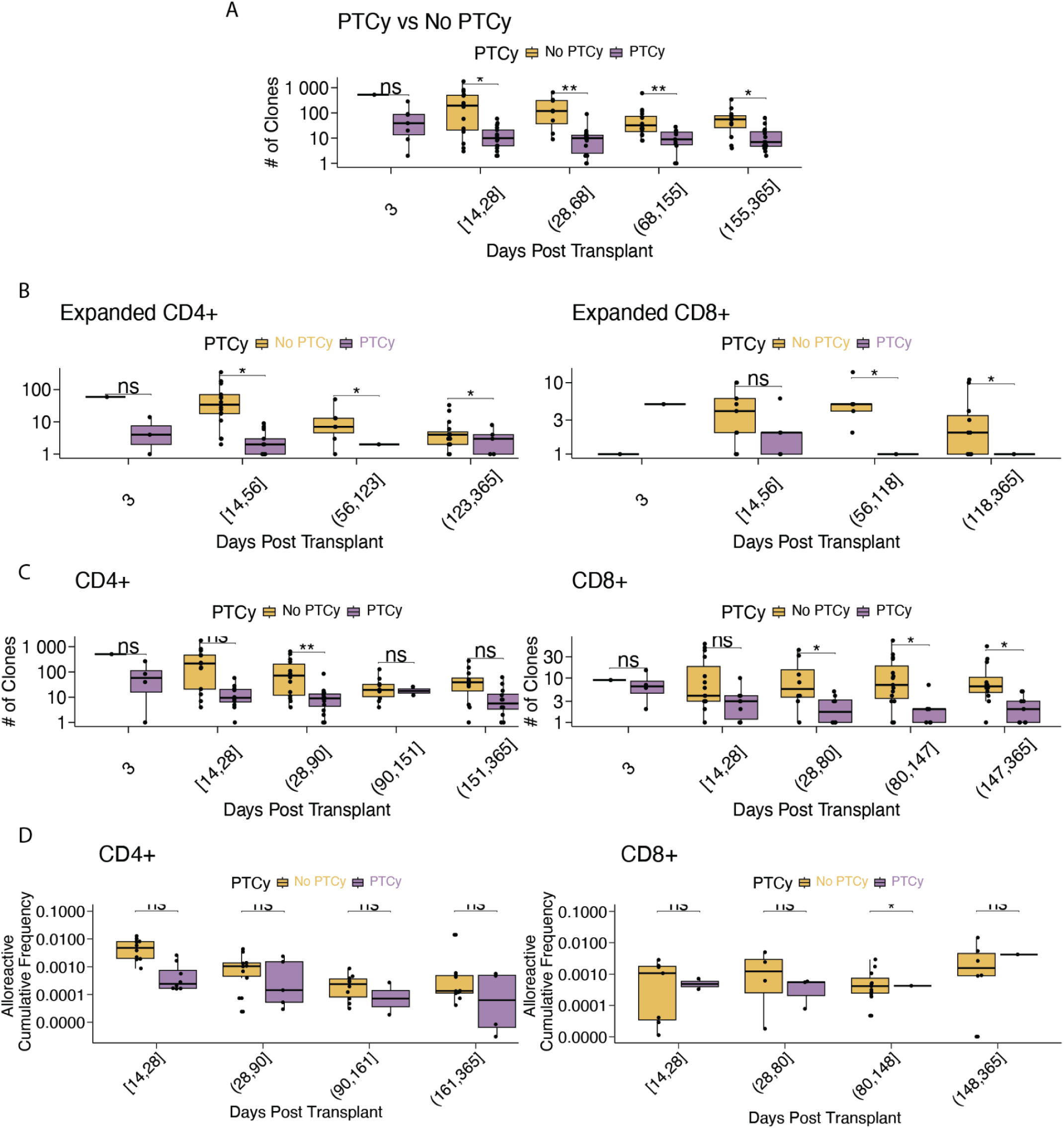
Impact of PTCy on alloreactive T-cell clones over time post-transplant. (**A**) Number of alloreactive clones across different time windows in patients with or without PTCy treatment. (**B**) Number of expanded CD4⁺ and CD8⁺ alloreactive clones at various time points. (**C**) Total number of CD4⁺ and CD8⁺ alloreactive clones in PTCy and no PTCy groups across time points. (**D**) Cumulative frequency of alloreactive CD4⁺ and CD8⁺ clones post-transplant. Statistical significance was determined by the Wilcoxon test. *p<0.05; **p<0.01; ***p<0.001; ns, not significant.

**Supplementary figure 8.**
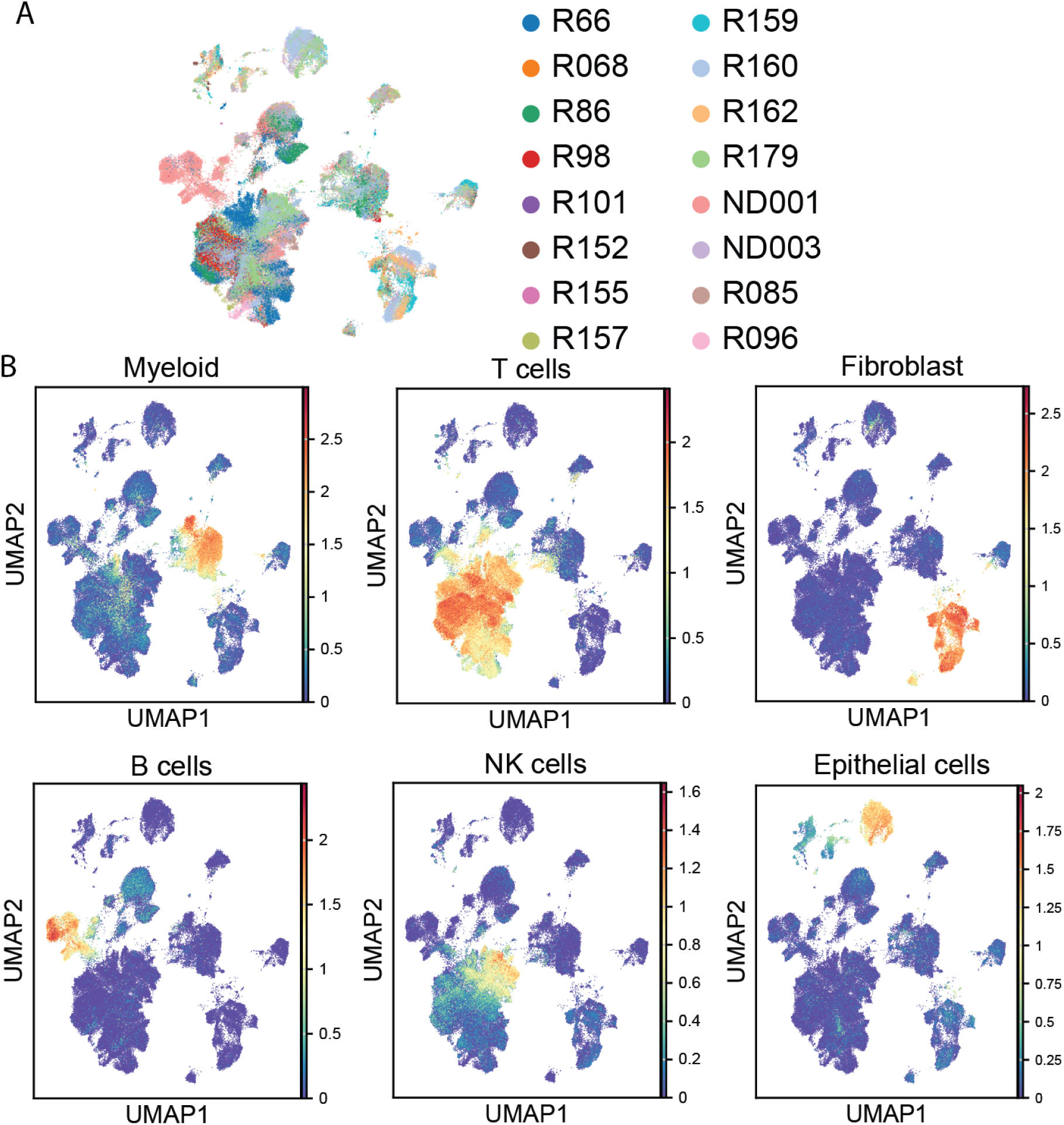
Cell type identification across all patients. (**A**) UMAP of single-cell RNA Seq data for all patients after batch correction. (**B**) Gene expression signatures that identify myeloid, T cells, fibroblasts, B cells, NK cells, and epithelial cells.

**Supplementary figure 9.**
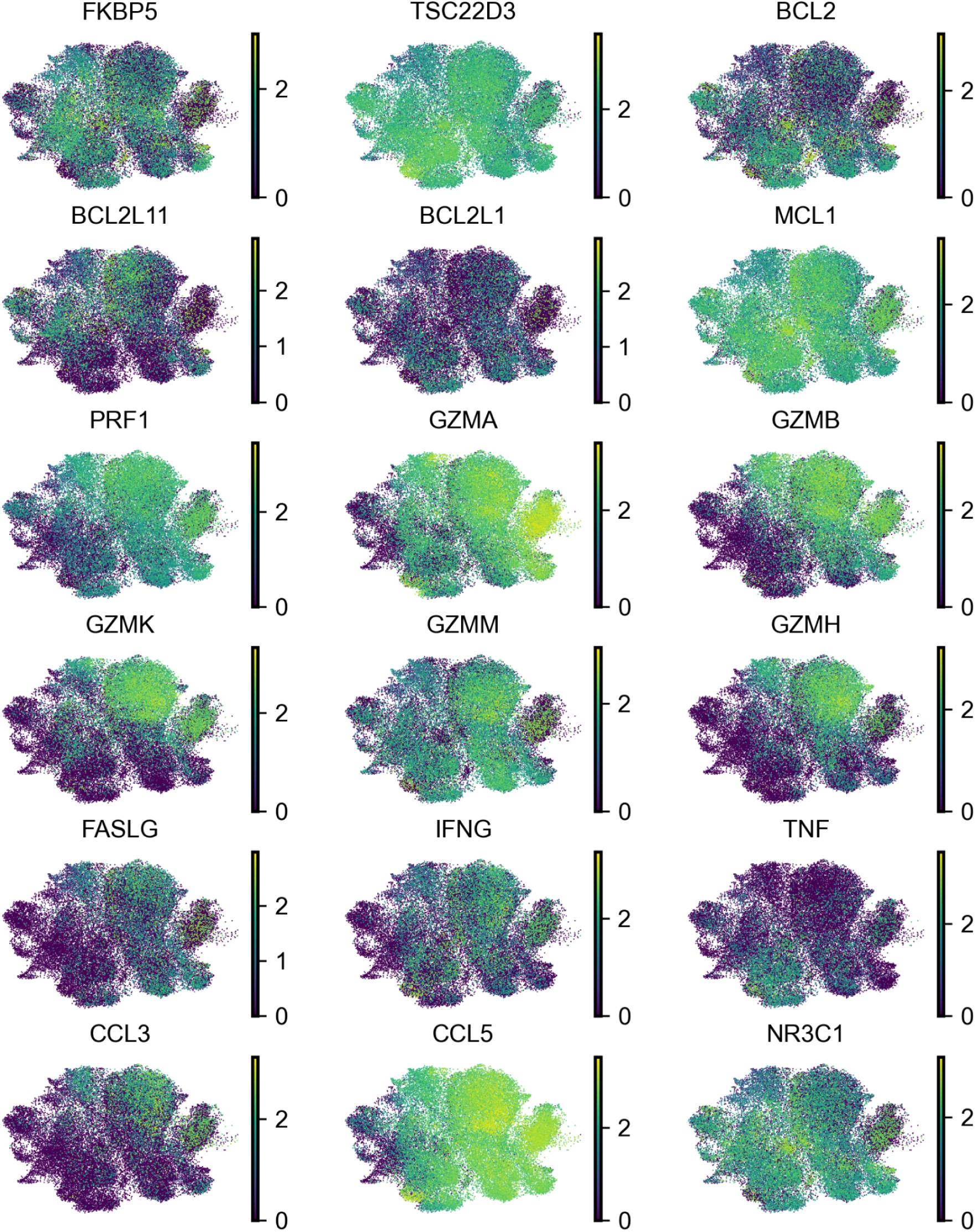
UMAP feature plots of select effector molecules, apoptosis markers and steroid resistance genes. Each panel shows the normalized expression of a selected gene projected onto the UMAP embedding of single-cell transcriptomes. Expression intensity is color-coded from low (purple) to high (yellow).

**Supplementary figure 10.**
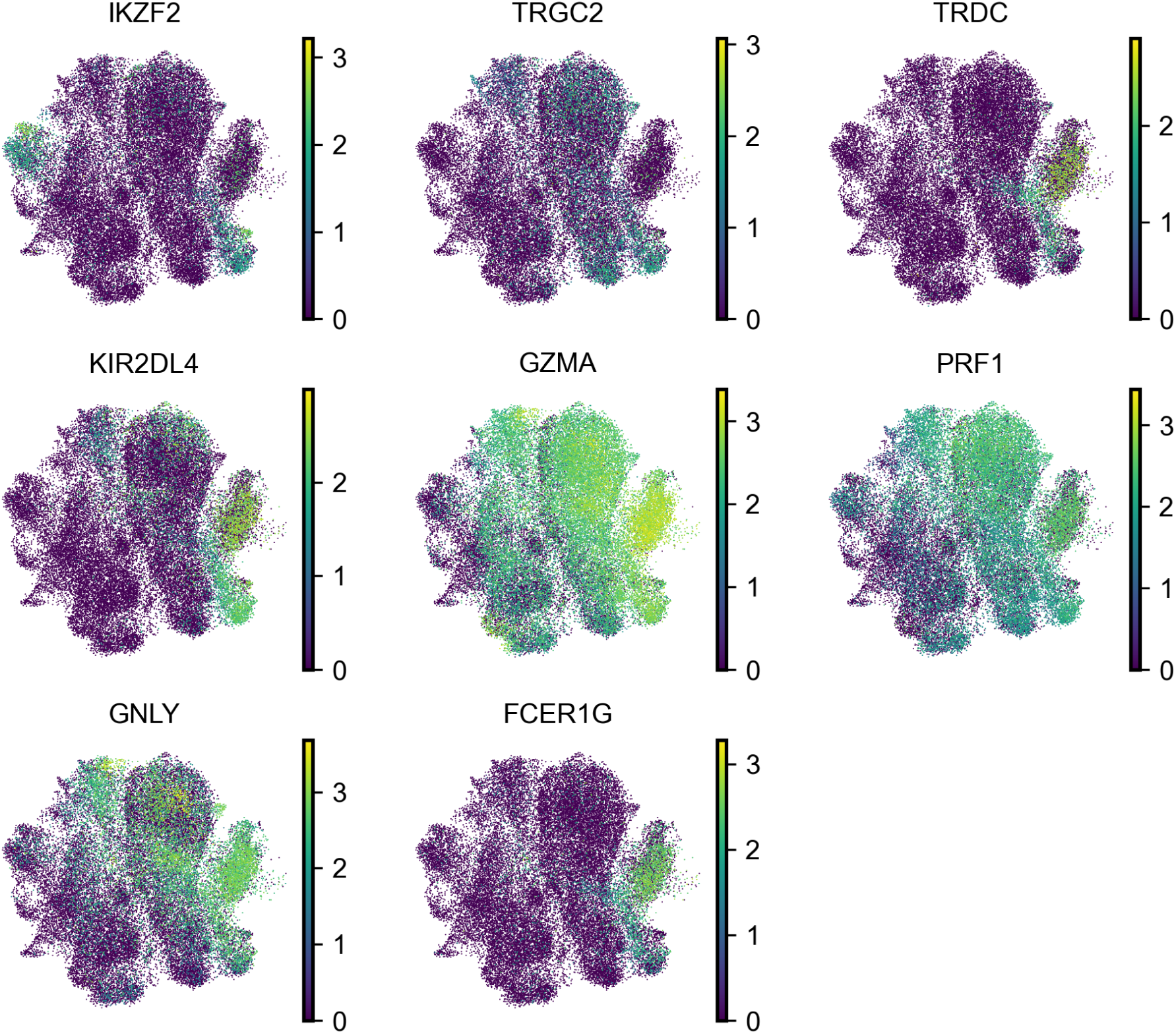
UMAP feature plots of unconventional T cell markers. Each panel shows the normalized expression of a selected gene projected onto the UMAP embedding of single-cell transcriptomes. Markers associated with cytotoxic T cells (e.g., *TRGC2*, *TRDC*, *KIR2DL4*, *GZMA*, *PRF1*, *GNLY*, *FCER1G*). Expression intensity is color-coded from low (purple) to high (yellow).

**Supplementary figure 11.**
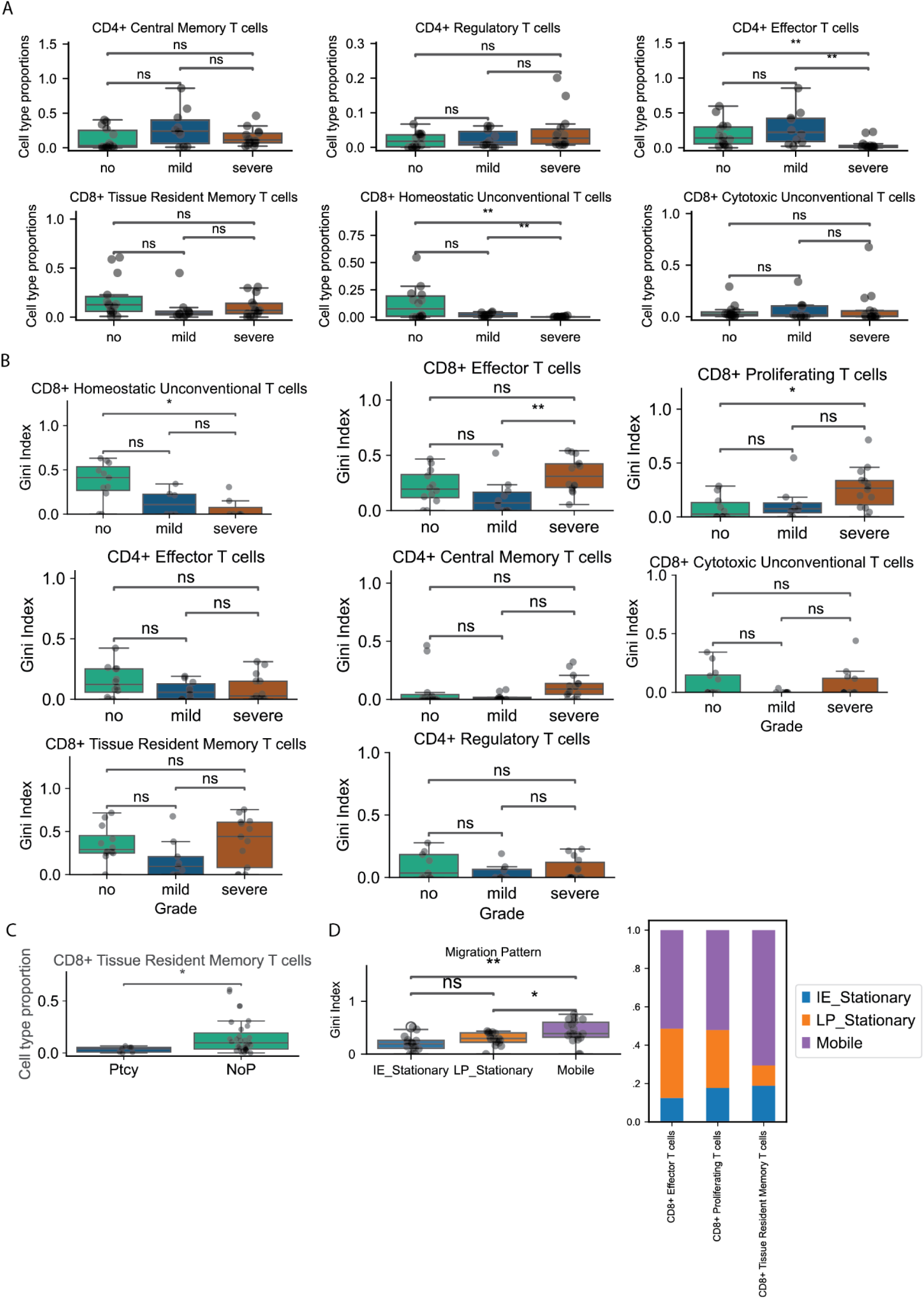
T cell proportions and clonal distribution in different GVHD grades. (**A**) Boxplots display cell type proportion across (remaining panels) various T cell subsets stratified by GVHD grade (no, mild, severe). (**B**) Boxplots display Gini indices indicating clonal inequality across various T cell subsets stratified by GVHD grade (no, mild, severe). Higher Gini indices represent lower diversity and greater clonal expansion, with statistical significance indicated. (**C**) Boxplots display cell type proportion comparing PTCy-treated patients vs untreated patients on CD8+ Tissue Resident Memory T cells. (*p<0.05, **p<0.01, ns=not significant, Mann Whitney with Holm multiple correction). (**D**) Boxplots display Gini indices indicating clonal inequality across different migration patterns (IE - Stationary, LP - Stationary, and Mobile), Bar plot for the mobile population on the mobile conventional population (chi-square, p = 5.4e-180).

**Supplementary figure 12.**
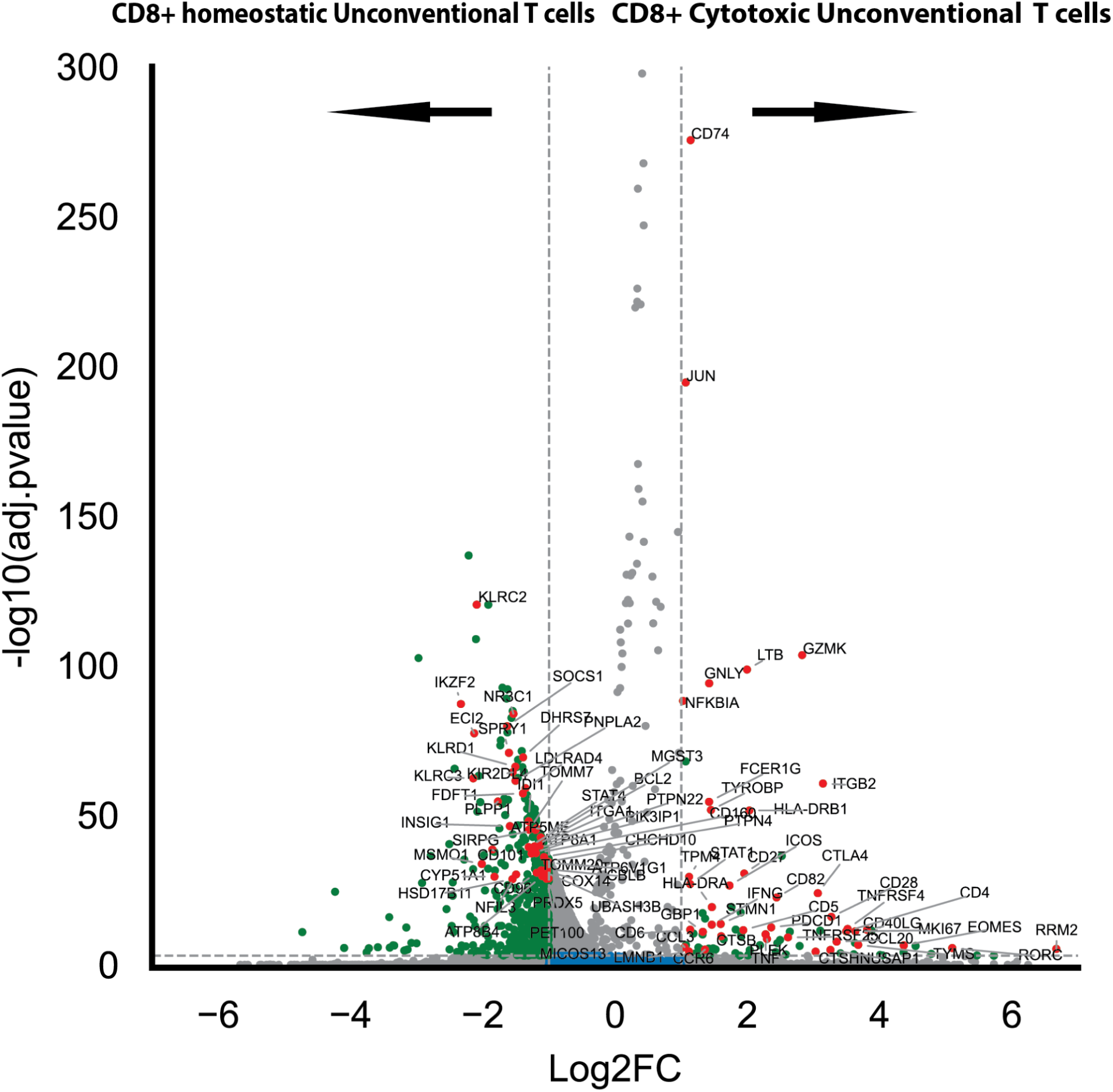
Volcano plots showing differentially expressed genes in CD8+ Unconventional cytotoxic compared to CD8+ Homeostatic Unconventional T cells (Wilcoxon, p <0.05, and log fold change > 0.5).

**Supplementary figure 13.**
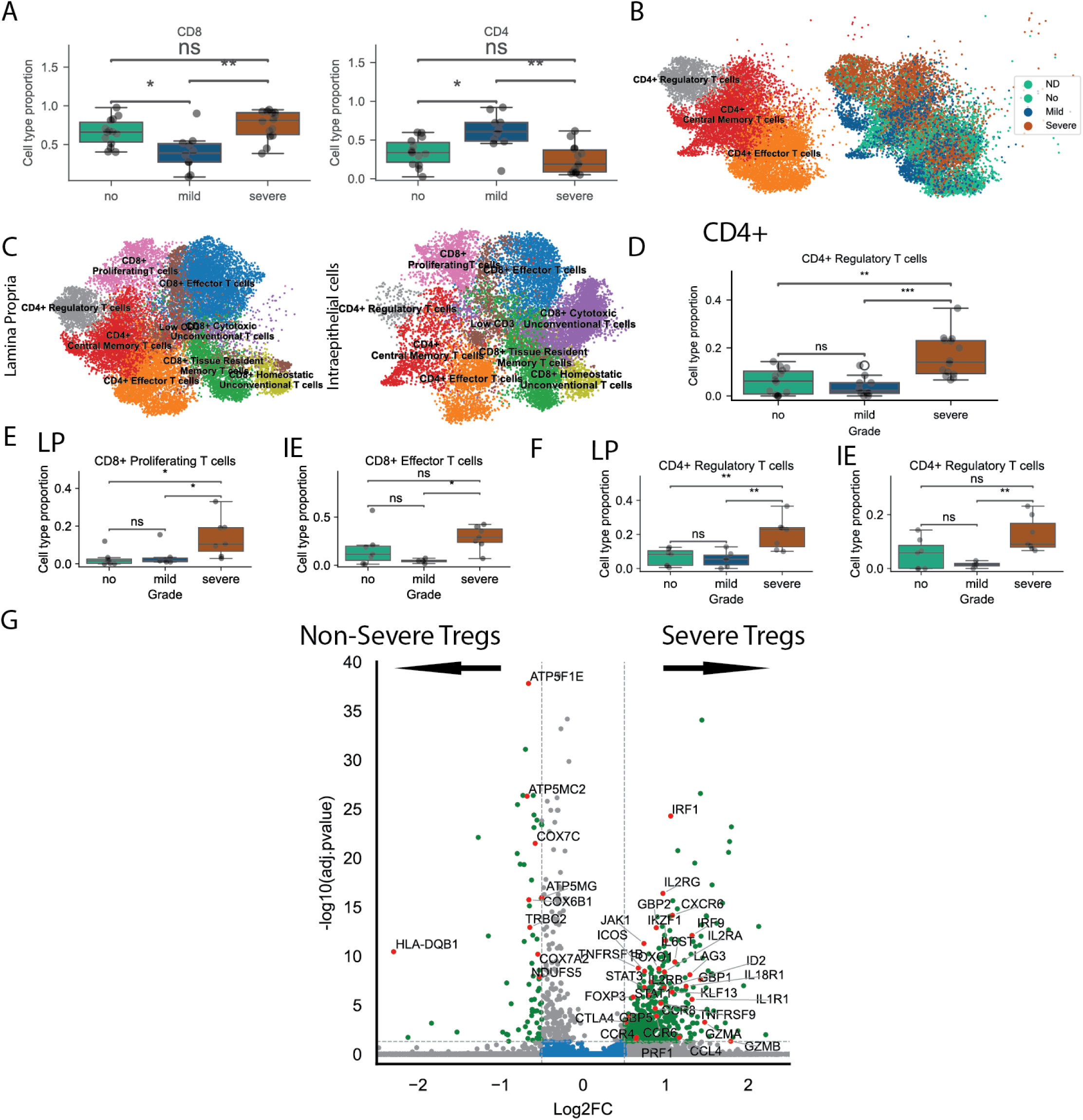
CD4⁺ Regulatory T cell enrichment and transcriptional profile in severe GVHD. (**A**) Boxplots showing the proportions of CD4⁺ and CD8⁺ T cells across GVHD grades (no, mild, severe). (**B**) UMAP visualization of CD4^+^ T cells colored by GVHD grade. (**C**) Annotated UMAP of T cell subtypes across lamina propria (LP) and intraepithelial (IE) compartments. (**D**) The proportion of CD4⁺ Regulatory T cells increases significantly in severe GVHD. (**E**) Subcompartment analysis of proliferating CD8⁺ T cells and CD8⁺ effector T cells in LP or IE. (**F**) Subcompartment analysis of CD4⁺ Regulatory T cells in CD4^+^ T cells in LP or IE. (**G**) Volcano plots showing differentially expressed genes in severe CD4⁺ Regulatory T cells compared to other Tregs (Wilcoxon, p <0.05, and log fold change > 0.5).

**Supplementary Figure 14.**
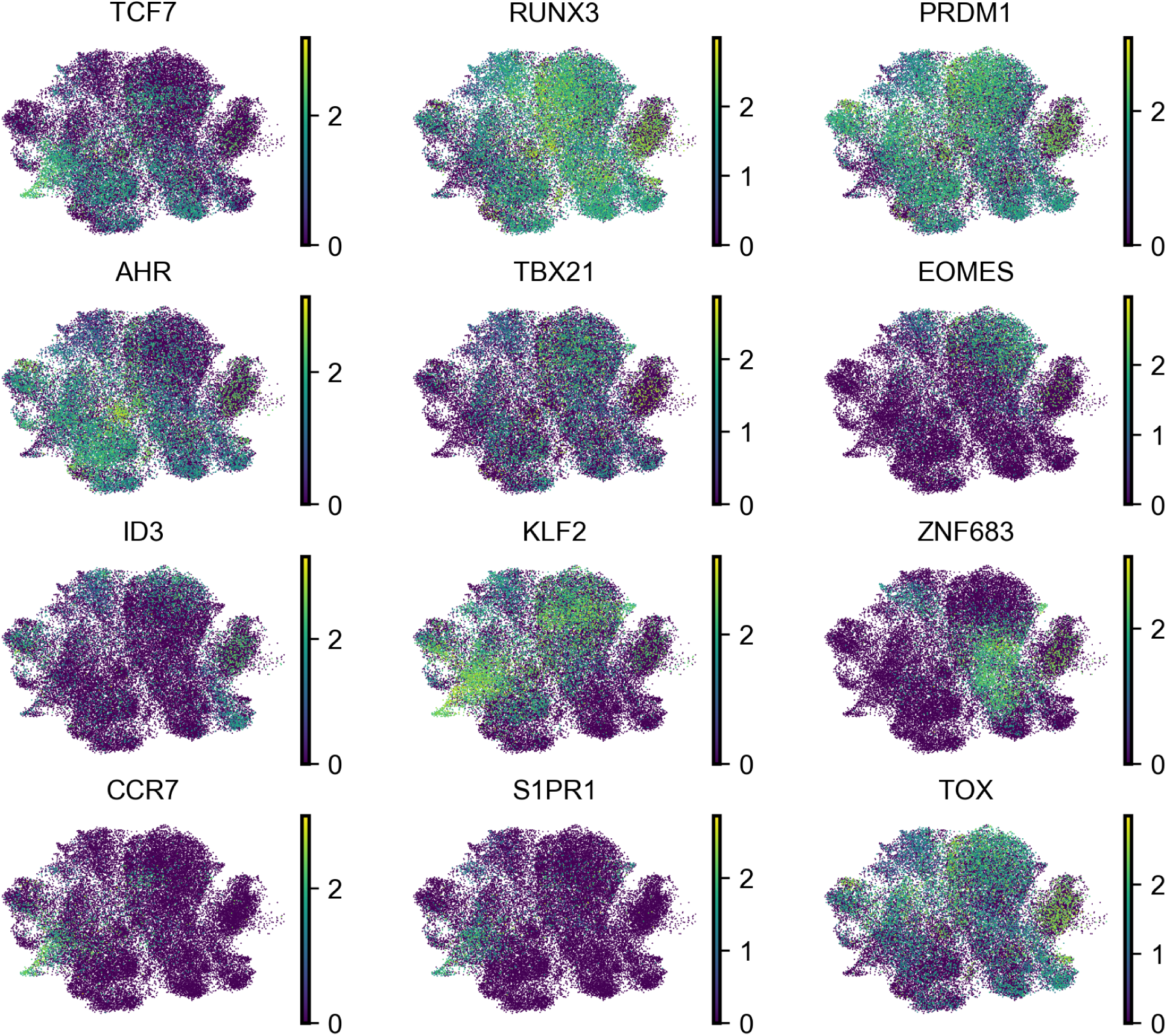
Expression of key transcription factors and migration markers across T cell subsets. UMAP plots display normalized expression levels (log-transformed) of genes associated with T cell differentiation and residency programs, including transcription factors such as *TCF7*, *RUNX3*, *PRDM1*, *TBX21*, *EOMES*, *ZNF683*, and regulators of migration like *CCR7*, *S1PR1*, and *KLF2*.

**Supplementary figure 15.**
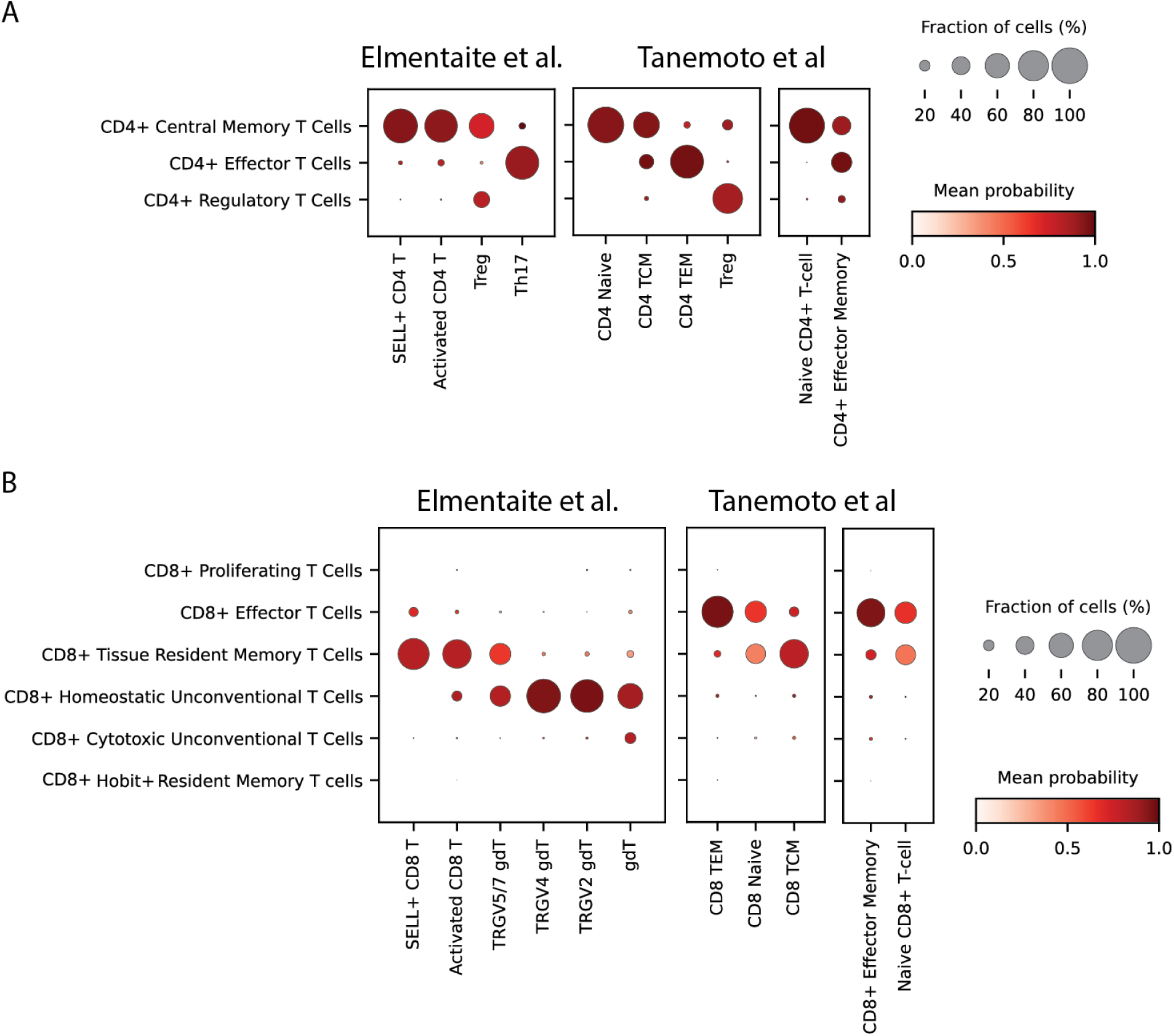
Cross-dataset mapping of T cell subsets using Celltypist predictions. (**A**) CD4+ T cell subsets and (**B**) CD8+ T cell subsets from our dataset were mapped onto reference cell types annotated in the Elementaite et al.^37^ and Tanemoto et al.^38^ datasets. The size of each dot represents the fraction of cells assigned to each reference population, and the color intensity reflects the mean prediction probability. Stronger red color indicates higher confidence in the assignment.

**Supplementary figure 16.**
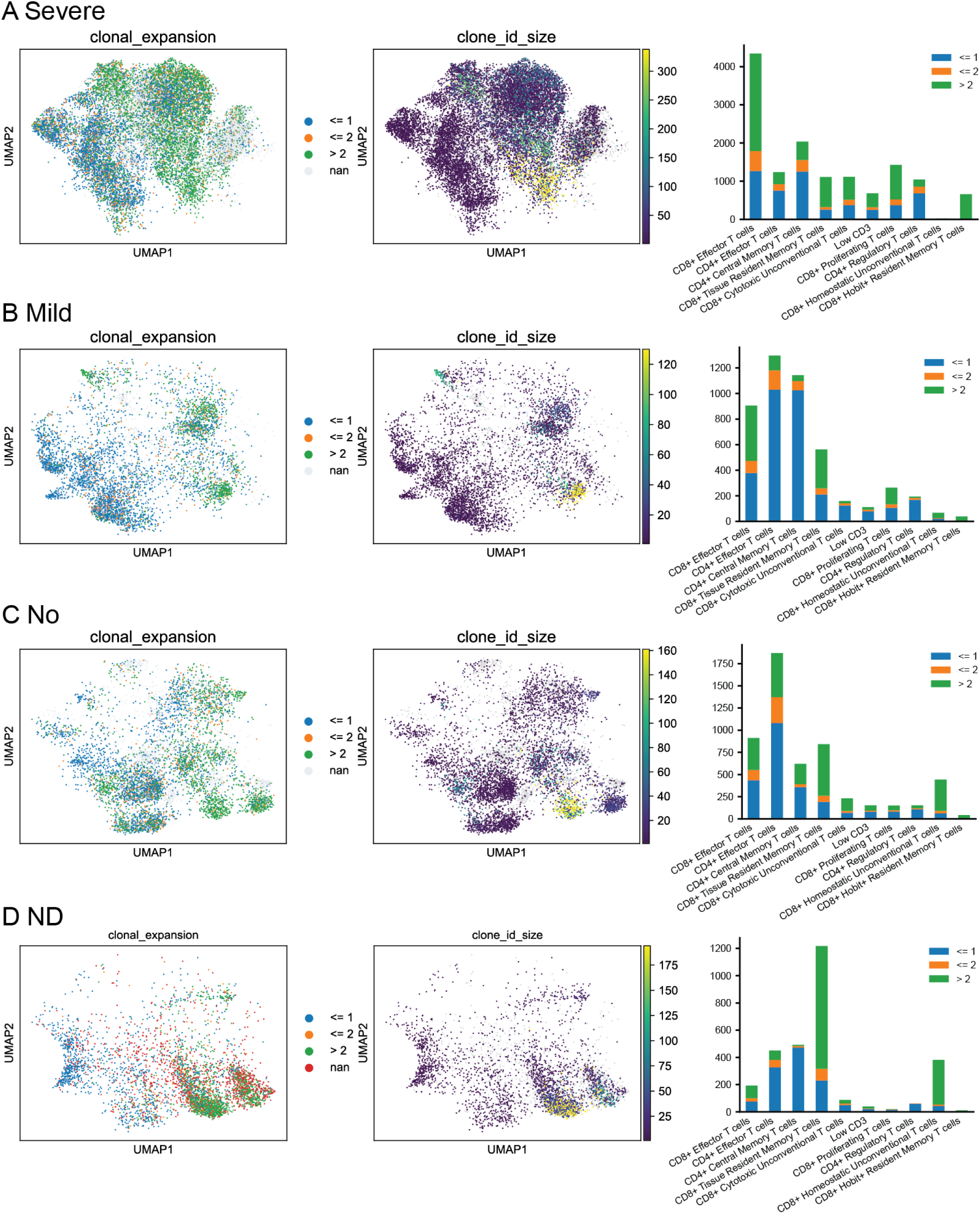
T Cell Clonal Expansion and Abundance Across GVHD Severity Levels. (**A, B, C, D**) Clone Size Distribution and Non-Normalized Clonal Abundance in severe, mild, no GVHD, and ND Groups.

**Supplementary figure 17.**
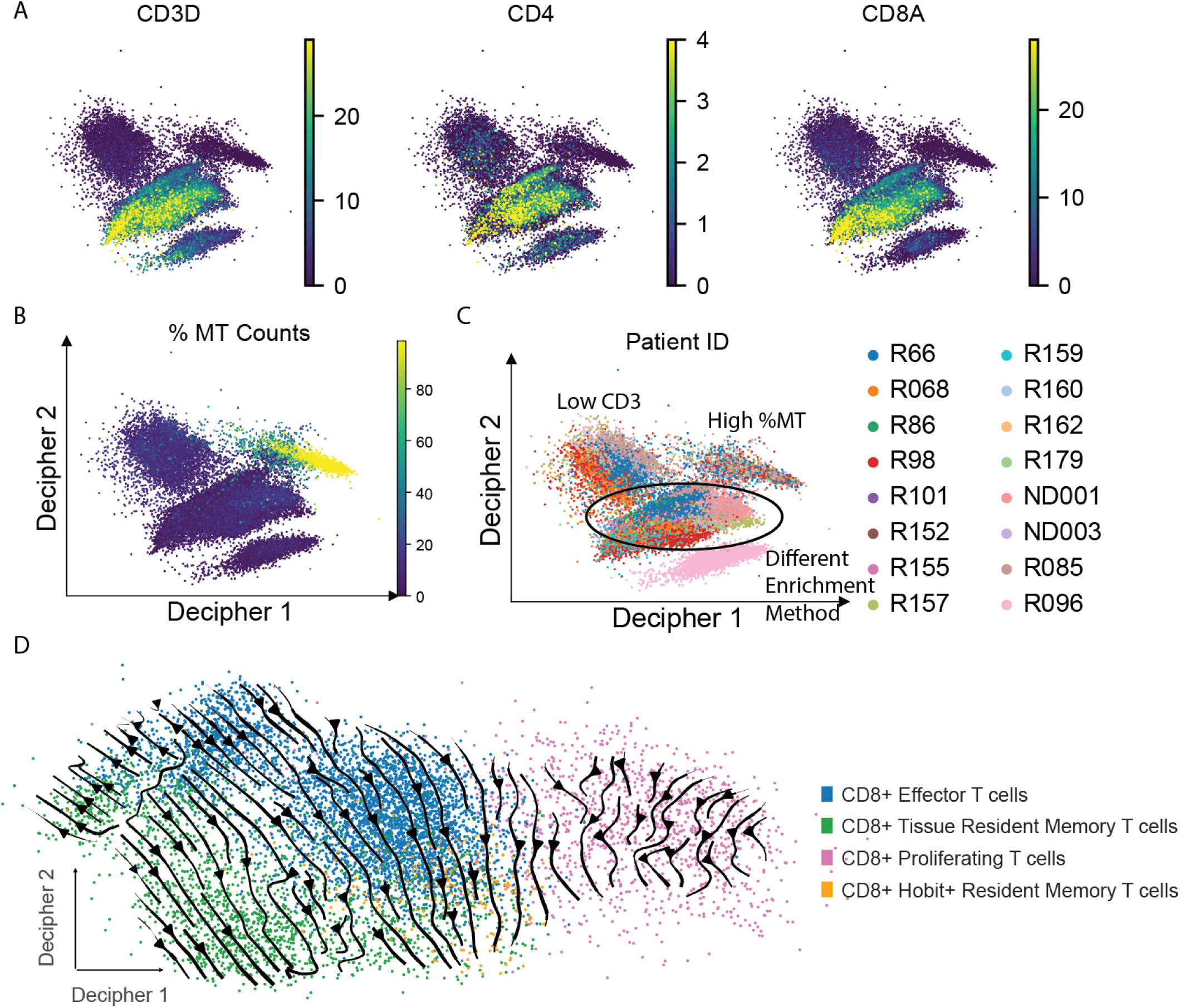
Decipher analysis separating cells with high mitochondrial counts and low CD3 cells. (**A**) CD3D, CD4, and CD8A expression in decipher space on all T cells. (**B**, **C**) % MT counts and Patient ID from all T cells on decipher space. (**D**) Decipher analysis projecting CD8^+^ T cell populations based on Decipher 1 and Decipher 2 axes, and the arrow is drawn from LP to IE among shared clones whose clone size is smaller than 20.

**Supplementary figure 18.**
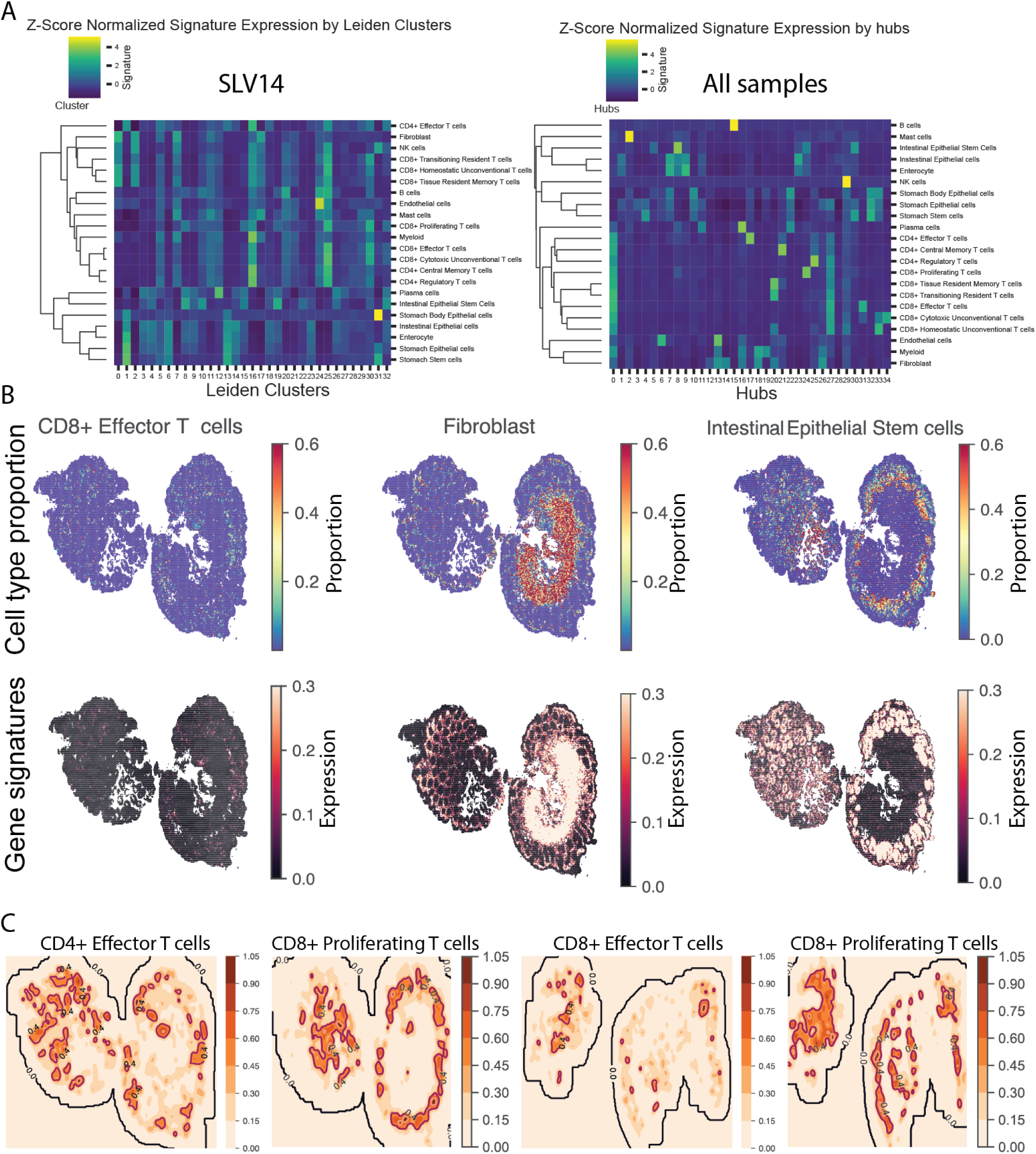
Cell type signatures across hubs and leiden clusters in intestinal tissues. (**A**) Heatmaps showing Z-score normalized expression of cell type-specific gene signatures by spatial hubs (left, sample SLV24) and by Leiden clusters (right, from all samples). Each row represents a cell type signature, and each column corresponds to a hub or cluster. Clusters reflect mixed signals from diverse cell types and immune cell states cannot be disentangled. Fine-grain cell states can be better distinguished in hubs identified by StarfyshHD. (**B**) Cell type proportions and gene signatures in tissue sections. The top row shows cell type proportion (CD8⁺ Effector T cells, Fibroblasts, Intestinal Epithelial Stem cells) across tissue sections from SLV1. The bottom row displays gene signature expression for the same cell types. (**C**) Contour plot of cell type proportions for SLV14 and SLV16.

**Supplementary figure 19.**
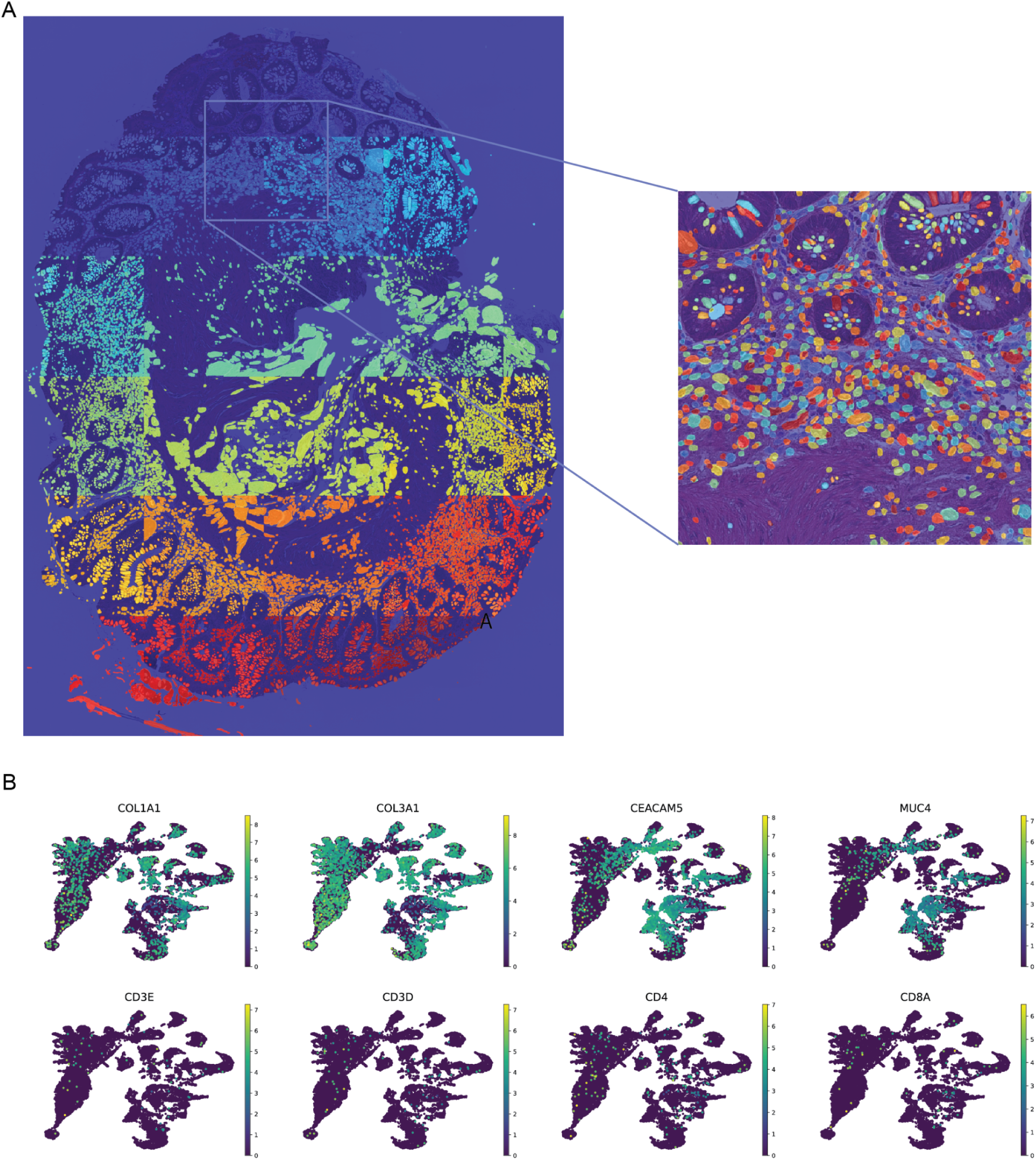
Single cell segmentation and transcriptional distribution of T cell markers in intestinal tissue. (**A**) CellSAM segmentation mask overlaid with the histology sample SLV14, highlighting cell-level boundaries across distinct anatomical regions. Expanded example patch (right) shows undersegmentation of cell types with diverse morphology in gut tissue. (**B**) UMAP projections of transcriptomes quantified according to segmented masks (each dot represents a cell mask) reveal discrete clusters corresponding to fibroblast (COL1A1, COL3A1) and epithelial (CEACAM5, MUC4) gene signatures. In contrast, canonical T cell markers (CD3E, CD3D, CD4, CD8A) appear broadly mixed with other cell types and do not form distinct clusters. These findings motivate the need for transcriptomic deconvolution of cell states, even with high-resolution Visium HD data.

**Supplementary figure 20.**
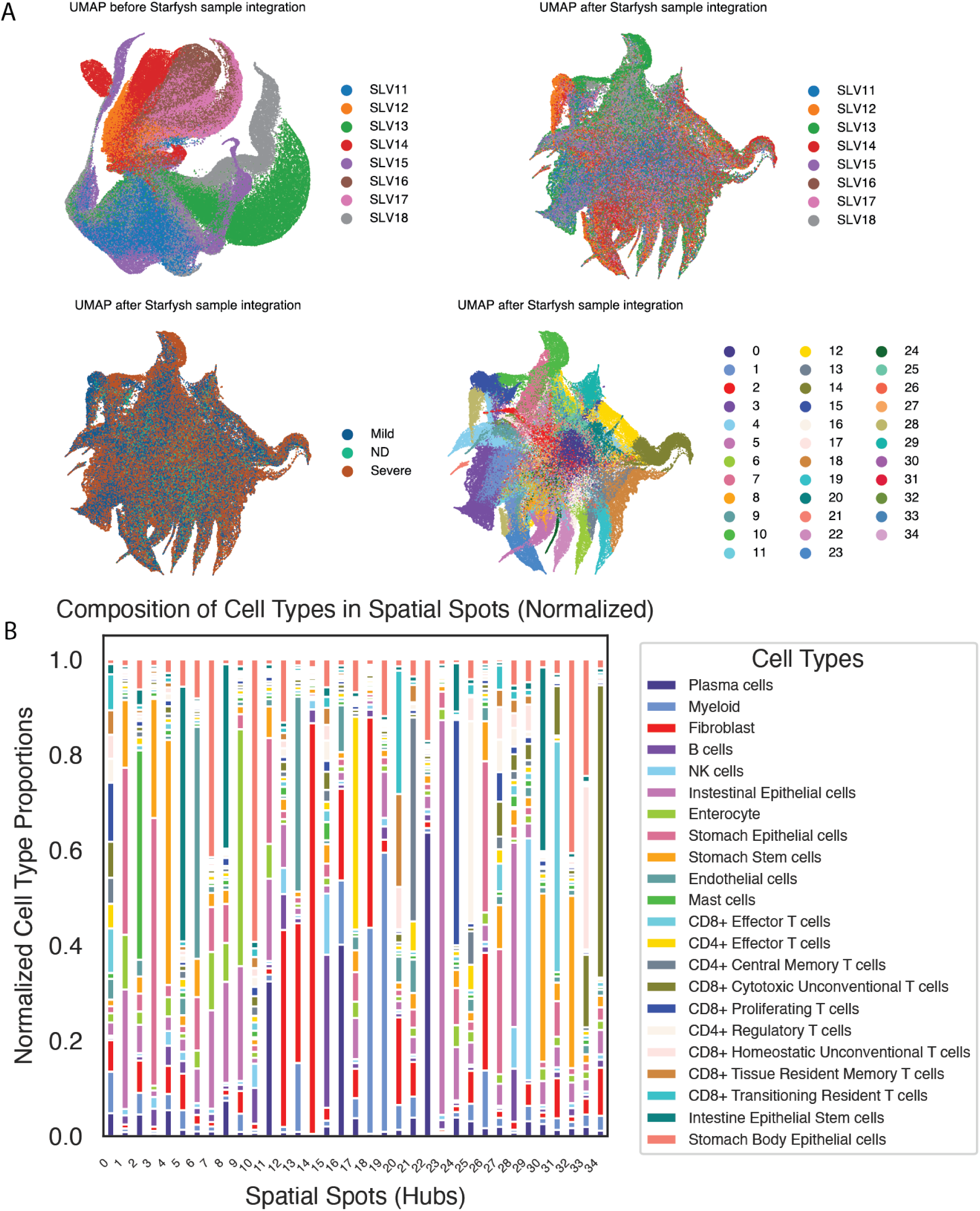
StarfyshHD Integration Improves Spatial and Cell Type Alignment Across Samples. (**A**) UMAP visualization of spatial transcriptomics data before and after sample integration with Starfysh. Each point represents a spatial spot, colored by sample before batch correction (top left), sample after batch correction (top right), GVHD severity (bottom left), and spatial hub identity (bottom right) after integration. (**B**) Normalized composition of cell types across spatial hubs after integration.

**Supplementary figure 21.**
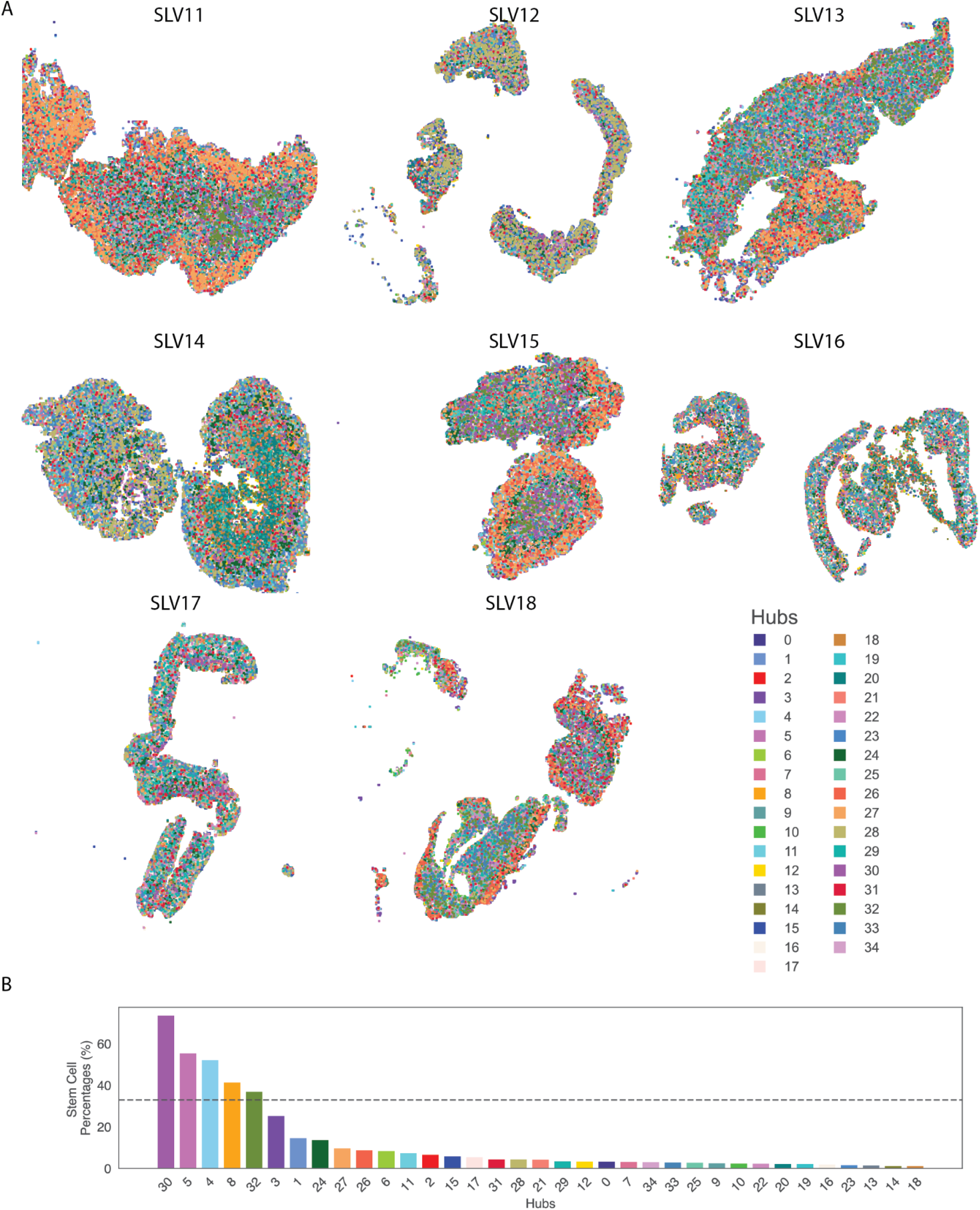
Integrated characterization of spatial hubs across intestinal samples and their association with stem cell enrichment. (**A**) spatial plots of tissues with spots colored by spatial hub ID inferred across eight gut samples (SLV11 to SLV18). (**B**) Bar plot depicting the percentage of stem cells found in each hub. Hubs are ranked by stem cell percentage. The dashed line represents the filtering threshold to identify stem cell-enriched hubs.

**Supplementary figure 22.**
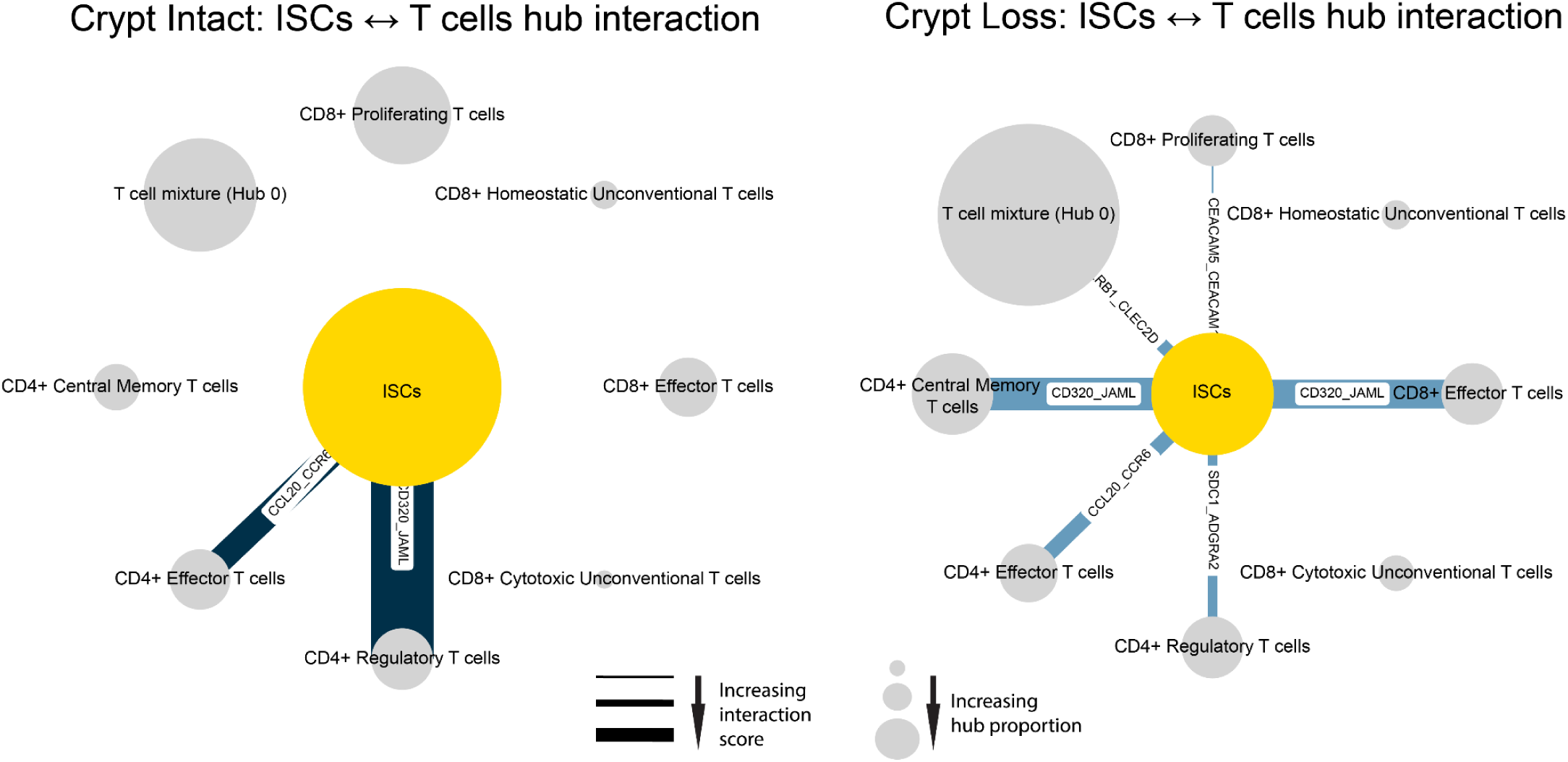
Intestinal Stem Cell (ISC)–T cell hub interaction networks differ between intact and crypt-loss regions. Network diagrams show ligand–receptor interactions predicted between ISC hubs (yellow) and various T cell subset hubs (gray nodes) in crypt intact (left) and crypt loss (right) regions. Edge thickness represents interaction strength, and labels denote specific ligand–receptor pairs.

## SUPPLEMENTARY INFORMATION

### KEY RESOURCES TABLE

| REAGENT or RESOURCE | SOURCE | IDENTIFIER |
| --- | --- | --- |
| **Antibodies** |  |  |
| Anti-human CD19 | BD Biosciences | Cat#561295; Clone HIB19 |
| Anti-human CD14 | BD Biosciences | Cat#562693; Clone MΦP-9 |
| Anti-human CD56 | BD Biosciences | Cat#560842; Clone B159 |
| Anti-human CD3 | BD Biosciences | Cat#563275; Clone UCHT-1 |
| Anti-human CD4 | BD Biosciences | Cat#560769; Clone RPA-T4 |
| Anti-human CD8 | BD Biosciences | Cat#560273; Clone SK1 |
| Anti-human CD45 | BD Biosciences | Cat#555480; Clone HI30 |
| Cell Viability<br>Solution(7-Aminoactinomycin D) | BD Biosciences | Cat#555815 |
| **Biological Samples** |  |  |
| Human peripheral blood | Columbia University<br>Medical Center | N/A |
| Human gastrointestinal biopsies | Columbia University<br>Medical Center | N/A |
| **Chemicals, Peptides, and**<br>**Recombinant Proteins** |  |  |
| Human GM-CSF Recombinant<br>Protein | Fisher Scientific | Cat#215GM010CF |
| Human IL-4 Recombinant Protein | Fisher Scientific | Cat#204IL010CF |
| TNF-α, Human | Millipore Sigma | Cat#SRP3177-50UG |
| Prostaglandin E2 | Fisher Scientific | Cat#22-961-0 |
| Dithiothreitol (DTT) | Sigma-Aldrich | Cat# 43816 |
| DNase | Sigma-Aldrich | Cat#11284932001 |
| Collagenase IV | Gibco | Cat#17104019 |
| 0.5M EDTA | Invitrogen | Cat#AM9260G |
| HEPES Buffer Solution (1M) | Gibco | Cat#15630080 |
| RPMI 1640 Medium | Gibco | Cat#11875119 |
| Human Serum AB | GeminiBio Product | Cat#100-512-100 |
| Gentamicin | Gibco | Cat#15710072 |
| L-Glutamine | GeminiBio Product | Cat#400-106 |
| Phosphate-buffered saline (PBS, 1X), sterile-filtered | ThermoScientific | Cat#J61196AP |
| HBSS [Ca<sup>2+</sup> & Mg<sup>2+</sup> free] | Gibco | Cat#14170161 |
| **Critical Commercial Assays** |  |  |
| CellTrace CFSE Proliferation Kit | Thermo Fisher Scientific | Cat#C34554 |
| CellTrace Far Red Cell Proliferation Kit | Thermo Fisher Scientific | Cat#C34564 |
| Via-Probe Cell Viability Solution | BD Biosciences | Cat#555815 |
| CD14 MicroBeads, Human | Miltenyi Biotec | Cat#130-050-201 |
| CD45 MicroBeads, Human | Miltenyi Biotec | Cat#130-045-801 |
| QIAamp DNA Mini Kit | Qiagen | Cat#51304 |
| DNeasy Blood & Tissue Kit | Qiagen | Cat#69504 |
| 10X Genomics Chromium Single Cell 5' Kit | 10X Genomics | PN-1000263 |
| 10X VisiumHD WT Panel Gene Expression | 10X Genomics | PN -1000675 |
| immunoSEQ TCRB sequencing | Adaptive Biotechnologies | N/A |
| **Deposited Data** |  |  |
| TCR sequencing data | This paper | Will be deposited to GEO upon publication. |
| Single-cell RNA-seq data | This paper | Will be deposited to GEO upon publication. |
| Spatial RNA-Seq data | This paper | Will be deposited to GEO upon publication. |
| **Software and Algorithms** |  |  |
| immunoSEQ Analyzer | Adaptive Biotechnologies | https://www.immunoseq.com |
| cdr3tools | Obradovic et al., 2021 | https://github.com/caparks2/cdr3tools |
| scanpy | Wolf et al., 2018 | https://scanpy.readthedocs.io/en/stable/# |
| scVI | Lopez et al., 2018 | https://scvi-tools.org |
| Decipher | Nazaret et al., 2023 | https://github.com/azizilab/decipher |
| Phenograph | DiGiuseppe et al., 2019 | https://dpeerlab.github.io/dpeerlab-website/phenograph.html |
| scrublet | Wolock et al., 2019 | https://github.com/swolock/scrublet |
| DecompTCR | This paper | https://github.com/azizilab/GVHD_Reproducibility |
| DecipherTCR | This paper | https://github.com/azizilab/GVHD_Reproducibility |
| StarfyshHD | This paper | https://github.com/azizilab/GVHD_Reproducibility |
| **Other** |  |  |
| gentleMACS Dissociator | Miltenyi Biotec | N/A |
| BD Influx Cell Sorter | BD Biosciences | N/A |

### EXPERIMENTAL MODEL AND SUBJECT DETAILS

#### Human subjects

Pre-transplant recipient and donor blood samples, post-transplant PBMCs, and tissue biopsies were collected at Columbia University Irving Medical Center following approval by the Institutional Review Board (Protocol AAAF2395). Informed consent was obtained from all participants. Samples from unrelated donors were received from the donor centers using a protocol approved by the Institutional Review Board of the National Marrow Donor Program. Peripheral blood mononuclear cells (PBMCs) were isolated using Ficoll density gradient centrifugation and cryopreserved according to standard laboratory protocols.

### CONTACT FOR REAGENTS AND RESOURCE SHARING

Further information and requests for resources should be directed to and will be fulfilled by Lead Contact, Ran Reshef.

### METHOD DETAILS

#### Sample summary

The MLR dataset includes 20 patients (**Supplementary Table 2**). For the single-cell RNA/TCR-seq dataset, we analyzed samples from 14 patients and 2 normal donors (NDs). Among these, 11 individuals (patients or NDs) had both upper and lower GI biopsies, resulting in 28 unique biopsies. Of these, 22 biopsies were processed into separate lamina propria and intraepithelial regions, thus totaling 50 samples for RNA/TCR-seq (**Supplementary Table 6**). For the spatial transcriptomics dataset, we analyzed 8 samples from 4 patients and 1 ND (**Supplementary Table 11**). Combining all datasets (MLR, single-cell, and spatial transcriptomics), the study includes a total of 27 unique patients and 2 unique NDs.

#### Cell cultures

CD14+ monocytes were isolated from recipient PBMCs and cultured at 37°C in R10 media (RPMI supplemented with 10% human serum, 0.6% HEPES, 10mg/mL Gentamycin, and 200mmol/L Glutamine) for dendritic cell differentiation.

#### Monocyte-derived dendritic cell generation

The protocol for the maturation of human monocyte-derived dendritic cells was adapted from Rubio et al^86^. Cryopreserved recipient PBMCs were thawed, washed, and resuspended in FACS buffer. CD14+ cells were positively selected using CD14 MicroBeads (Miltenyi Biotec) according to manufacturer’s protocol. Cells were resuspended in R10 media at 1 × 10^6 cells/mL per well in 24-well flat-bottom plates. GM-CSF (50ng/mL) and IL-4 (20ng/mL) were added to each well. Cultures were incubated at 37°C for 9 days. On day 7, TNF-α (25ng/mL) and Prostaglandin E2 (1μg/mL) were added to induce maturation. Cells were harvested on day 9, washed, and resuspended in R10 media at 2 × 10^6 cells/mL.

#### Mixed lymphocyte reaction

The protocol was adapted from Morris et al. with modifications^11^. Cryopreserved donor PBMCs (responders) were thawed, washed, resuspended in 1×PBS at 1 × 10^6 cells/mL and stained with 0.05μM CFSE. After 20 minutes incubation at 37°C, cells were washed with R10 media and resuspended at 2 × 10^6 cells/mL. Cryopreserved recipient PBMCs (stimulators) were thawed, washed, resuspended in 1×PBS at 1 × 10^6 cells/mL, and stained with 0.01μM Far Red. After 20 minutes incubation at 37°C, cells were washed with R10 media, resuspended at 6 × 10^6 cells/mL, and irradiated at 35Gy.

For each MLR well, 200,000 CFSE-stained responders, 600,000 Far Red-stained stimulators, and 40,000 monocyte-derived dendritic cells were combined (5:15:1 ratio) in 96-well round-bottom plates and incubated at 37°C for six days.

#### Flow cytometry and cell sorting

On day 6, MLR cultures were harvested, washed, and resuspended in FACS buffer. Cell surface staining was performed for 30 minutes at 4°C with fluorochrome-conjugated antibodies against CD19, CD14, CD56, CD3, CD4, and CD8. Cells were washed, resuspended in FACS buffer, and stained with cell viability solution. Flow sorting was performed on a BD Influx cell sorter to obtain CD19-CD14-CD56-FarRed-CFSEloCD3+ cells, or both CD19-CD14-CD56-FarRed-CFSEloCD3+CD4+ and CD19-CD14-CD56-FarRed-CFSEloCD3+CD8+ populations, depending on cell quantity.

For donor samples, cryopreserved PBMCs were thawed, washed, and stained with antibodies against CD19, CD14, CD3, CD4, and CD8. The cell viability solution was added after washing. Cells were sorted for CD19-CD14-CD3+ or both CD19-CD14-CD3+CD4+ and CD19-CD14-CD3+CD8+ populations.

#### DNA isolation and TCRB sequencing

Genomic DNA was isolated from post-transplant biopsy specimens using the Qiagen DNeasy Blood & Tissue Kit according to the manufacturer’s protocol and stored at −20°C. DNA from MLR, donor, and post-transplant PBMC samples were isolated using the Qiagen QIAamp DNA Mini Kit according to the manufacturer’s protocol and stored at −20°C. DNA samples were shipped on dry ice to Adaptive Biotechnologies for TCRB sequencing, and data were accessed via the immunoSEQ platform.

#### IEL and LPL isolation

The intraepithelial lymphocyte (IEL) and lamina propria lymphocyte (LPL) isolation protocol was adapted from previously described reports with modifications^13,87^. Biopsies from the lower and upper GI regions were processed separately. Each patient sample (1-4 biopsy pieces) was collected in 1×PBS and kept on ice for less than 15 minutes before processing. Specimens were washed twice in 1×PBS. Tissues were transferred to 50 mL tubes containing IEL buffer #1 (1×HBSS [Ca2+ & Mg2+ free] supplemented with DTT, 0.5 mM EDTA, 0.1mg/mL DNase, 10 mM HEPES, and 5% human serum) and incubated at 37°C for 15 minutes with gentle rotation. Following a 15-second vortex, the suspension was filtered through a 100 μm cell strainer to isolate IELs. This process was repeated once more using fresh IEL buffer #1. A final round was performed using IEL buffer #2 (1×HBSS [Ca2+ & Mg2+ free] supplemented with 0.5 mM EDTA, DNase, and 5% human serum). For LPL isolation, tissues were processed in LPL isolation buffer (RPMI 1640 supplemented with collagenase IV, DNase, HEPES, and gentamycin) using a gentleMACS machine. The resulting suspension was filtered through a 100μm cell strainer with R10 media, washed, and resuspended in FACS buffer.

For single-cell sequencing, CD45+ cells were positively selected using CD45 microbeads when processing time permitted. Otherwise, IEL and LPL fractions were cryopreserved and later thawed for CD45+ cell sorting.

#### 10X scRNA/TCR library preparation

CD45+ cells from IEL and LPL compartments were processed for single-cell RNA and TCR sequencing using the 10X Genomics platform according to the manufacturer’s protocols. 14,000 cells were loaded on a Chromium X using the Chromium Next GEM Single Cell 5’ Kit v2 (10x Genomics, PN-1000263) following the Chromium Next GEM Single Cell 5’ v2 (Dual Index) user guide (10x Genomics, CG000331 Rev E). After reverse transcription and cleanup, cDNA libraries were generated as per the 10x user guide. Construction of final gene expression (GEX) libraries was performed using the Library Construction Kit (10x Genomics, PN-1000190) and Dual Index Kit TT set A (10x Genomics, PN-1000215) according to the user guide. The fragment size distribution of cDNA and final sequencing-ready GEX libraries was assessed using a TapeStation 2200 system (Agilent) with TapeStation D5000 reagents. Matched single-nuclei T cell Receptor libraries were prepared from amplified cDNA libraries using the Chromium Single Cell Human TCR Amplification Kit (10x, PN-1000252).

#### scRNA library sequencing

Final sequencing libraries were quantified using the TapeStation 2200 system (Agilent) with TapeStation D5000 reagents. The concentrations of final GEX libraries were further confirmed using a Qubit fluorometer (Thermo Fisher Scientific). GEX libraries were sequenced on the Illumina NovaSeq X Plus platform with a paired-end 150bp read configuration, targeting ≥ 20,000 read pairs per cell, while TCR libraries were targeted to have >5000 reads per cell. A 2% PhiX spike-in was included to monitor sequencing quality and provide an internal control.

#### 10X VisiumHD spatial transcriptomics

Gastrointestinal biopsies were collected during endoscopies at symptom onset or subsequent time points. Tissues were formalin-fixed, paraffin-embedded (FFPE) and histologically assessed for GVHD by a board-certified pathologist. Clinical GVHD severity was determined using the MAGIC guidelines^25^. FFPE slides (5μm thick) were prepared and assessed for RNA quality using the DV200 assay. Samples with DV200 scores >30% were processed using Visium HD spatial gene expression by the Columbia University Human Immune Monitoring Core. Histomorphological annotation was performed by a board-certified pathologist.

### QUANTIFICATION AND STATISTICAL ANALYSIS

#### T cell receptor data analysis

Twenty patients were analyzed, with 5-12 time points each. Raw adaptive sequencing files were loaded into R with cdr3tools::read_immunoseq() based on functions published in Obradovic et al^88^. The total number of templates ranged from 27,041 to 1,664,944, and the number of MLR templates ranged from 566 to 224,424. A unique clone was defined as a clone with a unique CDR3β amino acid sequence and V and J regions. Template counts were used for the calculation of clonal frequency. The total number of clones from each patient ranges from 4,186 to 377,381 (**Supplementary Table 3**). A clone is considered alloreactive if its frequency is 2-fold greater in the CFSElo fraction compared to the unstimulated donor sample or greater than 1e-5 if not detected in the unstimulated donor sample. In previous studies, Morris et al.^11^ and DeWolf et al.^89^ used a threshold of 1e-5 based on read counts, while Fu et al.^78^. applied a threshold of 2e-5 calculated using template counts. Sensitivity analyses were also conducted here using both 1e-5 and 2e-5, with no major differences observed in the overall analysis results. We identified a total of 51,411 unique alloreactive clones across 20 MLRs, ranging from 256 to 6,243 clones per donor-recipient pair with 1e-5 threshold.

A clone is considered non-alloreactive if detected in either the unstimulated or MLR-CFSElo samples, but did not meet the criteria for alloreactivity. Expanded clones were determined by finding a threshold frequency via the findCutoff function from the KneeArrower package^90^. For each patient, the clones from the MLR-CFSElo sample were arranged in descending order. The loess function from the {stats} package in R was used to smoothen the frequencies, with parameters set to 0.4 and degree to 1. After smoothing, the findCutoff function with the method set to curvature determined the cutoff frequency.

#### Cumulative frequency, Shannon’s Diversity, and clone counts

Three main metrics were calculated for downstream analysis: cumulative frequency, Shannon’s Diversity, and clone counts. The cumulative frequency was calculated as the sum of all clones at a given time point. Both alloreactive and non-alloreactive clones’ cumulative frequencies were calculated. Shannon’s Diversity was implemented via the cdr3tools::repertoire_diversity() function in R. Clone count was defined as the number of unique clones at a given time.

#### TCR data statistical tests

The Wilcox test function from the stats R package was performed for statistical analysis (α = 0.05). Multiple comparison adjustment was performed using the Holm test or Benjamini-Hochberg. Sample binning was performed to understand clonal temporal dynamics. Sample binning was achieved using the quantile function in R.

#### Analysis of temporal dynamics of T cell clones

To further understand T cell clonal temporal dynamics, we adapted a previous basis decomposition method^23^ to find shared and distinct clonal expansion patterns across patients.

##### Input data

Clonotype frequencies were obtained from the Adaptive ImmunoSeq analyzer. To prevent the model from fitting noisy data, the data was filtered to include only clones present in two or more time points. The data was preprocessed with the TCR pipeline. We modeled data from all post-transplant time points through day 180. For each clonotype 𝑖 of a patient, we obtain a vector of clonotype frequencies on each day the patient was measured.

##### The DecompTCR model

The DecompTCR model adapts the model implemented in Nazaret et al.^23^ Theupdated plate model is shown in **Supplementary Fig. 4A**. For clonotype 𝑖, the observed data µ*_i_* of frequencies over time is decomposed as:

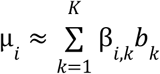

The variables *b_k_* are basis functions of patterns ranging from 0 to 1 which are shared across all clonotypes and β_𝑖,𝑘_ are non-negative weights unique to each clonotype. This decomposition is akin to non-negative matrix factorization^23^. Weights β_𝑖,𝑘_ are independently drawn from a Dirichlet prior with hyperparameter α_0_, and density

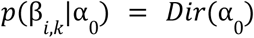

ensuring they are positive and sum to 1.

The basis functions 𝑏_𝑘_ were sampled by drawing neural network parameters (θ, 1D neural network with two hidden layers of 32 units each with tanh activation). For each basis function 𝑘, θ_𝑘_ was sampled from a centered multivariate normal with the diagonal covariance matrix being set to the inverse length scale parameter 𝑙. 𝑙 is a length scale hyperparameter controlling variability—larger 𝑙 allows more complex, flexible basis shapes. The sampled output was then passed through a sigmoid function, ensuring the values ranged from 0 to 1. The overall generative process for each clonotype is interpreted as a Gaussian Process (GP) with the above decomposition serving as the mean function, and a white noise kernel:

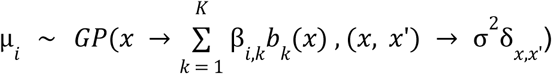

The generative process is as follows:

1. Draw 𝐾 basis functions, 𝑏_𝑘_∼ψ(𝑏_𝑘_) (according to prior described above)
2. For each clone:

a. For each factor dimension 𝑘, draw the basis weight β_𝑖,𝑘_∼ 𝐷𝑖𝑟(α_0_)
b. Draw observed function

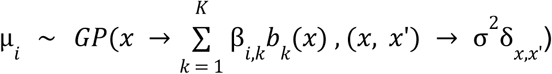

The model uses variational inference to approximate the posterior over the latent variables (θ,β). A mean-field variational family is used, meaning the variational posterior is fully factorized:

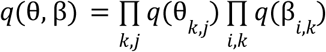

ELBO (evidence lower bound) is then used to approximate the posterior. Since the model cannot handle negative frequency values, the input was shifted by the minimum value to make everything non-negative and normalized to the maximum value in the dataset.

In summary, the key changes compared to the basis decomposition technique previously used for gene expression analysis^23^ included changing to modeling clonotype frequencies, removing the condition plate and including the inverse length scale parameter, which modifies the variability of the basis function with which the model is initialized. Increasing or decreasing the length scale determines how quickly the functions can change. For alloreactive T cells, which tend to have more sporadic dynamics, a larger inverse length scale was required. The conditioning plate was removed due to the need for the exact clone in both conditions, which does not occur in practice. Instead of the standard Gaussian distribution. Alloreactive clones were fitted separately with learning rate=1e-2, beta prior=1.1, inverse length scale=4, weight decay=1e-7, and trained for 5000 iterations. The DecompTCR tool is publicly available on GitHub: https://github.com/azizilab/GVHD_Reproducibility.

##### Evaluation with simulated data

To ensure that DecompTCR could fit real data and discover their true patterns, we first created a simulated dataset with known basis functions. For the simulation, we chose five separate functions: linear increase (𝑦 = 𝑥, 𝑥 ∈ [0, 1]), linear decrease (𝑦 =− 𝑥 + 1, 𝑥 ∈ [0, 1]), sine wave 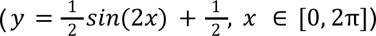, increasing sigmoid (early-rise) 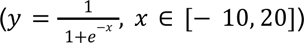, and a later decreasing sigmoid 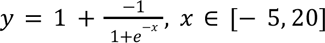. For each function, 181 points were generated within each given range of X. A Dirichlet distribution with a prior of 0.5 was used to sample weights for the summation of each of these functions. Once added together, we downsampled using Bernoulli sampling, weighting the success of accepting a sample higher towards the beginning and decreasing the probability of success linearly for later time points, similar to the collected data. After downsampling, basis decomposition was fitted to determine whether the known basis functions were recovered. R2 scores were used to evaluate the model fit.

##### Interpretation of DecompTCR output

After fitting the model and sampling the beta weights for each clone, hierarchical clustering was used to investigate separation between different grades of GVHD. Some basis functions were repeated by the model; these were combined using hierarchical clustering based on the correlation between basis weights. Using a cutoff of 0.1, three groups of basis functions were identified. We then summed the basis functions and normalized them back to values between 0 and 1. After clustering again on the correlation, no highly correlated basis functions remained. The newly combined clusters were subjected to hierarchical clustering, which preserved separation by GVHD grade.

Values close to zero were filtered out using an approach similar to that used for determining expanded clones. Clones were sorted in descending basis weight order, and the KneeLocator function from the KneeLocator Python package was used to find the elbow point, with parameters S=10, interp_method=“polynomial”, and degree=4. One-sided Mann-Whitney U tests were used to determine whether each basis was greater in one grade than in others. Benjamini-Hochberg correction was applied for multiple comparisons. A basis was considered significant if it was greater than both other grades of GVHD with p-values < 0.05. To accurately assign temporal patterns to GVHD grades, we determined thresholds for each basis by identifying elbow points, excluding clones with low weights. If the proportion of clones from a specific GVHD grade exceeded 33% of the total, that basis was assigned to the corresponding GVHD grade.

##### Hydrophobicity and sequence motif analysis

CDR3β amino acid sequences were grouped based on previously identified GVHD-associated bases. Sequence similarities were analyzed using the BLOSUM62 substitution matrix implemented via the Biostrings^91^ R package. Sequences were preprocessed for quality and converted into AAStringSet objects before calculating pairwise distances with the stringDist function. Mean BLOSUM62 distances were computed for comparisons across three clinical groups: “No to Mild,” “Mild to Severe,” and “No to Severe.” Differences between groups were evaluated using Wilcoxon rank-sum tests, with p-values adjusted using the Holm method.

Shannon’s Diversity Index was calculated for each position in the amino acid sequence using the vegan R package. Positions 5 through 13 showed elevated diversity. The hydrophobicity index was calculated using the Kyte-Doolittle scale for both the entire sequence and specifically for positions 5-13 by taking the mean. Differences between groups were evaluated using Wilcoxon rank-sum tests with p-values adjusted using the Holm method. The lengths of CDR3β sequences were analyzed and compared using Wilcoxon rank-sum tests, with p-values adjusted using the Holm method.

#### Single cell RNA data preprocessing and integration

Raw sequencing data for each sample were processed using the 10x Genomics Cell Ranger pipeline^92^. A total of 50 samples were profiled from 14 patients and 2 healthy donors. Each patient had up to 4 samples per time point, corresponding to the following regions: Upper GI-LP, Upper GI-IE, Lower GI-LP, and Lower GI-IE. Two patients (R160, R066) had samples collected at two different time points. The processed data were loaded into Python using the scanpy.read_10x_mtx() function with var_names=’gene_symbols’ parameter to use gene symbols for variable names. Individual sample datasets were combined into a single AnnData object using anndata.concat() with the parameter join=“outer” to include all genes across samples. The data were normalized using sc.pp.normalize_total() with target_sum=1e4, followed by log transformation using sc.pp.log1p(). Highly variable genes were identified using sc.pp.highly_variable_genes() with parameters n_top_genes=8000, subset=False, layer=“counts”, and batch_key=’PatientID’. Additionally, cell type-specific marker genes were identified from the marker gene table and added to the highly variable gene set, creating a combined gene list for downstream analysis.

##### Doublet detection

Cell doublets were identified using the Scrublet algorithm. For each sample, a Scrublet object was initialized with the raw count matrix (adata.X) and the expected doublet rate specific to each sample. The scrub_doublets() function was called with parameters min_counts=2, min_cells=3, min_gene_variability_pctl=85, and n_prin_comps=30. Doublet scores and predictions were stored in the AnnData object’s observation attributes as ‘doublet_score’ and ‘is_doublet’, respectively.

##### Batch correction using scVI

The AnnData object was set up for scVI^93^ using scvi.model.SCVI.setup_anndata() with batch_key=’PatientID’ and layer=’counts’. A scVI model was created with parameters n_latent=30 and gene_likelihood=’nb’. The model was trained using the train() method with default parameters. The latent representation was extracted using get_latent_representation() and stored in adata.obsm[“X_scVI”]. All patients with samples from both upper and lower GI biopsies were combined and batch corrected.

##### Dimensionality reduction and clustering

Neighbor graph construction was performed on the batch-corrected data using sc.pp.neighbors() with n_neighbors=30 and use_rep=“X_scVI” to use the scVI latent representation. UMAP embedding was generated using sc.tl.umap() with default parameters. Clustering was performed using sc.tl.leiden() with resolution=0.15 to identify major cell populations, and the results were stored in adata.obs[’leiden_0.15’].

##### Secondary doublet detection for T cells

A secondary doublet detection approach was implemented to identify potential doublets within the T cell population that expressed markers of other cell types. The function view_doublets_bycluster() was created to visualize the expression of T cell markers versus markers of other cell types across different clusters. The function doublet_lst() was used to generate boolean indices indicating potential doublets based on co-expression patterns. The following marker gene sets were used:

- T cell markers: CD3E, CD3D, TRAC, TRBC1, TRBC2, TRDC
- B cell markers: CD19, MS4A1
- Myeloid cell markers: CD68, APOE, CD163, CD1C, CD33, CSF1R, MERTK
- Macrophage markers: CD68, MRC1, MSR1, NRP1
- Monocyte markers: CD33, LYZ, FCN1, CSF3R
- Epithelial cell markers: VIL1, CLDN3, OLFM4, LGR5
- Fibroblast markers: ACTA2
- Plasma cell markers: TNFRSF17, SDC1

For each marker gene set, the expression values were averaged across all genes in the set for each cell. Density plots of T cell marker expression versus other lineage marker expression were generated using gaussian_kde. Cells were classified as potential doublets if they expressed markers of non-T cell lineages above a background threshold. Boolean indices for each doublet type were stored in the observation attributes of the T cell AnnData object. A combined doublet indicator (’if_doublet’) was created by identifying cells flagged as doublets in any of the individual doublet detection analyses.

Final T cell analysis was performed using the filtered dataset that excluded all identified doublets, resulting in a refined T cell dataset that was saved for downstream analyses. The total number of T cells after filtering is 38,886. The number of cells for each sample after preprocessing is shown in **Supplementary Table 5**.

##### T cell subset annotation

Dimensionality reduction was performed on the T cell subset by constructing a neighbor graph using scanpy.pp.neighbors() with n_neighbors=30, use_rep=“X_scVI”, and n_pcs=25, followed by UMAP embedding using scanpy.tl.umap(). Leiden clustering was applied specifically to T cells using scanpy.tl.leiden() with resolution=0.5. T cell annotation we performed based on differential gene expression analysis on normalized data with method=’wilcoxon’. Signatures used are listed in **Supplementary Table 5**. Gene expressions were visualized on umap with sc.pl.umap(). For visualization of gene expression across clusters, dot plots were generated using the following parameters: standard normalization, no dendrogram, and viridis colormap.

##### Cross-dataset validation using CellTypist

To validate our T cell subset annotations and assess their generalizability across datasets, we employed CellTypist^94^, a machine learning tool designed for cell type classification in single-cell RNA sequencing data. We use raw counts as input, and the data are normalized (target_sum=1e4) and log transformed. We trained two separate CellTypist models: one for CD4+ T cell subsets and another for CD8+ T cell subsets, using our annotated GVHD dataset as the reference. The model training was performed using celltypist.train() with parameters, n_jobs=10, and feature_selection=True to automatically select discriminative genes.

We applied these models to classify T cells in three independent reference datasets to validate our T cell subset annotations. The first reference dataset was a gut single-cell atlas from healthy adult individuals^37^, which we filtered to include only T cells based on existing annotations. We used cells annotated as “Activated CD4 T,” “Treg,” “SELL+ CD4 T,” and “Th17” for the CD4+ T cell analysis and cells annotated as “Activated CD8 T,” “SELL+ CD8 T,” various γδ T cell subsets, and “CD8 Tem” for the CD8+ T cell analysis. The second and third reference datasets were derived from an inflammatory bowel disease (IBD) study^38^, with T cells annotated using both the Azimuth reference^95^ (CD4: “CD4 TCM,” “CD4 TEM,” “Treg,” “CD4 Naive”; CD8: “CD8 TEM,” “CD8 Naive,” “CD8 TCM”) and the Novershtern panel^96^ (CD4: “CD4+ Effector Memory,” “Naive CD4+ T-cell”; CD8: “CD8+ Effector Memory,” “Naive CD8+ T-cell”). Each reference dataset underwent quality control filtering to remove low-quality cells and was normalized using the same parameters as our GVHD dataset. Cell type prediction was performed using celltypist.annotate() with the parameter majority_voting=False to obtain the most likely cell type for each cell without applying majority voting among neighbors. The results were visualized using celltypist.dotplot() to compare the correspondence between the reference dataset annotations and our model’s predictions.

##### Differential gene expression analysis

Differential gene expression between clusters was performed using the Wilcoxon rank-sum test implemented in scanpy.tl.rank_genes_groups(), use_raw=False, and method=’wilcoxon’. The results were exported using scanpy.get.rank_genes_groups_df() for further analysis. Genes with an adjusted p-value < 0.05 were considered significantly differentially expressed.

#### Cell type proportional analysis across disease severity

To analyze the distribution of T cell clusters across disease severity grades (no GVHD, mild GVHD, severe GVHD), we implemented a computational approach that calculates the proportional representation of each cell cluster within patient samples grouped by disease severity. T cell clusters with fewer than 200 cells were excluded from analysis. Each data point represents a sample (lamina propria or intraepithelial/upper or lower GI). Samples with less than 100 cells were also filtered out to ensure accurate representation of cell type proportion. Statistical comparisons between disease severity groups were performed using the Mann-Whitney U test with Holm-Bonferroni correction for multiple comparisons. Box plots with overlaid strip plots were generated to visualize the distribution of cell type proportions across severity grades, with significance annotations added using the statannotations package. We generated a radar plot visualization approach to visualize the relative distribution of T cell subsets across disease severity grades. The radar plot represents each T cell subset as a distinct axis, with the distance from the center indicating the relative proportion of that subset within each severity group. We calculated the median proportion of each cell 66 type for each disease grade, then normalized these values so that the sum of median proportions equals one within each grade, allowing for direct comparison of T cell subset distribution patterns across disease severity categories.

#### T cell receptor (TCR) repertoire analysis

T cell receptor (TCR) repertoire analysis was performed using scirpy. Filtered contig annotations were processed using the 10x VDJ module. To merge TCR data with the gene expression anndata objects, we applied the ir.pp.merge_with_ir function for each sample individually before concatenating the datasets. Clonotypes were defined using the ir.tl.define_clonotypes function with parameters “receptor_arms=’all’” and “dual_ir=’primary_only’”. For similarity-based analyses, we calculated pairwise distances between TCR sequences using ir.pp.ir_dist with amino acid sequences and alignment-based metrics (cutoff=15). Clonotypes that share similar amino acid sequence were grouped together as one clonotype using ir.tl.define_clonotype_clusters with parameters “sequence=’aa’”, “metric=’alignment’”, “receptor_arms=’all’”, and “dual_ir=’any’”. There were a total of 26,893 cells with clonotype information and a median of 169 clones per sample (**Supplementary Table 5**).

#### Clonotype expansion threshold detection

We implemented an automated approach to identify biologically relevant clonal expansions by determining sample-specific thresholds. For each sample, clonotype frequencies were analyzed using the Kneedle algorithm, which detects the inflection point in the sorted frequency distribution curve. This point represents the natural boundary between expanded and rare clonotypes. A minimum threshold of 2 cells was enforced to exclude singletons. This method accounts for sample-specific clonal diversity, enabling more accurate comparisons of expansion patterns across varying disease severities.

#### Clonal diversity analysis

To assess T cell clonal diversity across disease severity grades, we calculated the Gini index for each sample and T cell cluster. The Gini index quantifies inequality in clonotype frequency distributions, with values ranging from 0 (complete equality where all clonotypes have equal frequency) to 1 (maximum inequality where a single clonotype dominates).

Results were stratified by disease severity (no GVHD, mild GVHD, severe GVHD), and statistical comparisons were performed using the Mann-Whitney U test with Holm-Bonferroni correction for multiple comparisons. Box plots with overlaid strip plots were generated to display the distribution of Gini indices across severity grades, with significance annotations added using the statannotations package.

#### Clonotype-based trajectory analysis

To investigate T cell migration patterns between lamina propria (LP) and intraepithelial (IE) compartments, we developed a T cell receptor (TCR)-based trajectory analysis approach. This method traces shared T cell clonotypes (defined by identical TCR sequences) between tissue compartments to infer migration patterns and phenotypic transitions.

Mobile CD8+ T cells were isolated from the dataset and filtered to include only conventional CD8+ T cell subsets (CD8+ Effector, CD8+ Tissue Resident Memory, and CD8+ Proliferating T cells) and the newly identified CD8+ Hobit+ TRM subset. For each patient sample and disease severity grade (no GVHD, mild GVHD, severe GVHD), we identified T cell clonotypes present in both LP and IE compartments, creating a unique clonotype identifier that combined TCR sequence information with patient metadata.

Migration patterns were quantified by calculating:

- The proportion of each T cell subset in LP and IE compartments
- The proportion of each LP subset transitioning to each IE phenotype
- For visualization of migration patterns, we implemented Sankey diagram plots using Plotly’s go.Sankey function. The Sankey diagrams represented:
- Source nodes: CD8+ T cell phenotypes in the LP
- Target nodes: CD8+ T cell phenotypes in the IE
- Links: Shared clonotypes between compartments, with link widths proportional to the normalized frequency of migration events
- Node colors: Consistent color scheme across diagrams (CD8+ Effector: blue, CD8+ Tissue Resident Memory: green, CD8+ Transitioning Resident: orange, CD8+ Proliferating: pink)

To ensure accurate representation of migration patterns, link values were normalized by compartment size and visualized with semi-transparent connections (opacity 0.3). Statistical analysis of migration patterns across disease severity grades was performed using chi-square tests on the contingency tables of cell type frequencies.

#### T cell trajectory analysis with Decipher

To characterize T cell states and their distribution across patients and conditions, we applied the Decipher method^23^ to the T cell compartment (38,886 cells total from 14 patients and two normal donors). The Decipher algorithm was used to learn a low-dimensional embedding that captures the differentiation of T cells while preserving biological heterogeneity across tissue compartments. We configured Decipher with (decipher_config = dc.tl.DecipherConfig()) and applied it to the T cell raw count data. Training was performed using dc.tl.decipher_train() with plot_every_k_epochs=5 to visualize the embedding during training. After training, the Decipher latent space was rotated to align with CD4/CD8 T cell identities using dc.tl.decipher_rotate_space(). The resulting two-dimensional embedding (stored in adata.obsm[“decipher_v”]) provided a biologically interpretable representation of the T cell states.

We performed clustering on the Decipher latent space using dc.tl.cell_clusters() with leiden_resolution=0.05, n_neighbors=10, and rep_key=‘decipher_z’ to identify distinct T cell states. The clustering results were stored in adata.obs[“decipher_clusters”]. We visualized the T cell states, clinical variables, and gene expression patterns in the Decipher space using dc.pl.decipher() and custom visualization functions. The Decipher embedding revealed distinct clusters characterized by low CD3 expression, a high proportion of mitochondrial reads, and one patient sample (R096) that was enriched for TCRα/β rather than CD45, which was used for enrichment in the other samples. (**Supplementary Fig. 17A-C**). Subsequently, these populations were removed in further analysis of T cell differentiation (13,572 cells left).

In the next analysis, we reapplied Decipher on CD8^+^ conventional T cells after filtering (remove low CD3 expression, and high proportion of mitochondrial reads, as well as patient R096 (13,572 cells left)) to align T cell differentiation trajectories between tissue compartments (Fig. 5E**-G****; Supplementary Fig. 17D).** We limited genes to significant (crit_pval = 0.001, crit_l2fc = 2) differentially expressed genes (DEGs) between the mobile conventional populations (4 clusters including, Hobit+ Trm). The model is trained with the default decipher parameters. The Decipher 1 axis is rotated to align with the Trm to CD8+ effector T cells transition.

#### Clonotype migration flow visualization

We developed DecipherTCR, an extension of the Decipher framework for integrated scTCR/scRNA analysis. Our adapted framework overlays single-cell TCR-seq clonal lineages on the Decipher space obtained from paired scRNA-seq data and enables visualization of T cell migration trajectories between tissue compartments based on shared TCR clonotypes. Our custom function(plot_cell_transitions_stream()) calculates transition vectors between lamina propria and intraepithelial compartments for each clonotype, projecting these vectors onto the 2D Decipher space. After hematopoietic stem cell transplantation (HSCT), immune reconstitution involves tissue-specific T cell migration. Thus, it is reasonable to assume that T cells move from the lamina propria to the intraepithelial region^97^, particularly in the gut—a key immune site and frequent target of graft-versus-host disease (GVHD).

The approach identifies shared clonotypes in both compartments, calculates their mean positions, derives transition vectors, applies spatial weighting to generate a continuous vector field, and creates variable-width streamlines proportional to clone size. Specifically, cell coordinates were extracted from Decipher 2D embedding, and a regular grid was constructed within convex hull boundaries with 2% padding. For each clone *c* present in both compartments, transition vectors were calculated as 𝑣_𝑐_ = µ(𝑋_𝐼𝐸_) − µ(𝑋_𝐿𝑃_), where µ(𝑋_𝐼𝐸_) and µ(𝑋_𝐿𝑃_) represent mean coordinates (2D decipher space) in respective compartments. At each grid point g, unified vector fields were computed using k-nearest neighbors 𝑘 = 𝑚𝑖𝑛(𝑛_𝑐𝑒𝑙𝑙𝑠_/50, 50), with Gaussian weighting 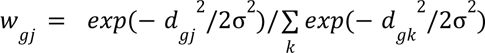, where 𝑛*_cells_* is the total number of cells, σ = smooth × mean_axis_range / n_grid and 𝑑_𝑔𝑗_ is the distance between grid point 𝑔 and cell 𝑗, and 𝑘 is neighbor cell index (50 was selected because 50 neighbors provides a good balance between accuracy and speed), yielding 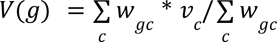 where 𝑤_𝑔𝑐_ is the weighted contribution of clone 𝑐 to grid point 𝑔, and 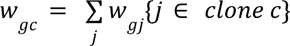. Regions with insufficient cell density were filtered out with density_threshold.

Streamline widths were adaptively scaled by normalized clone size contributions. This approach quantifies directional clone migration patterns while accounting for spatial distribution and clone size heterogeneity. The resulting visualization shows color-coded T cell phenotypes with overlaid streamlines indicating migration patterns, where arrow direction represents LP-to-IE migration and thickness corresponds to clonotype frequency (Fig. 5E**-G**). Parameters were optimized for clarity (n_grid=70, smooth=0.8, density=1.8, arrow_size=2.5, linewidth=2.5, density_threshold=0.1, scale_arrows_by_clone_size=True, min_arrow_width=1.0, max_arrow_width=7.0). The DecipherTCR tool is publicly available on GitHub: https://github.com/azizilab/GVHD_Reproducibility.

#### Spatial transcriptomics analysis

We first attempted to analyze VisiumHD data by clustering 2 μm spots or binning them at 8 μm and 16 μm resolutions, approximating average cell sizes. However, this approach failed to resolve immune cell types and fine-grained cell states (Fig. 6A **middle; Supplementary Fig. 18A left**). We then sought to leverage the histology images to perform cell segmentation and group spots according to cell boundaries (**Supplementary Fig. 19**).

##### Cell segmentation on histology images using CellSAM

Single-cell segmentation was performed on sample SLV14 using CellSAM’s *segment_cellular_image()* function^98^. We first cropped a large patch of the sample and input 2000 by 2000 tiles except for edge tiles to CellSAM segmentation. Masks were stitched back together and visualized on the histology (**Supplementary Fig. 19**).

##### Spatial segmented single-cell transcriptomics analysis

VisiumHD 2 μm binned outputs were allocated to the CellSAM masks and concatenated into one AnnData object with 17584 cells. To do this, we first map each spot to the segmented cell mask, discarding spots that do not fall into any segmented cell, then we sum each segmented cell’s expression across its allocated spots to produce an AnnData object that is single-cell. We preprocess this data by first dropping any cells with no allocated spots, then performing library-size normalization using *scanpy.pp.normalize_total()*^99^ setting *target_sum* to 10,000 and log-transformation using *scanpy.pp.log1p()*. We ran Highly Variable Gene (HVG) analysis using *scanpy.pp.highly_variable_genes(),* selecting the top 2000 HVGs across cells. We then conduct Principal Component Analysis (PCA) using *scanpy.tl.pca()* with 10 principal components on the HVG-filtered log-transformed expression. We used *scanpy.pp.neighbors()* on 25 nearest neighbors and visualized UMAPs using *scanpy.tl.umap()* colored by fibroblast, epithelial and T cell signatures.

This approach also presented its challenges including under-segmentation of specific cell types (e.g. within crypts) and did not resolve immune cells (**Supplementary Fig. 19A,B**) and we found similar challenges with other segmentation approaches^47^. These issues could be related to lateral RNA diffusion across spots and overlapping cells in the 3D tissue architecture^45,46^, e.g. epithelial cells above or below immune cells. To overcome these limitations, we adapted Starfysh^24^, our previous framework for spatial transcriptomic analysis, for application to Visium HD data.

##### Deconvolution of T cell states using StarfyshHD

The StarfyshHD workflow involves three key stages for integrative spatial transcriptomics analysis at high (2µ𝑚) resolution. We analyzed a total of 8 samples (1 sample from ND, 4 from mild, and 3 from severe, **Supplementary Table 11**). First, the data loading and preprocessing stage requires importing spatial transcriptomics count matrices (16µ𝑚 resolution for deconvolution) with their corresponding histology images, along with annotated marker genes for known cell types, followed by creating comprehensive metadata containing patient information, tissue type, and disease grade. The data undergoes quality control including filtering, normalization, and highly variable gene selection to prepare it for integration. Spots with fewer than 50 or 150 UMIs were filtered out depending on sample quality. Spots with more than 20% mitochondrial gene content were also excluded. Additionally, both mitochondrial and ribosomal genes were removed. The top 2,000 highly variable genes were selected for downstream analysis. The median number of spots per sample after filtering is 28,135 (**Supplementary Table 11**).

The second stage involves setting up the Starfysh model parameters and training the model to obtain cell-type proportions per spot throughout joint deconvolution and multiple-sample integration. Gene signatures were generated from scRNA-seq data (**Supplementary Table 7**) and limited to genes exclusive to each cluster to improve deconvolution, which highlights the emphasis of novel T cell state markers. We adopted the default training setup with 100 epochs and 3 restarts. Finally, the downstream analysis phase focuses on examining the learned joint embeddings through UMAP visualization after multi-sample integration, calculating spatial hubs via Phenograph^100^ clustering of spots according to the composition of cell states, characterizing the cellular composition of each identified hub, and quantifying the distributions of these hubs across different samples and disease grades to uncover potential biological patterns. The number of hubs was adjusted to achieve minimum overlaps between cell states.

Cell type proportions were validated using cell type signatures, and visualized with contour plots showing the spatial distribution of this cell type across the tissue (**Supplementary Fig. 18B**). It first selects data for the current sample, then establishes a coordinate grid from the spatial metadata with appropriate padding to avoid visual cropping. The algorithm then calculates the average cell proportion for each grid cell by filtering data points within specific coordinate ranges. To create a smooth visualization, a Gaussian filter is applied to the resulting matrix with a sigma value of 3.0, and the values are normalized to a 0-1 scale. The cell type proportions across different disease severity grades were conducted by constructing normalized frequency matrices of hub cell types across sample categories (fibroblast hub removed). Statistical validation is performed using unpaired t-tests between mild/no and severe disease groups, with results visualized through boxplots incorporating individual data points and significance annotations. ISC hub is defined by any hub that has higher than 30 percent of ISC proportion **(Supplementary Fig. 14)**. The dominating cell type proportion is considered the cell type of the hub. The StarfyshHD tool is publicly available on GitHub: https://github.com/azizilab/GVHD_Reproducibility.

#### Nearest neighbor distance method

We analyze the spatial proximity between cell types across different grades of GVHD through *compute_nndistance(),* a custom function we wrote in the spatial analysis pipeline that calculates the spatial proximity between two specified cell types within a tissue sample. The inputs of the function include the AnnData object *adata*, user-specified *cell_type1* and *cell_type2*, as well as the *sample_id*. Given *adata*, *cell_type1, cell_type2*, and *sample_id*, the function builds cell type masks to extract the spatial coordinates of cells of both cell types for the specified tissue sample. The function then uses *scipy.spatial.cKDTree()*^101^ to build a tree of the *cell_type2* coordinates which represents a spatial index. To query distances and indices of the nearest neighbor in the tree for each point in *cell_type1*, the function uses *scipy.spatial.tree.query()*. The final output of the *compute_nndistance()* is an array of the same shape as the number of cells in *cell_type1* for the specified tissue sample, giving an euclidean distance of each cell to its closest neighbor in *cell_type2*.

#### Crypt loss vs crypt intact analysis

Crypt loss regions were annotated by a board-certified pathologist, and crypt intact region is adjacent to crypt loss region. Cell type proportion differences were compared between crypt loss region and crypt intact region, and statistical significance is assessed using the two sided Mann-Whitney U test to compare the non-parametric distributions between cell types. Hubs enrichment analysis was performed with Fisher’s exact test to identify significantly enriched or depleted hubs in specific tissue conditions by constructing contingency tables comparing hub frequencies between experimental groups. The method calculates counts and proportions for each hub type across different conditions, determines odds ratios to quantify enrichment magnitude, and applies Benjamini-Hochberg false discovery rate correction to adjust p-values for multiple testing. The resulting data frame provides a comprehensive enrichment profile with statistical significance indicators (adjusted p-values < 0.05), allowing identification of spatial hubs that are disproportionately associated with specific experimental conditions.

#### CellphoneDB hub-hub interaction analysis

CellphoneDB^48^ was used to investigate hub-hub interactions in both crypt loss and crypt intact regions of the severe grade sample SLV14. Using CellphoneDB’s *cpdb_statistical_analysis_method()* function and setting 5% expression threshold, we visualized hub-hub interactions, focusing on intestinal epithelial stem hubs’ interactions with T cell hubs. Interaction network plots were produced using *networkx.draw_networkx_nodes()*^102^ and *networkx.draw_networkx_edges()*, with node sizes scaled by hub proportions in the crypt intact and crypt loss regions, respectively to each associated visualization. Edge thickness was scaled by CellphoneDB’s output interaction scores.

